# Revisiting genetic associations with educational outcomes during and after the Soviet era in Estonia

**DOI:** 10.64898/2026.01.06.697950

**Authors:** Ivan A. Kuznetsov, Margherita Malanchini, Oliver Pain, Jonathan Coleman, Philip S. Dale, Jean-Baptiste Pingault, Kathleen Rastle, Estonian Biobank Research Team, Peeter Hõrak, Robert Plomin, Andres Metspalu, Kelli Lehto, Vasili Pankratov, Kaili Rimfeld

**Affiliations:** Institute of Genomics, University of Tartu, Tartu 51010, Estonia; School of Biological and Behavioural Sciences, Queen Mary University of London, London, UK; Social, Genetic and Developmental Psychiatry Centre, King’s College London, London, UK; NIHR Maudsley Biomedical Research Centre, South London and Maudsley NHS Trust, London, UK; Department of Speech and Hearing Sciences, University of New Mexico, Albuquerque, NM, USA; Department of Clinical, Educational and Health Psychology, University College London, London, UK; Department of Psychology, Royal Holloway, University of London, Egham, UK; Department of Zoology, Institute of Ecology and Earth Sciences, University of Tartu, Tartu, Estonia; Bioinformatics Research Centre, Aarhus University, Aarhus, Denmark

**Keywords:** genetic associations, heritability, gene–environment interaction, social transition, educational attainment, occupational status, participation bias

## Abstract

The origins of individual differences in socioeconomic outcomes, including educational attainment and occupational status, reflect a combination of genetic and environmental factors whose relative contributions may differ across historical contexts. Previous research suggested a doubling of the relative role of genetics in explaining variation in these traits in independent Estonia compared with the Soviet era. Using the Estonian Biobank, now tenfold larger than at the time of the original study, we aimed to replicate and extend these findings. We found converging evidence that the relative contribution of genetic factors to educational attainment, but not to occupational status, was higher in the post-Soviet era, although the differences were smaller than previously reported and varied across analytical approaches. However, analyses using narrower birth cohorts revealed a more complex pattern, with the relative genetic contribution to educational attainment decreasing gradually across successive cohorts. Thus, while the Soviet-to-post-Soviet transition was associated with an increase in the relative role of genetic factors in explaining individual differences in educational attainment, it represents only one component of the broader historical changes that have shaped genetic associations with educational attainment in Estonia over the last century.

## Introduction

Socioeconomic status (SES) is linked to many significant life outcomes, including physical and mental health, life satisfaction, happiness, and longevity^1–4^. As with any complex trait, individual differences in SES, whether considered as a composite or through its components (occupational status, educational attainment, wealth, and income), are shaped by environmental and genetic factors. However, the relative contribution of these two, commonly expressed as heritability (the proportion of individual differences attributable to inherited DNA differences), has been a topic of active research and debate. Twin-, family- and population-based studies all converge on both genetic and environmental contributions to individual variation being non-negligible^5–10^.

By definition, heritability of any complex trait is a population-specific parameter that can differ across populations and environments^5,11–13^. Given the importance of environment and gene– environment interactions for SES-related traits, it is natural to hypothesise that heritability of such traits might change with significant environmental shifts, including major social policy changes^5,11^. For example, if as a result of a social transition the variation in educational outcomes due to environmental influences is decreased while the genetic contribution remains constant, that would result in an increase in heritability^11,14,15^. Note, however, that under this model “environment” refers to any source of differences that is not captured by genetics.

We previously showed a massive shift in the aetiology of SES-related traits in Estonia following an abrupt social change following the collapse of the Soviet Union. Genetic differences explained up to twice as much variance in occupational status and educational attainment among individuals who were 15 or younger when Estonia regained independence, that is, those who spent most of their adult life in independent Estonia, compared to those who were older at the time and spent more of their adult life under Soviet rule^16^. These results were interpreted as suggesting that regaining independence led to an increase in meritocratic selection into occupation and education, increasing the importance of genetic differences in explaining individual outcomes in SES compared to the Soviet era, when environmental variation had a larger effect. We also reported suggestive sex differences, with the shift much larger in females than in males, but we were cautious in our conclusions due to limited power in the sex-stratified analyses. Recent evidence from Germany examining cohorts around reunification reported that the genetic contribution to variation in educational attainment was comparable in East and West Germany before reunification, but increased in East Germany afterwards, providing further evidence for changes in heritability for SES outcomes with shifts in historical context^12^.

There are several methods of assessing genetic influence on a phenotype. Approaches that rely on comparing the similarity of identical and non-identical twin pairs result in heritability estimates for socioeconomic outcomes as high as 50%^5,8,17^. Alternatively, heritability can be estimated by quantifying the extent to which phenotypic similarity between individuals is explained by their genetic similarity, as captured by hundreds of thousands of common genetic variants, typically single-nucleotide polymorphisms (SNPs). These SNP-based heritability estimates of SES outcomes average around 20%^7,18–20^. Thus, SNP heritability for educational and occupational outcomes is less than half that of twin-based heritability estimates^21^. This discrepancy likely arises because SNP heritability is estimated from genotyped or imputed common genetic variants and therefore captures only the additive effects of common SNPs represented on genotyping arrays or imputation panel^22^. Alternatively, it has been argued that twin heritability overestimates true heritability^23^, which explains (or rather, explains away) the missing heritability (the gap between family and SNP-based heritability) due to unaccounted confounding factors, assortative mating or violated model assumptions^23^.

Another group of methods for estimating genetic influence on complex traits is based on genome-wide polygenic scores (PGSs). PGSs are constructed from summary statistics of genome-wide association studies (GWAS), which identify genetic variants associated with a phenotype of interest, by aggregating the estimated effects of thousands of SNPs across the genome into a single predictor^24^. These PGSs can serve as predictors of a trait of interest, like socioeconomic outcomes^14^.

One approach is to estimate the proportion of trait variance explained by a PGS, that is, its predictive accuracy. For example, the latest PGS for educational attainment have been shown to explain 12-16% of the trait variance in a sample of European genetic ancestry^25^. Estimates based on PGS predictive accuracy can vary substantially depending on the details of the GWAS and on genetic and environmental similarities between the GWAS cohort and the cohort used to evaluate the PGS^26–30^. Furthermore, these estimates can be downward-biased if effect sizes of individual genetic variants differ between the testing and the GWAS cohorts. As with heritability estimates, the predictive accuracy of PGS may not only reflect direct genetic effects but also factors such as population stratification, assortative mating, and genetic nurture or dynastic effects^31–33^. The latter are particularly relevant for SES-related traits, for which parental genetic influences may shape children‟s environments.

As mentioned earlier, genetics and environment not only contribute independently to trait variation but also can interact with each other: genetic effects on a trait may be stronger in one environment than in the other. Such gene–environment (G×E) interactions can be explored using a variance-component approach, which is generally limited to binary or categorical environmental variables^34^. Alternatively, linear regression can be used to include an interaction term between the PGS and the environmental factor of interest. This approach complements the variance-component analyses by testing whether the mean genetic association captured by a specific PGS differs across environments, while allowing adjustment for interactions between genotype and other environmental factors.

To determine whether cohort differences in estimated genetic associations with SES reflect genuine historical change or measurement and sampling artefacts, this study re-examines the questions raised in the previous report. The Estonian Biobank sample has increased from 20,000 genotyped individuals, who comprised about 2% of the adult population in Estonia, to 210,000 participants, or 20% of the adult population^35,36^. The earlier study had limited power, particularly for sex-stratified analyses, a limitation that is no longer the case. The larger sample also allows us to test whether estimates differ across rural and urban environments, which may have captured different educational and occupational opportunities during and after the Soviet era. Moreover, the recruitment procedure changed after the earlier study, enabling direct tests of participation bias and population-scale generalisability. Methods and discovery resources have also substantially improved. More advanced methods are now available for constructing polygenic scores, and GWAS sample sizes have increased alongside the development of more powerful analytical tools. Together, these advances have greatly improved the predictive accuracy and applicability of polygenic scores.

In this preregistered study (https://osf.io/ug5jp/), we aimed to analyse SNP heritability and polygenic-score prediction for educational attainment, occupational status, height, and BMI across Soviet and post-Soviet birth cohorts; assess whether educational-attainment results hold for both quantitative and binary outcomes; consider the possibility of gradual environmental change; and implement sensitivity analyses that address outcome distributions, participation bias, and recruitment-wave differences. In additional analyses, we examined sex differences, genetic nurture, and whether estimates differed across rural and urban environments. This study aimed to verify whether the previously reported shift in genetic association with socioeconomic outcomes following Estonia‟s independence could be reproduced by re-analysing the same population using updated methods and the same research question. We also examined whether these findings replicate, that is, whether they are observed in a larger sample, remain consistent across traits, are robust to differences in outcome definitions, and are unaffected by sampling and analytical approaches.

## Results

### Data and analysis overview

The Estonian Biobank (EstBB) cohort comprises a volunteer-based sample of the Estonian adult population (aged 18 years or older)^35,36^. The EstBB participants were recruited between 2000 and 2021 through two major data collection waves (Fig. 1a). During the first wave (2000-2016), approximately 52,000 individuals were recruited. A subset of these, genotyped by 2017, formed the sample used in the study by Rimfeld et al.^16^. Approximately 158,000 participants were recruited after 2016. These two periods differ substantially in their recruitment strategies. During the first wave, general practitioners informed their patients about the opportunity to join the EstBB, typically during routine health visits. Additionally, during the pilot period of wave 1, volunteers among university staff and students were actively recruited, likely leading to an overrepresentation of individuals with a university degree in the sample used by Rimfeld et al.^16^ (Supplementary Table 1). In the second wave, participants were primarily recruited through a broad media campaign, including online and television advertisements. Neither wave is entirely representative with respect to educational attainment (EA), with the second wave showing a notable overrepresentation of individuals with tertiary education (Supplementary Table 1).

**Fig. 1.**
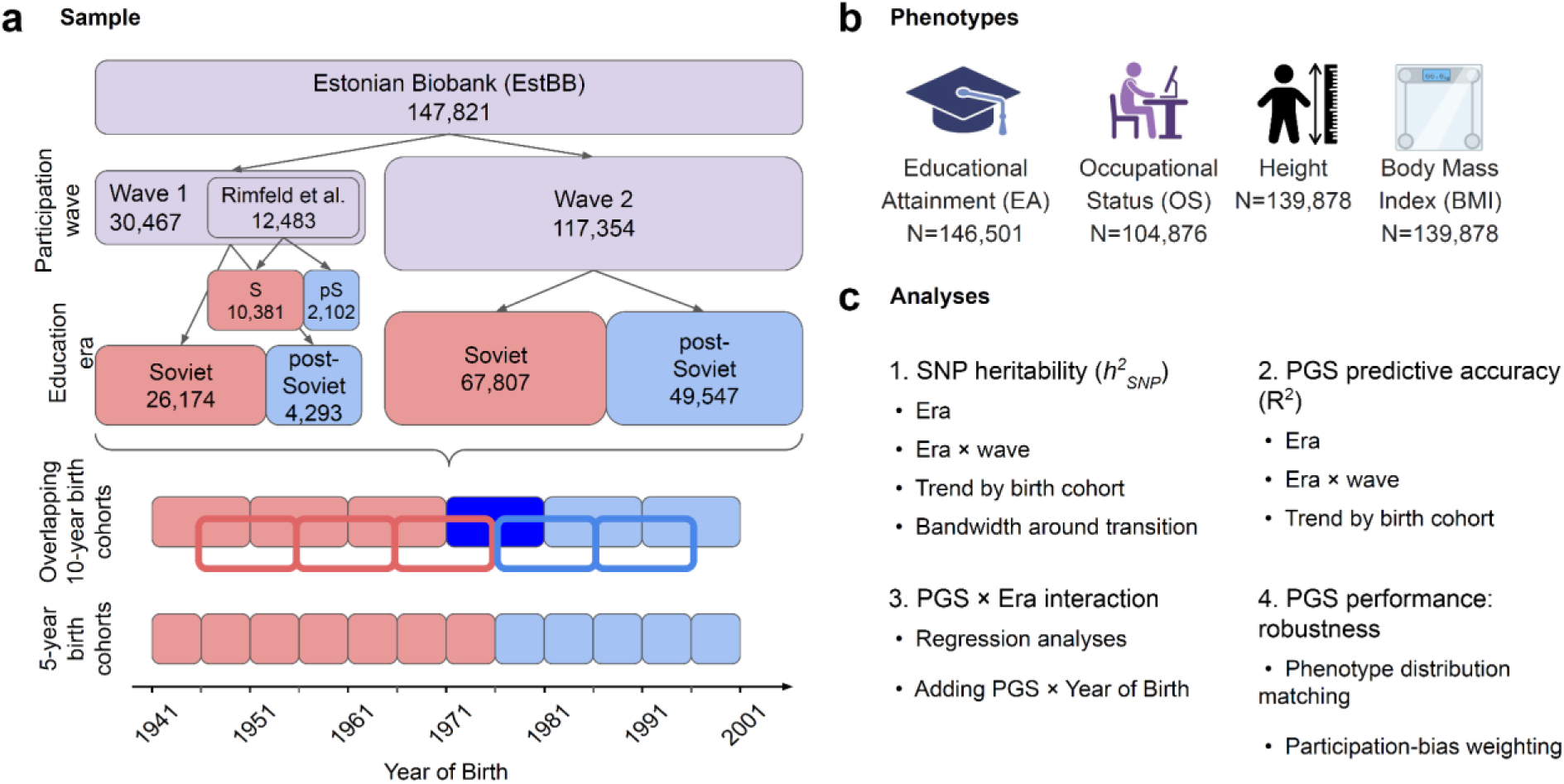
Study overview. (**a**) Data overview showing the composition of the subcohorts used in the analyses. The sample size before filtering related individuals is presented for each subcohort, defined by era and participation wave. The EstBB comprises wave 1 and wave 2, which differ in both recruitment period and strategy. The dataset analysed by Rimfeld et al.^16^ represents a subset of wave 1. In the figure, the Soviet (S, peach) and post-Soviet (pS, light blue) eras were defined using a 15-year cutoff at the time of the Soviet Union‟s collapse. In some analyses, 10- or 5-year birth cohorts were used; adjacent 10-year cohorts overlapped by 5 years. Colours correspond to the era encompassing a given birth cohort, whereas a cohort split evenly across eras is shown in dark blue. (**b**) The phenotypes examined in the study, along with the sample size for each, are reported for individuals before filtering for relatives within the corresponding era and participation wave. (**c**) Overview of the main analyses performed in this study. Numbers indicate the order in which the analyses are presented in the text.

Participants were divided into two birth cohorts, designated as the “Soviet era” and “post-Soviet era,” to maintain consistency with the previous study. We included individuals born before 1976 (older than 15 years old in 1991, when the Soviet Union collapsed and Estonia regained independence) as the “Soviet era”, while younger individuals were included in the “post-Soviet era”. The results with the 15-year cutoff are primarily reported. As a sensitivity check, we repeated our analyses using an alternative grouping method, with 1981 as the birth-year cutoff (i.e., older than 10 years at the time of the collapse of the Soviet Union), again following the Rimfeld et al. approach^16^. The sample sizes of all participants and of unrelated participants (excluding second-degree and closer relatives) in each group used in the analyses are presented in Supplementary Table 2. Descriptive statistics for age, sex, and SES-related variables, including overall sample sizes, are presented in Supplementary Table 3.

In this study, we mainly focus on educational attainment (EA) and occupational status (OS). To put the results for these phenotypes into perspective, we performed the same analyses for two anthropometric traits, height and body mass index (BMI) (Fig. 1b). The sample sizes for the traits range from 139,878 to 146,501 individuals (used in the heritability analysis), and from 87,458 to 115,218 unrelated individuals within the corresponding era and participation wave (used in the PGS analyses and in sensitivity analysis of heritability), when individuals older than 25 at the time of enrolment were included (Supplementary Note 1).

We perform four sets of analyses, with the main analyses presented in Fig. 1c, using all available samples. First, we analyse SNP heritability (*h^2^_SNP_*) in subcohorts divided by era, era and wave, decade of birth and finer year-of-birth bandwidths around the era transition. Second, we compare the polygenic score predictive accuracy, quantified as the coefficient of determination (R²) from linear regression, across subcohorts by era, era and wave, and half-decade of birth. Third, we directly test for PGS×Era interactions using linear regression models with varying specifications. Finally, we analyse the effects of phenotype distribution and participation bias on PGS predictive accuracy.

In addition, we reproduce the original study by Rimfeld et al.^16^ using the original study sample (Supplementary Note 2; Supplementary Figs. 33-35; Supplementary Tables 20, 21), using updated methods. We observe the same trend as in the original study: point estimates of both *h^2^_SNP_* and PGS R^2^ for EA and OS are consistently higher in the post-Soviet era than in the Soviet era across all variations of the analysis, including different phenotype specification, birth year cutoffs defining the Soviet and post-Soviet eras, and methods for estimating heritability. In line with the original study, the differences between eras are statistically significant for most of the PGS R^2^ comparisons, but not for comparisons of *h^2^_SNP_*.

### Preregistered and post hoc analyses

The primary analyses of SNP heritability, PGS predictive accuracy, genetic correlations comparing the Soviet and post-Soviet eras were specified in the preregistered analysis plan (see Methods). We also performed several additional analyses that were not included in the preregistration. These post hoc analyses were undertaken to further investigate the observed differences between eras, assess their robustness to analytical choices and temporal trends, and address issues raised during peer review. They included linear regression with PGS×Era interaction term, analyses using narrower birth-year windows, analyses addressing potential participation bias, analyses using alternative phenotype scaling and heteroscedastic regression, Bayesian analyses of the main comparisons, and G×Era REML analyses.

Unless otherwise stated, analyses presented as part of the primary results were specified in the preregistered analysis plan. Post hoc analyses are explicitly labelled as such when their results are presented below. Preregistered analyses not included in the main text are presented in Supplementary Note 3.

### Analysis of heritability

The results of Rimfeld et al.^16^ suggested that SNP heritability for EA and occupational status (OS) might be higher in the post-Soviet era than in the Soviet era, although these differences were not statistically significant. Before comparing the proportion of trait variance explained by genotype, we estimated the genetic correlations between the eras using bivariate GREML in a non-preregistered analysis. For neither EA nor OS was the genetic correlation significantly different from one, providing no evidence for substantial differences in the genetic factors influencing these traits between the two eras (Supplementary Table 4). Next, we estimated SNP heritability (*h^2^_SNP_*) separately for the Soviet and post-Soviet eras using the restricted maximum likelihood (REML) method implemented in LDAK^21^. To assess the robustness of findings we considered three analytical approaches: (i) using all individuals while modelling genetic similarity with two kinship matrices to maximise sample size and avoid inflation due to shared environment^37^family-inclusive” model, ^37^ and (iii) excluding individuals with second-degree or closer relatives in the sample ^37^. Here and further in the main analysis that follows, EA was measured as years of education (i.e., a continuous measure corresponding to completed years of schooling), and OS on a discrete numeric scale of occupational status categories, with higher values indicating occupations associated with higher socioeconomic status.

Consistent with the original study, EA heritability in the post-Soviet era was significantly higher than in the Soviet era (Δ*h^2^_SNP_*= 0.036, s.e. = 0.010, *p* = 2.7×10^-4^; α = 0.0125), while there was no significant difference in OS heritability (Δ*h^2^_SNP_*= -0.012, s.e. = 0.013, *p* = 0.360; α = 0.0125) (Fig. 2a,b, Supplementary Table 5). The corresponding post hoc Bayesian analysis for EA yielded a posterior mean for *h^2^_SNP_* difference of 0.034 (95% credible interval, [0.016; 0.054]), with a posterior probability of 0.9998 that the difference was positive (Supplementary Table 6). The difference in EA *h^2^_SNP_* remained significant when *h^2^_SNP_*was estimated using the relaxed-threshold model (Δ*h^2^_SNP_*= 0.036, s.e. = 0.012, *p* = 2.7×10^-3^; α = 0.0125) or the stringent-threshold model (Δ*h^2^_SNP_* = 0.035, s.e. = 0.021, *p* = 4.5×10^-3^; α = 0.0125) (Supplementary Figs. 1 and 2, Supplementary Table 5). Using GCTA-GREML to estimate *h*^2^, we received similar results (Δ*h^2^_SNP_*= 0.031, s.e. = 0.012, *p* = 7.5×10^-3^, in the relaxed-threshold model; Δ*h^2^_SNP_* = 0.031, s.e. = 0.019, *p* = 0.092, in the stringent-threshold model; α = 0.0125; Supplementary Fig. 3, Supplementary Table 7). The difference in EA heritability between the Soviet and post-Soviet eras using the alternative 10-year cutoff remained significant only in the family-inclusive model (Δ*h^2^_SNP_* = 0.042, s.e. = 0.013, *p* = 9.4×10^-4^; α = 0.0125) (Supplementary Fig. 4a,c and Supplementary Table 8).

**Fig. 2.**
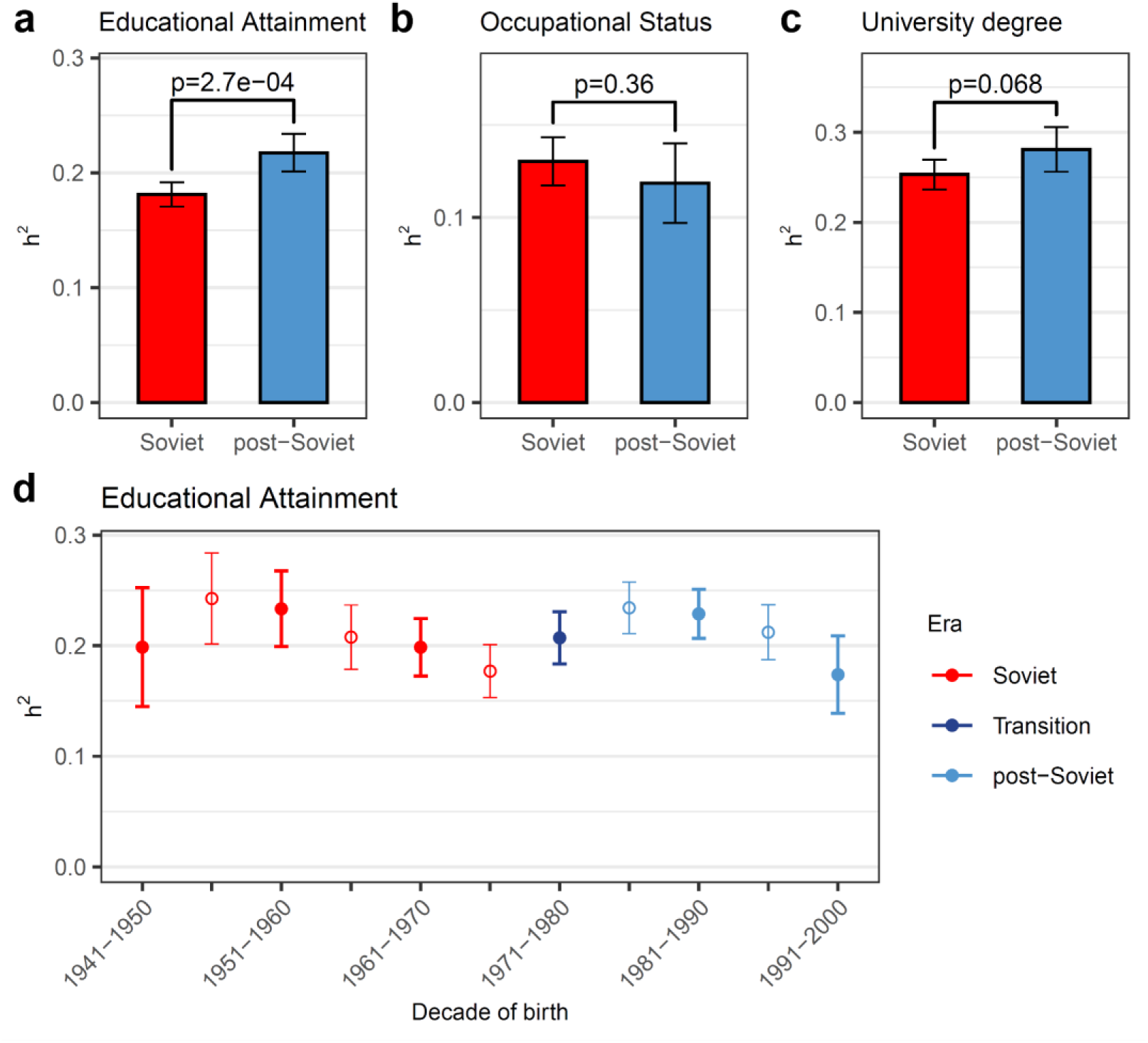
Trait heritability (*h^2^_SNP_*) for educational attainment (EA) and occupational status (OS). (**a**) EA and (**b**) OS heritability is compared across the Soviet and post-Soviet eras on the continuous scale. (**c**) Heritability of EA as a binary trait (University Degree acquired or not), estimated on the liability scale. *P*-values for the difference in h^2^ are shown; the Bonferroni-corrected significance level was α = 0.0125, corresponding to four independent tests. (**d**) Heritability of EA on the continuous scale in decade-long birth subcohorts with 5-year overlap between adjacent subcohorts. Thick error bars and filled circles or thin error bars and empty circles represent non-overlapping subcohorts. Those subcohorts belonging to the Soviet era are coloured in red, and the post-Soviet era in light blue. A transition group comprising individuals from both the Soviet and post-Soviet eras is shaded in dark blue. Error bars correspond to 95% CI.

When stratified by sex, the observed differences between eras were only significant in women in the stringent-threshold model (women: Δ*h^2^_SNP_*= 0.070, s.e. = 0.025, *p* = 0.012; men: Δ*h^2^_SNP_*= 0.024, s.e. = 0.035, *p* = 0.050; α = 0.025; Supplementary Fig. 5, Supplementary Tables 9). Although the estimated difference between eras was larger in women, the difference in Δ*h^2^_SNP_* between women and men was not statistically significant (*p* = 0.28). In the analysis by the type of settlement of residence no significant difference was observed for the 15-year cutoff (Supplementary Fig. 6, Supplementary Table 10).

EA is commonly analysed as a continuous trait^7,25^, and we followed the same approach in the previous analysis. However, despite being transformed into years of education, it retains features characteristic of an ordinal trait. For example, it cannot exceed the value corresponding to a PhD degree. When the mean EA changes, the distribution of EA also changes, affecting the scale on which *h^2^_SNP_* is estimated. Thus, *h^2^_SNP_* on the observed scale may differ between groups even when it does not differ on the liability scale. To address this problem, we conducted a post hoc sensitivity analysis in which we treated EA as a binary trait indicating the presence or absence of a university degree. We estimated its heritability and transformed it to the liability scale (Fig. 2c, Supplementary Fig. 7). No significant difference in *h*^2^ was observed between Soviet and post-Soviet eras when *h^2^_SNP_* was estimated with the family-inclusive model with a 15-year cutoff but the point estimate of *h^2^_SNP_*remained higher in the post-Soviet era than in the Soviet era (Δ*h^2^_SNP_* = 0.028, s.e. = 0.015, *p* = 0.068; α = 0.0125; Supplementary Tables 5 and 8).

Alternatively, to address the scaling issue, we applied a rank-based inverse normal transformation to EA in an additional non-preregistered analysis. First, years of education were adjusted for the covariates, and the resulting residuals were then rank-transformed and mapped to a standard normal distribution separately for men and women. We subsequently estimated SNP heritability of the transformed trait using the stringent-threshold model with the 15-year cutoff (Supplementary Fig. 8). The heritability of inverse normal-transformed EA was significantly higher in the post-Soviet era (Δ*h^2^_SNP_* = 0.067, s.e. = 0.020, *p* = 9.1×10^-4^; α = 0.0125; Supplementary Table 5).

Considering the collapse of the Soviet Union as an event that substantially affected the relative genetic contribution to EA suggests pronounced changes in the temporal trend of EA heritability across the Soviet and post-Soviet eras. To examine whether this historical transition corresponds to a notable shift in the heritability of EA, we estimated EA *h^2^_SNP_*within overlapping decade-long birth subcohorts in a non-preregistered analysis (Fig. 1a). Each bin spanned ten years to ensure sufficient sample sizes for reliable estimates, with a five-year step between bins to provide higher temporal resolution. The EA heritability fluctuates over time, with an overall decreasing trend interrupted by an increase around the focal transition (Fig. 2d). This pattern was reproduced with the binary EA heritability estimates on the liability scale and with inverse normal-transformed EA (Supplementary Fig. 9), and with the estimates from GCTA-GREML (Supplementary Fig. 10). The temporal trend in EA heritability was also consistent across the stringent- and relaxed-threshold models (Supplementary Figs. 11, 12). We also looked at the temporal changes in genetic or environmental variance separately showing that both components change with period of birth for different traits (Supplementary Note 4).

In an additional non-preregistered analysis, to formally assess a potential shift in EA *h^2^_SNP_* around the transition between the Soviet and post-Soviet eras, we estimated differences in EA *h^2^_SNP_* between the two eras across varying birth-year bandwidths centred on the transition year using the stringent-threshold model. We repeated the heritability analyses using symmetric birth-year windows centred on 1976, corresponding to the 15-year cutoff, with bandwidths ranging from 2 to 20 years in 2-year increments. The largest differences in EA *h^2^_SNP_*were observed for bandwidths of 10–14 years around 1976, with the lowest *p*-values obtained for the 10- and 14-year bandwidths (Fig. 3). These results suggest that although the overall trend in EA *h^2^_SNP_* across the full observed birth-year range may be influenced by multiple factors that could obscure differences between eras, the pattern is also consistent with a localized change in EA *h^2^_SNP_* occurring around the transition from the Soviet to the post-Soviet era.

**Fig. 3.**
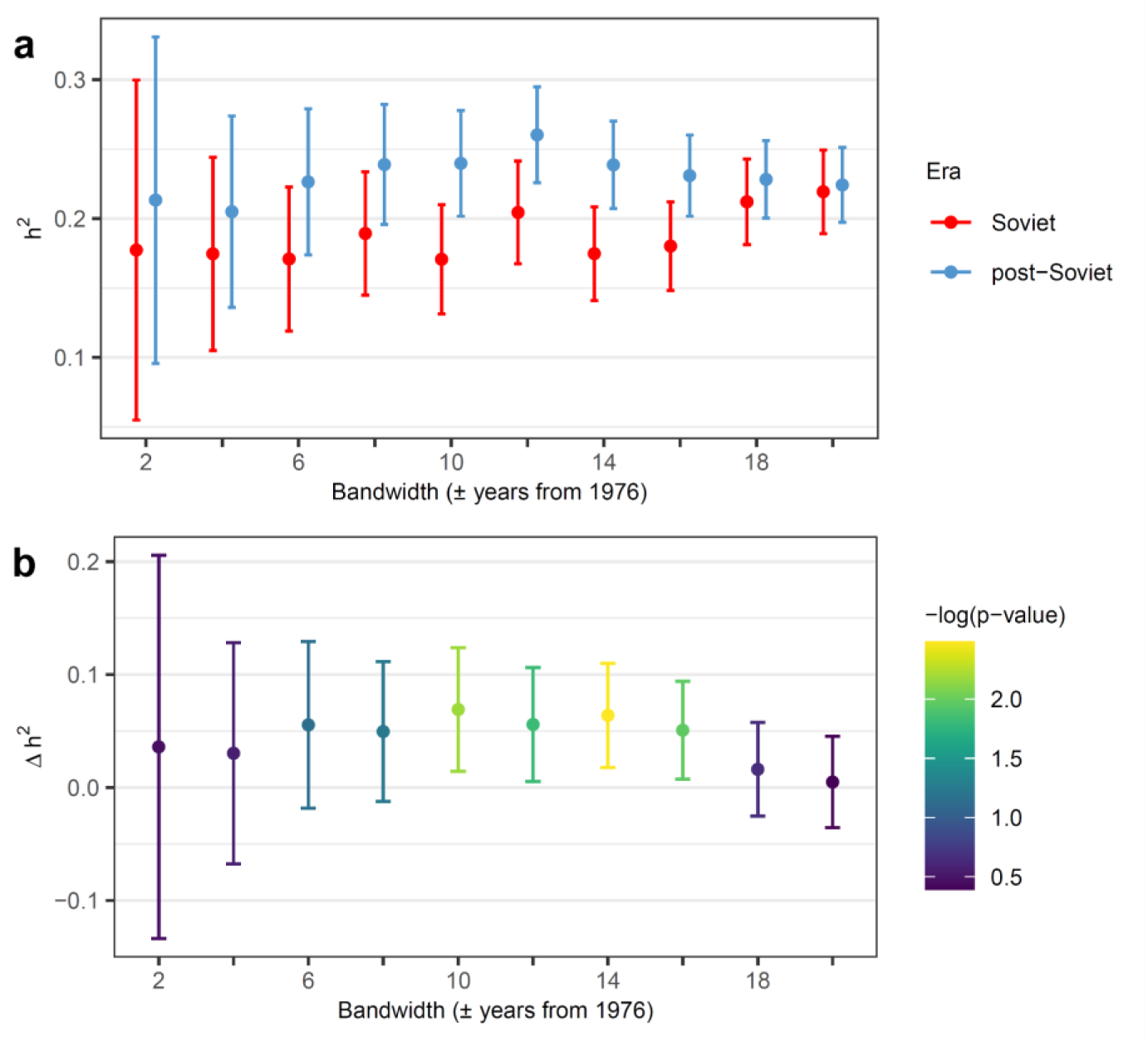
Bandwidth analysis of EA heritability around the transition from the Soviet to the post-Soviet era. Heritability estimates within symmetric birth-year windows around the 1976 boundary (**a**) and the corresponding differences between the post-Soviet and Soviet era estimates (**b**). Bandwidths ranged from 2 to 20 years on each side of the 1976 boundary, in 2-year increments, including individuals born in 1976 to the post-Soviet era. For example, a bandwidth of 10 years corresponds to a comparison between individuals born in 1966-1975 (Soviet era) and those born in 1976-1985 (post-Soviet era). Error bars correspond to 95% CI.

To contextualise these results, we compared the heritability of two anthropometric traits, height and body mass index (BMI), between the Soviet and post-Soviet eras (Fig. 4a,c). Both traits exhibited significantly higher heritability in the post-Soviet era, indicating that such a difference is not specific to educational attainment (Supplementary Table 5). Heritability estimated in decade-long birth subcohorts showed within-era variation (Fig. 4b,d). Moreover, the temporal *h^2^_SNP_* patterns were distinct across the three traits, suggesting that the factors underlying temporal fluctuations in heritability are likely trait-specific. The differences between eras and the temporal trends were consistent with those observed in the stringent- and relaxed-threshold models, although statistical power differed between the models (Supplementary Figs. 11, 12; Supplementary Tables 5, 8).

**Fig. 4.**
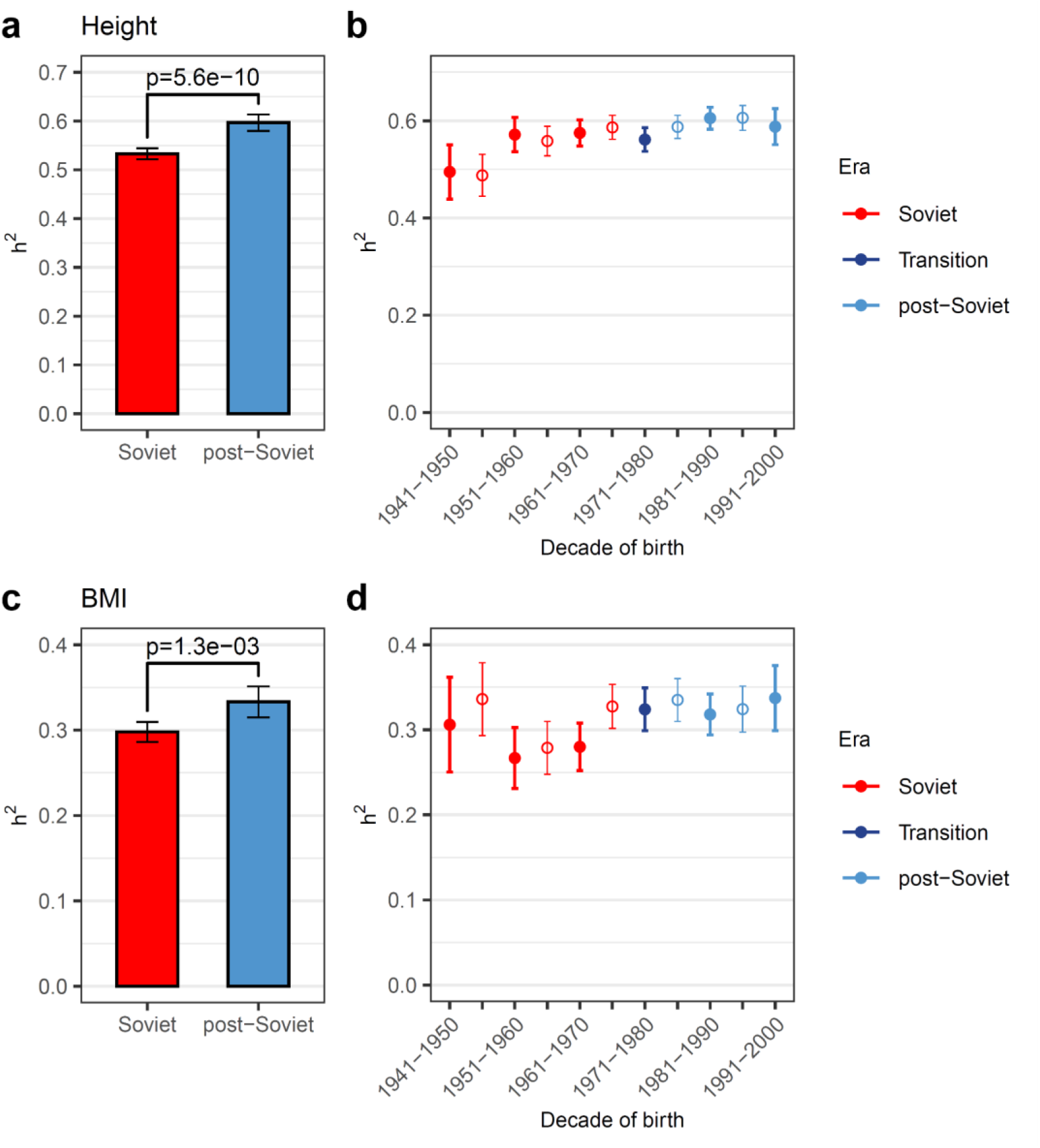
Trait heritability (*h^2^_SNP_*) for height and Body Mass Index (BMI). (**a**) Height and (**c**) BMI heritability are compared across the Soviet and post-Soviet eras on the continuous scale. (**b**) Height and (**d**) BMI heritability in decade-long birth subcohorts with a 5-year overlap between adjacent subcohorts. Thick error bars and filled circles or thin error bars and empty circles represent non-overlapping subcohorts. Those subcohorts belonging to the Soviet era are coloured in red, and the post-Soviet era in light blue. A transition group comprising individuals from both the Soviet and post-Soviet eras is shaded in dark blue. *P*-values for the difference in h^2^ are shown; the Bonferroni-corrected significance level was α = 0.0125, corresponding to four independent tests (panels **a** and **c**). Error bars correspond to 95% CI.

Differential participation bias can affect representativeness of subsamples in different ways, potentially leading to differences in *h^2^_SNP_* estimates between samples without actual differences between the underlying population cohorts^38^. For heritability comparisons across subcohorts to be valid, the results must generalise to the underlying population and not be driven by participation bias. Accordingly, two criteria should hold. First, comparisons between eras within participation waves should not contradict patterns observed in the full sample. Second, no significant differences are expected between participation waves within the same era. In a non-preregistered analysis, we tested whether these criteria hold in the EstBB sample, which comprised two enrolment waves. In practice, significant cross-era differences in *h^2^_SNP_* for height and BMI were observed in wave 2 after Bonferroni correction, with higher heritability in the post-Soviet era, consistent with the main results (Fig. 5, Supplementary Table 5). The large standard errors in wave 1, due to its limited sample size, make it unlikely that significant *h^2^_SNP_* differences can be detected. The comparison of *h^2^_SNP_* estimates between waves within the same era yielded a single significant result: height heritability in the Soviet era differed significantly between wave 1 and wave 2 (*p* = 3.3×10^-5^; α = 0.00625). Thus, when it comes to EA, we do not see any differences in the estimates for wave 1 and wave 2 whether on the scale of years of education, university degree, or inverse normal-transformed EA. Still, significant differences in height *h^2^_SNP_* between wave 1 and wave 2 suggest that the two waves cannot be considered two random samples from the same underlying population, which impairs generalisability. The violation of the second criterion for height is likely explained by the coincidence of an increasing heritability trend across birth years and the earlier mean year of birth among participants in wave 1 (Supplementary Table 3).

**Fig. 5.**
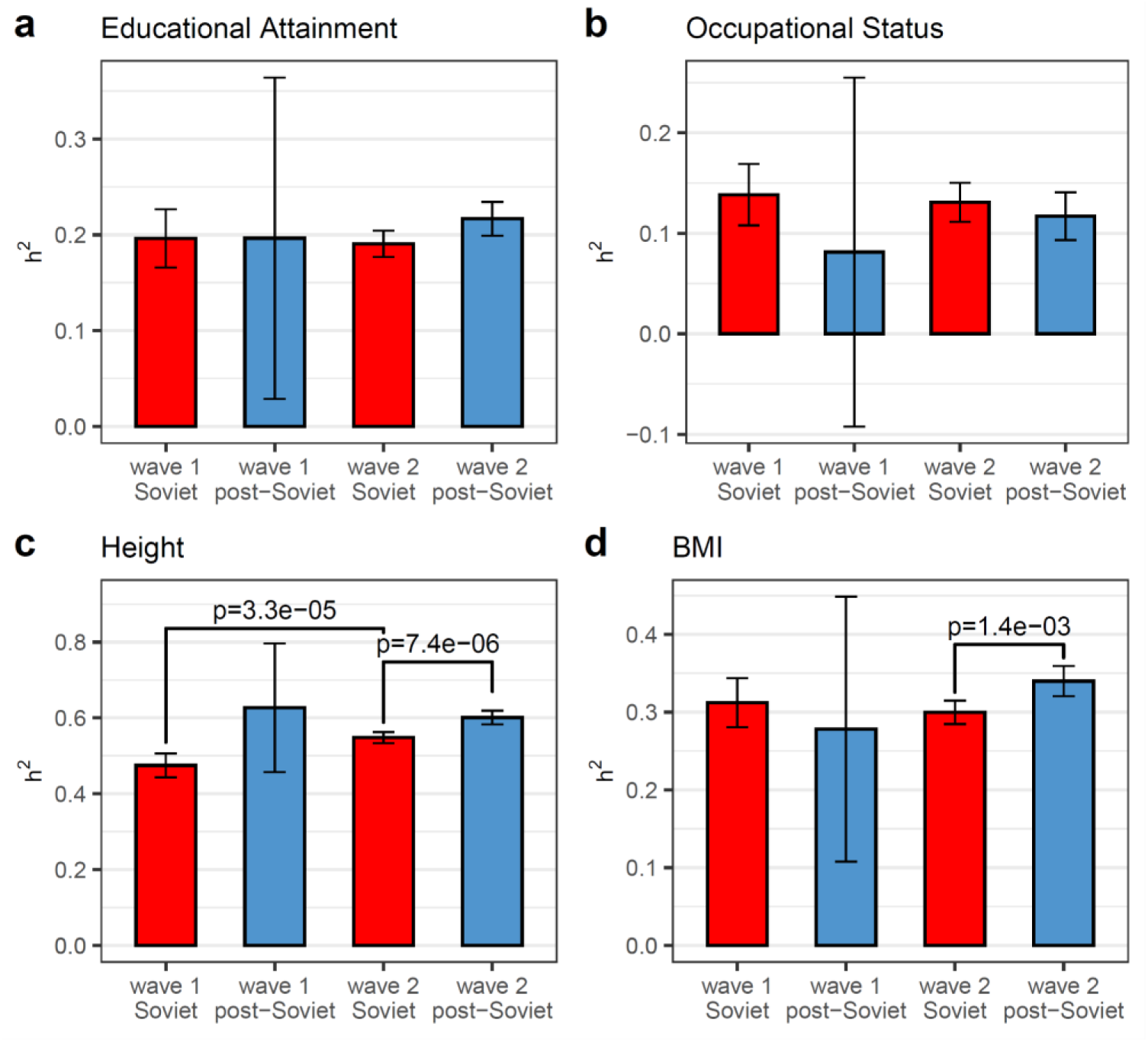
Trait heritability in the Soviet and post-Soviet eras by participation wave. Trait heritability (*h^2^_SNP_*) is shown in groups divided by era of basic school education (left panels) and wave of biobank enrolment for (**a**) educational attainment (EA), (**b**) occupational status (OS), (**c**) height, and (**d**) body mass index (BMI). Those subcohorts belonging to the Soviet era are coloured in red, and the post-Soviet era in light blue. Pairwise comparisons of *h^2^_SNP_* were conducted either between waves within each era or between eras within each wave. The Bonferroni-corrected significance level was α = 6.25×10^-3^, corresponding to eight independent tests. Only *p*-values below the corresponding significance threshold are shown. Error bars correspond to 95% CI.

### Predictive accuracy of polygenic scores

Rimfeld et al.^16^ reported that the polygenic score for educational attainment (PGS_EA_) explains more variance in educational attainment and occupational status in the post-Soviet era than in the Soviet era. In this study, we evaluated and compared across eras the predictive power of a polygenic score from the PGI repository^39^ constructed from the most recent and well-powered EA GWAS summary statistics (EA4^25^), with Estonian Biobank cohort excluded from the meta-analysis. For the sensitivity analysis, we used GWAS summary statistics from the meta-analysis with the 23andMe cohort additionally excluded. The SBayesR method^40^ was used to re-estimate SNP effects while accounting for linkage disequilibrium (LD) structure.

Differences in the genetic architecture of a trait between the discovery population used for GWAS-based PGS construction and the target population in which the PGS is applied can reduce predictive accuracy. Therefore, we first verified that the genetic architecture of EA in the GWAS sample was compatible with that in the EstBB. The genetic correlation between EA in the EA4 meta-analysis sample (excluding the EstBB and 23andMe cohorts, because we did not have access to the raw summary statistics with the 23andMe cohort included) and in the EstBB, estimated using LD score regression, was not significantly different from one (r_g_ = 0.98, s.e. = 0.03, *p* = 0.53). This high genetic correlation indicates that EA in the EstBB shares a similar genetic architecture with the EA4 sample, supporting the validity of applying EA4-based polygenic scores in the Estonian population.

We used PGS_EA_ as a predictor of educational attainment and occupational status in two models. In the first model, we estimated the incremental R^2^, defined as the proportion of variance in the trait explained by the PGS in addition to that explained by covariates (sex, age, sex×age, age^2^, and 40 genetic principal components). In the second model used for sensitivity analysis, the traits were pre-adjusted for these covariates by regressing them out, and R^2^ was then calculated from a univariate linear regression with the PGS as the sole predictor. First, we tested the predictive accuracy of PGS_EA_ in the entire EstBB cohort after excluding second-degree and closer relatives. The incremental R^2^ was 0.109 and 0.076 for the PGS_EA_ based on the EA4 meta-analysis including and excluding the 23andMe cohort, respectively.

We then compared the R^2^ estimates obtained for the Soviet and post-Soviet eras, both without and with splitting by participation wave. For EA, both incremental R^2^ and R^2^ for the pre-adjusted trait were significantly higher in the post-Soviet than in the Soviet era (ΔR^2^ = 0.011, p < 0.002; α = 0.0125 in both models) when the full sample was used (Fig. 6a; Supplementary Fig. 13; Supplementary Table 11). The corresponding post hoc Bayesian analysis yielded a posterior mean for incremental R^2^ difference of 0.011 (95% credible interval, [-0.008; 0.030]), with a posterior probability of 0.870 that the difference was positive (Supplementary Table 6). The era differences in R^2^ were not significant for inverse normal-transformed EA or when the PGS calculated from GWASs that excluded the 23andMe cohort was used (not preregistered; Supplementary Figs. 14-16; Supplementary Tables 11, 12).

**Fig. 6.**
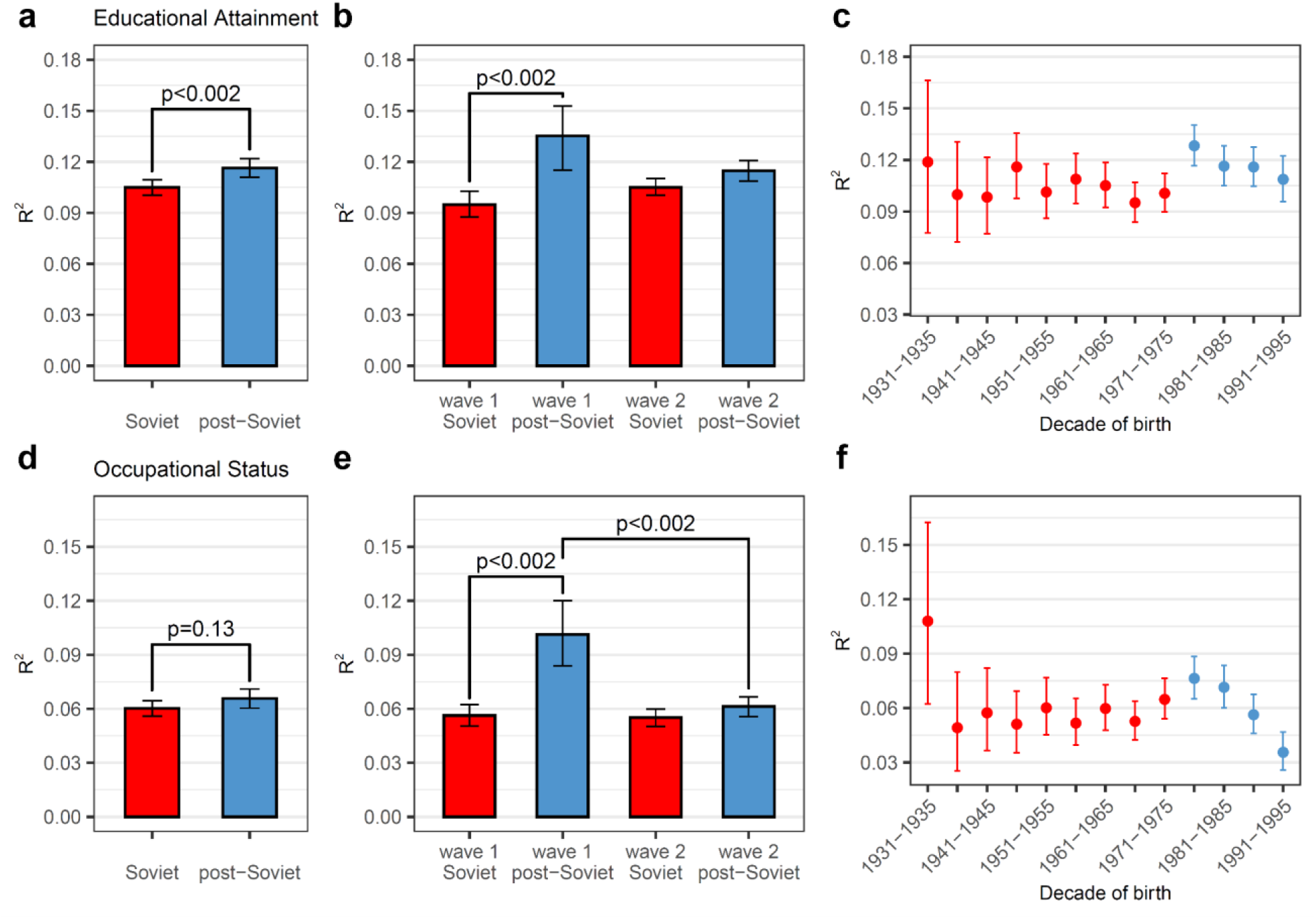
Trait variance explained by PGS (incremental. **R**^2^**) in the Soviet and post-Soviet eras.** R^2^ is shown in groups divided by era (in the left panels), by era and wave of biobank enrolment (in the middle panels), or by half-decade of birth (in the right panels) for educational attainment (**a, b** and **c**, respectively), and occupational status (**d, e** and **f**, respectively). Pairwise comparisons of R^2^ were conducted between eras (**a**, **d**), and between waves within each era or between eras within each wave (**b**, **e**). Those subcohorts belonging to the Soviet era are coloured in red, and the post-Soviet era in light blue. *P*-values for the difference in R^2^ are shown (panels **a**, **b**, **d**, **e**). The Bonferroni-corrected significance level was α = 0.0125 (panels **a** and **d**), corresponding to four independent tests, and α = 6.25×10^-3^ (panels **b** and **e**), corresponding to eight independent tests. Only *p*-values below the corresponding significance threshold are shown on panels **b** and **e**. Error bars correspond to 95% CI.

For OS, R^2^ values did not differ significantly between the eras (ΔR^2^ = 0.002, *p* = 0.13; α = 0.0125) (Fig. 6d; Supplementary Figs. 13, 15, 16; Supplementary Tables 11, 12). No significant differences in R^2^ between the eras were observed for EA or OS when the 10-year cutoff was used (Supplementary Figs. 17, 18; Supplementary Table 13). When the comparison was performed separately by enrolment wave (not preregistered), significant differences between eras were observed for EA in wave 1 (p < 0.002 for both incremental R^2^ and in R^2^ for the pre-adjusted trait) (Fig. 6b and Supplementary Table 11). In the sex-stratified analysis, the difference between eras was significant for women, with higher R^2^ in the post-Soviet era, but not for men (ΔR^2^ = 0.010, *p* = 0.026; ΔR^2^ = 0.011, *p* = 0.002, respectively; α = 0.0125). However, this difference is likely to reflect differences in statistical power, as the R^2^ estimates were very similar between the two sexes (Supplementary Fig. 19; Supplementary Table 14). We also found that the difference between eras was significant among individuals residing in rural areas but not among those living in towns or cities (Supplementary Fig. 20; Supplementary Table 15). For OS, R^2^ was significantly higher in the post-Soviet era than in the Soviet era in wave 1, but not in wave 2 (Fig. 6e; Supplementary Tables 11, 12).

As with heritability analysis, a comparison of PGS R^2^ is valid and generalisable only if there are no significant differences between participation waves within the same era. In a non-preregistered analysis we found that the PGS R^2^ did not fully meet this criterion. Wave 1 and wave 2 differed significantly in incremental R^2^ for OS, and in R^2^ for the pre-adjusted trait for both EA and OS (Fig. 6b,e; Supplementary Table 11). We note that R^2^ strongly depends on the distributions of the trait value and PGS and is especially prone to the effect of participation bias which cannot be easily corrected for (see “Stability of polygenic score predictive power” section). Therefore, we cannot disentangle differences in the predictive accuracy of PGS due to sample-related characteristics and historical context. Consequently, comparisons of PGS predictive accuracy between the Soviet and post-Soviet eras should be interpreted with caution.

In a non-preregistered analysis, to examine how the variance explained by PGS changed over time, we divided the sample into half-decade birth subcohorts and calculated incremental R^2^ for each subcohort. For EA and OS, R^2^ in the earliest post-Soviet birth cohort demonstrates an upward shift relative to the latest Soviet cohort (Fig. 6c,f). This pattern was replicated using PGS_EA_ based on summary statistics that excluded the 23andMe cohort from the meta-analysis and using inverse normal-transformed EA (Supplementary Figs. 14, 21). However, this increase was to a large extent driven by participants from wave 1, and was followed by a decline in R^2^ in later cohorts (Supplementary Fig. 22). Results for height and BMI at both the era and year-of-birth levels are presented in Supplementary Figs. 13, 15, 16, 21-24 and Supplementary Tables 11 and 12.

### Variation in polygenic score effects across eras

In a non-preregistered analysis, we explicitly tested the G×Era interaction for EA within the REML framework using the combined sample with the stringent relatedness threshold and found no evidence of a significant interaction (Supplementary Table 16).

Next, we investigated the linear interaction of the PGS_EA_ with the SES outcomes. This analysis was not included in the preregistered study plan. We first applied linear regression models with EA (measured in years of education) and OS as dependent variables in a subsample of unrelated individuals (no second-degree or closer relatives). The models included wave, era, year of birth (YoB), sex, and all pairwise interactions, including squared YoB, as covariates. Following standard recommendations^41^, we included PGS_EA_ and interaction terms between PGS_EA_ and wave, YoB, and sex, in addition to the primary PGS×Era interaction. Analyses were also stratified by participation wave (Fig. 7a,c).

**Fig. 7.**
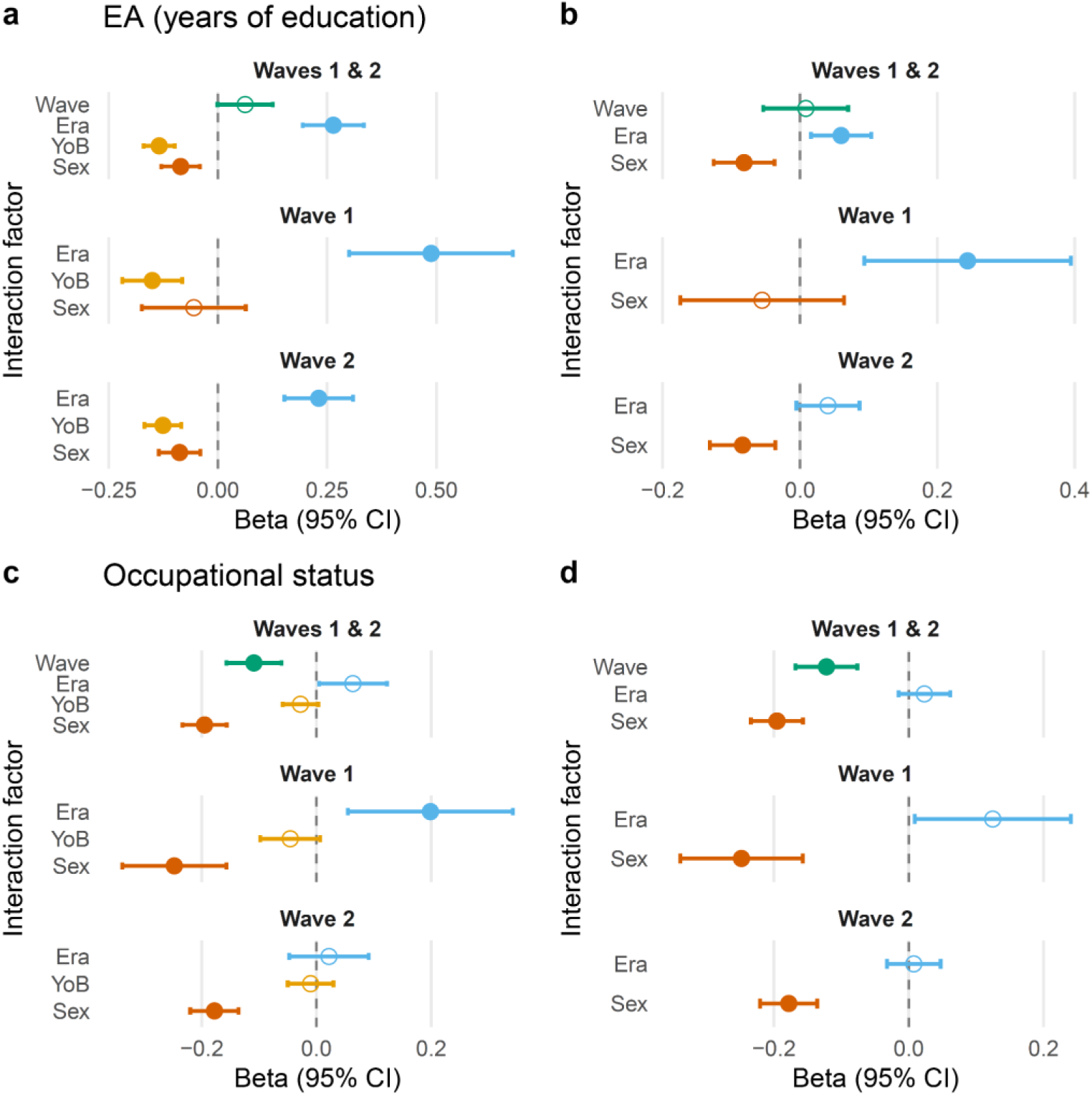
PGS interaction effects with era and demographic factors across outcomes. Linear ordinary least squares regression models for EA (years of education) (**a**–**b**) and occupational status (**c**–**d**) show estimated interaction effects (β, 95% CI) between PGS_EA_ and era (0 = Soviet; 1 = post-Soviet), wave (0 = wave 1; 1 = wave 2), sex (0 = male; 1 = female), and year of birth (YoB, z-standardised). PGS_EA_ was obtained from the PGI repository. Models in the left column include the PGS×YoB interaction; models in the right column exclude it. Points represent estimated interaction effects (β) with 95% confidence intervals. Filled circles denote statistically significant effects. The Bonferroni-corrected significance level was α = 0.0125, corresponding to four independent tests, in analyses of the combined wave 1 and wave 2 samples (Waves 1 & 2) and α = 6.25×10^-3^, corresponding to eight independent tests, in analyses stratified by wave.

The PGS×Era interaction was significant in the full sample and in both wave-specific subsamples, indicating a stronger PGS_EA_ effect in the post-Soviet era (β= 0.264, *p* = 1.4×10^-13;^ α = 0.0125). The corresponding Bayesian analysis yielded a posterior probability of > 0.999 that the coefficient for PGS×Era was positive (Supplementary Table 6). For OS, the interaction was not significant in the full sample. No significant PGS×Era interactions were detected for height or BMI (Supplementary Fig. 25).

Because the era was defined by year of birth, the post-Soviet era, coded as 1, corresponds to higher YoB values. Remarkably, the regression results showed that PGS×Era and PGS×YoB effects were in opposite directions, suggesting a non-linear relationship between PGS_EA_ effects and birth cohort. To examine this, we repeated the analysis excluding the PGS×YoB term (Fig. 7b,d). This led to an attenuation of PGS×Era effects in all subsamples. These results indicate that PGS_EA_ effects vary continuously across birth cohorts within each era, declining as birth year increases. When this trend is controlled for (i.e., when PGS×YoB is included), PGS×Era interactions are more pronounced, especially for EA. However, when comparing the average PGS_EA_ effects across the Soviet and post-Soviet eras, rather than modelling birth-year-related trends (i.e., excluding the PGS×YoB term), the differences in the PGS_EA_ effects are modest and participation-wave specific.

As a sensitivity analysis, we conducted a heteroscedastic regression in the full sample of unrelated individuals for inverse normal-transformed EA, in which trait variance, not only its mean, was modelled as associated with PGS_EA_. Supporting the main result, the effect of PGS×Era was positive when PGS×YoB was included and indistinguishable from zero without PGS×YoB (Supplementary Fig. 26).

Because EA and OS are ordinal rather than continuous traits, treating them as continuous in linear regression can bias interaction estimates^42^, and the inverse normal transformation does not necessarily adequately represent the underlying scale. To assess robustness from another angle, we transformed EA and OS into binary outcomes and applied logistic regression. EA was coded as the presence or absence of a university degree, and OS distinguished managers, professionals, technicians and associate professionals from all other categories, yielding two approximately equal groups. In this framework, PGS×Era remained a significant predictor of EA (Supplementary Fig. 27). For OS, PGS×Era was significant in the full sample when PGS×YoB was excluded from the model.

When using the year of birth 1981 (10 years in 1991) as the cutoff between Soviet and post-Soviet eras, PGS×Era coefficients were substantially attenuated in all the tested models (Supplementary Figs. 28 and 29), indicating that the 15-year cutoff better reflects the potential transition effects and emphasising that the results depend on specific cohort definitions and sample composition. The results with PGS_EA_ calculated using summary statistics without the 23andMe cohort involved (15-year cutoff) were in line with main results, although the interaction estimates were generally lower, consistently with lower predictive power of this PGS (Supplementary Figs. 30 and 31).

### Stability of polygenic score predictive accuracy

Despite the larger sample size, we were able to replicate the difference in EA R^2^ between the Soviet and post-Soviet eras reported by Rimfeld et al. using only the most powerful PGS, the untransformed trait, and the 15-year era cutoff. At the same time, we observed that R^2^ can vary considerably even between cohorts from the same era when recruitment times and procedures differ. One plausible explanation is that R^2^ is affected by the distribution of the dependent variable^43^. Indeed, the cohorts, whether grouped by participation wave or by era, show distinct EA distributions (Supplementary Table 1).

To examine how such distributional differences influence R^2^, we performed a post hoc resampling analysis. For each of the four groups defined by era (Soviet, post-Soviet) and participation wave (wave 1, wave 2), we repeatedly drew random subsamples with replacement from that group (“source group”). Individuals were sampled in such a way that the EA distribution in each subsample matched that of another group (“target group”) (Fig. 8). This procedure was also repeated while matching jointly on EA and sex (Supplementary Fig. 32).

**Fig. 8.**
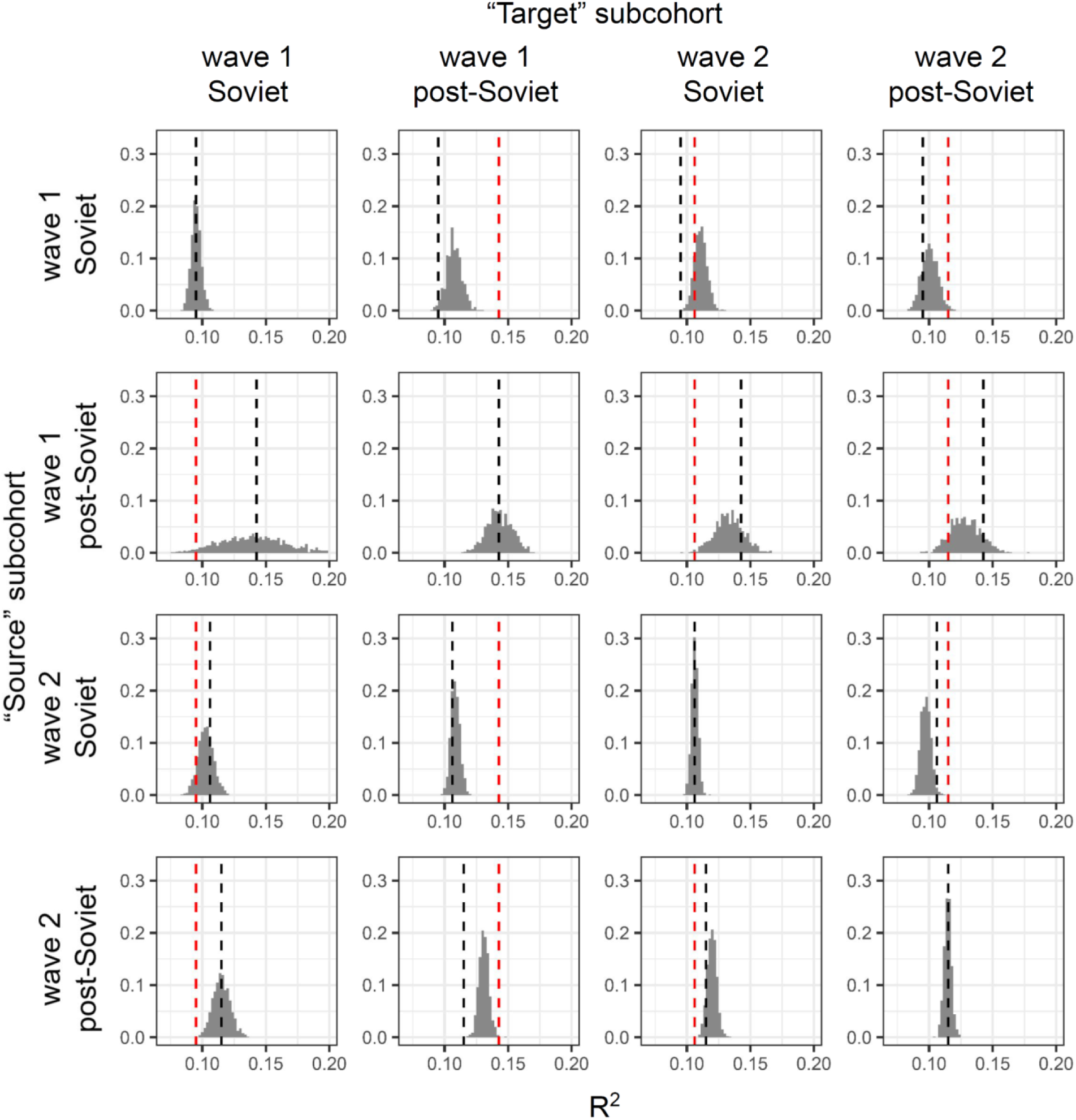
Distribution of EA variance explained (R^2^) by PGS_EA_ in subsamples generated to match the EA distribution of a target subcohort. Subsamples were drawn with replacement from a “source” subcohort defined by recruitment wave and era, matching the EA distribution. Rows indicate the source subcohort from which individuals were sampled, columns indicate the target subcohort whose EA distribution was used for matching. For each source–target pair, 1,000 resampled subsamples were generated, and R^2^ was calculated as the squared Pearson correlation between the pre-adjusted EA residuals (adjusted for sex, age, sex×age, age^2^, and 40 PCs) and PGS_EA_. Histograms show the distribution of R^2^ across resamples. Black lines indicate the original R^2^ in the full source subcohort, red lines indicate the R^2^ in the target subcohort. Diagonal panels represent cases where source and target groups are identical. PGS_EA_ was obtained from the PGI repository.

We then recalculated R^2^ for each resampled dataset. In most cases, the resulting R^2^ values shifted from the source group’s original R^2^ toward the target group’s R^2^, with the mode of the resampled R^2^ distribution lying between them. The extent of the shift ranged from negligible to nearly complete. In some cases, the distribution modes lie outside the range between the original R^2^ values of the source and target groups. These results demonstrate that R^2^ is sensitive to differences in the trait distribution, which may arise either from participation bias or from genuine population differences.

As shown in Fig. 8, PGS R^2^ differs between some cohorts from the same era but different recruitment waves. This may reflect, among other factors related to recruitment strategies, differences in the distribution of the phenotype itself. These distributional differences likely arise because individuals with different characteristics, particularly educational attainment, were more likely to participate in different waves. In an additional non-preregistered analysis, to test whether adjusting for participation bias could reduce these differences, we applied weights to the EstBB sample using census data from the period of data collection, accounting for age, sex, and EA. In the weighted subsamples, the distribution of EA across eras and waves was matched to the census distribution within each sex and 5-year age group. As an alternative approach, the joint distributions of EA, age and sex were matched to those in the census.

Comparing R^2^ across subcohorts (Fig. 9 and Supplementary Table 17) revealed that weighting had only a modest effect. Although the confidence intervals widened due to the reduced effective sample size after weighting, waves 1 and 2 still showed a significant difference in R^2^ for OS within the post-Soviet era and for height within the Soviet era. These findings indicate that participation bias related to age, sex, or educational attainment cannot fully explain the observed differences in PGS R^2^. This implies that additional, unobserved factors contribute to the variability in R^2^, limiting its utility for comparing genetic effects across time-related cohorts. Indeed, other wave-specific factors (e.g., health problems, since wave 1 recruitment was mostly conducted by general practitioners) may also play a role, and our current adjustment for participation bias is not exhaustive. Within this framework, the distribution of year of birth was not equalised between the waves, and between-wave differences could therefore reflect variation in PGS effects across birth years within each era.

**Fig. 9.**
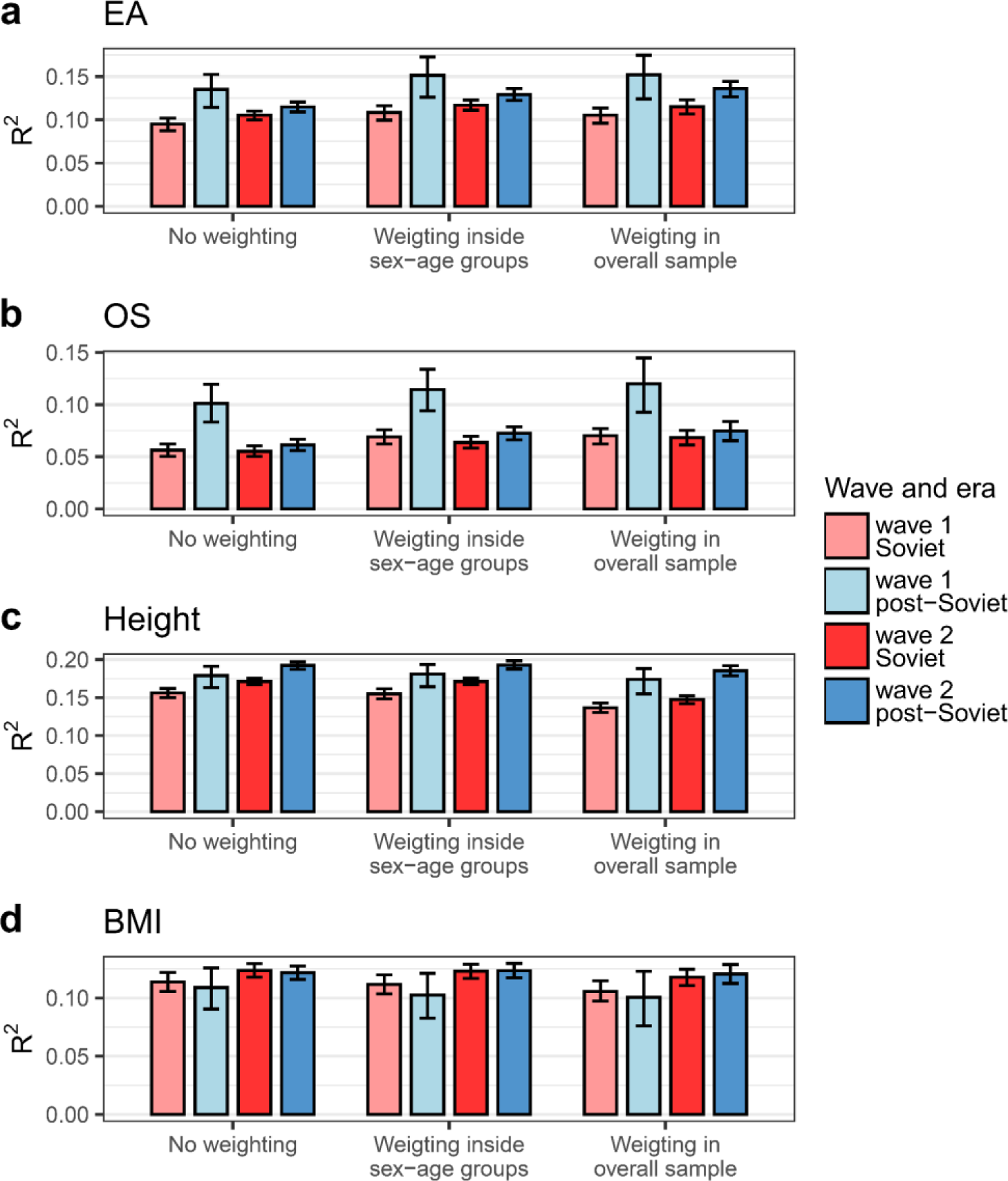
Trait variance explained by PGS in weighted subsamples. Subsamples are based on era and wave of biobank enrolment. In all cases, incremental R^2^ is shown. Results of three weighting procedures are shown: no weighting; based on the EA distribution in each sex-age group (“inside sex-age groups”); based on the joint distribution of sex, age, and EA (“in the overall sample”). Censuses of 2011 and 2021 were used as population reference for sex, age, and EA information for wave 1 and wave 2, respectively. Error bars correspond to 95% CIs from bootstrap analysis. PGS for EA was used for EA (**a**) and OS (**b**). PGS for height and body mass index (BMI) were used for the corresponding traits (**c** and **d**, respectively). PGS_EA_ was obtained from the PGI repository.

## Discussion

The cornerstone of science is reproducibility and replicability. Strong claims require strong evidence. However, many published social science results are false and attempts to replicate them often fail^44^. Furthermore, reproducibility alone is not enough; broader replicability is needed to ensure the robustness of scientific findings^45,46^.

The original work by Rimfeld et al. initiated a line of research comparing genetic associations with educational attainment between socialist and post-socialist societies^12,47^, and has since drawn attention beyond academic literature^48^. While the original study was performed according to the best standards of the field, it was limited by the methods, GWAS summary statistics and the EstBB sample size available at that time. In particular, the post-Soviet cohort comprised fewer than 2,500 individuals and was characterised by a bias towards individuals with a university degree (Supplementary Table 1). Thus, revisiting this study using updated methods and substantially expanded data from the EstBB is important not only for the Estonian context but also for the field as a whole. Besides attempting to validate the original findings in a larger sample using updated methods, we expanded the work by Rimfeld et al. in several directions. First, we traced the changes in heritability and PGS predictive accuracy not only in the two eras but also across narrower birth-year cohorts. Second, we added a third major line of evidence by directly modelling the interaction between PGS_EA_ and era while controlling for the interaction between PGS_EA_ and year of birth. Finally, we examined the extent to which our results could be driven by factors other than genuine changes in the genetic effects on SES-related traits, with particular attention to participation bias.

In general, our findings support the original report, showing that the relative genetic contribution to educational attainment is higher in independent Estonia than during the Soviet period. All three main analyses we performed (SNP heritability, polygenic score predictive accuracy, and interactions between era and polygenic score in a linear model) supported this conclusion. This convergence across methods strengthens the evidence that the finding is not specific to a particular measure of genetic associations. When exploring different cutoffs between Soviet and post-Soviet cohorts, phenotype definitions and transformations, and models for estimating heritability or polygenic score predictive accuracy, we almost always observed the same direction of the effect, with a higher genetic contribution in the post-Soviet cohort, although the statistical support for the difference varied depending on the details of the analysis. However, it is important to note that the observed differences were more modest than previously reported (Supplementary Table 18). Specifically, the estimated EA *h*^2^ was 106% higher in the post-Soviet than in the Soviet era in Rimfeld et al., compared with 20% in the present study using the 15-year cutoff. The corresponding increase in R^2^ was 60% in Rimfeld et al., compared with 11% in the present study using the 15-year cutoff; with the 10-year cutoff, the increase was 190% in Rimfeld et al., compared with 7% in the present study.

While the convergence across different approaches and model specifications adds credibility to our conclusion, several limitations of our study and similar studies should be considered. First and foremost is the generalisability of our findings to the general population. It is well documented that volunteer-based samples systematically differ from the wider population they are intended to represent^49–52^, a phenomenon known as participation bias. UK Biobank participants, for example, tend to be more educated, healthier, and have healthier lifestyles than the general UK population^49^. Importantly for our study, the extent of participation bias may depend on sex, age, or recruitment strategy^38,53^. Consequently, different subcohorts within a biobank may be biased in different ways relative to the underlying demographic groups in the populations. The Estonian Biobank provides a particularly relevant example because it comprises two recruitment waves: wave 1 recruited before 2016 and wave 2 after 2016. The predominant recruitment strategies differed between these waves, with wave 2 showing a more substantial “healthy volunteer bias” than wave 1^36^. When comparing recruitment waves within historical eras or within 5-year birth cohorts, we observed differences in height heritability within the Soviet cohort and in polygenic score predictive accuracy for OS within the post-Soviet cohort, particularly among individuals born between 1975 and 1985. These findings do not undermine our main conclusion as we do not see substantial between-wave differences in SNP heritability or polygenic score predictive accuracy for EA. However, they highlight that differential participation bias between subcohorts, related to on age, sex, or other characteristics, could produce apparent differences in genetic effects between groups within the studied cohort, even when such differences do not exist in the underlying population. Our resampling approach further illustrates that polygenic score predictive accuracy is particularly sensitive to changes in trait and PGS distributions due to participation bias. Weighting to account for non-representativeness in age, sex, and educational attainment reduced some of the between-wave differences but did not eliminate them, suggesting that additional, unmeasured aspects of participation may influence our estimates. This does not rule out a true change in the relative genetic contribution during the transition to the post-Soviet era, but suggests that its magnitude is difficult to estimate precisely in a volunteer-based biobank.

Another potential obstacle to interpreting our findings is that the participants in the Soviet cohort tend to be older at the time of recruitment (Supplementary Table 3). This could affect our analyses through survival bias, which may skew the distribution of both trait and PGS values in older cohorts^54^. In addition, the heritability of some traits, such as BMI, may change with age due to an increasing contribution of environmental factors^55,56^. However, such age-related changes in heritability are unlikely to substantially affect EA.

Finally, direct comparison of the EA phenotype across time is not straightforward. First, some educational categories have appeared or disappeared over time, such as the introduction of the master‟s degree in independent Estonia. Second, while uncommon, some participants assigned to the Soviet cohort could have pursued higher levels of education later in life after the collapse of the Soviet Union, meaning that part of their educational attainment may have been shaped by the post-Soviet environment. Third, changes in the distribution of the phenotype across cohorts may also contribute to our findings. In our data, later-born cohorts tend to have higher mean EA and, consequently, a more skewed distribution of years of education. Differences in the shape of the phenotype distribution can affect heritability estimates and PGS predictive accuracy even when the underlying genetic effects remain constant. Rank-based inverse normal transformation of EA and analysis of university degree on the liability scale partially mitigate these potential effects.

One important extension of the original study is a finer temporal resolution made possible by the larger sample, as well as the modelling of PGS interactions with both birth year and historical era. The analyses of SNP heritability and PGS predictive accuracy show that both measures change across birth years. This is true not only for EA and OS, but also height and BMI, traits that are less directly linked to socio-political changes. These findings suggest that the Soviet-to-post-Soviet transition is not the only factor contributing to temporal changes in the genetic contribution to variation in the analysed traits. Specifically, we observed an overall decrease in heritability and PGS predictive accuracy for EA as birth year increases. Such long-term changes in the genetic contribution to EA have also been observed in other societies, including those that did not experience similarly abrupt transformations, although the direction of change differs between populations and studies^57–61^.

However, in a linear model in which PGS_EA_ interacts with both birth year and era, we found that the effect of PGS_EA_ on EA increases in the post-Soviet era relative to the Soviet era, despite the overall decrease of PGS predictive accuracy with birth year. This provides stronger evidence for an independent effect of this transition, especially given that there is no significant effect of PGS×Era in the same model for height and BMI. One possible scenario consistent with our observations is that the transition from the Soviet to the post-Soviet era produced a period of disruption during which genetic differences became more predictive of educational attainment, but this effect weakened as the educational system, labour market, and broader social context stabilised.

The linear interaction analysis provides complementary evidence for a change associated with the Soviet-to-post-Soviet transition. It captures differences in the absolute genetic associations with the trait, rather than differences in the proportion of trait variance explained by genetic factors. In a linear model including interactions of PGS_EA_ with both birth year and era, the association of PGS_EA_ with EA was higher in the post-Soviet era than in the Soviet era, despite the overall decrease in the PGS_EA_ effect with birth year. Thus, this result converges with the heritability and PGS predictive accuracy analyses. Together, they indicate higher relative and also absolute genetic contributions to the EA variance in the post-Soviet era, strengthening the evidence that the observed difference reflects a real change associated with the transition rather than a feature of a particular analytical approach. Importantly, we found no significant PGS×Era interaction in the corresponding models for height or BMI. This suggests that the pattern is not simply a general temporal change affecting genetic associations with any trait, but is more closely related to EA and the social changes surrounding the transition. One possible scenario consistent with these observations is that the transition from the Soviet to the post-Soviet era produced a period of disruption during which genetic differences became more predictive of educational attainment, but that this effect weakened as the educational system, labour market and broader social context stabilised.

Overall, our findings provide evidence that genetic associations with educational attainment changed after the collapse of the Soviet Union. However, the estimated differences were substantially smaller than previously reported. Beyond the shift coinciding with the transition from the Soviet to the post-Soviet era, estimates of relative genetic contribution to educational attainment and other traits also varied within these time periods.

Our findings also highlight the need for caution when interpreting historical changes in the relative contribution of genetic factors to educational attainment. Such changes are often taken as evidence of greater equality of educational opportunity, on the assumption that environmental differences play a smaller role. However, this interpretation does not necessarily follow, as genetic associations with educational attainment may also reflect dynastic effects and other indirect genetic effects. Educational systems may amplify genetic differences by selecting or sorting individuals based on characteristics associated with their genotypes, even when those characteristics are unrelated to educational abilities. A better understanding of gene–environment interplay across development and historical contexts will therefore be essential for interpreting changes in the relative contribution of genetic factors to educational attainment and for identifying the social processes that shape educational opportunity.

## Methods

### Preregistered and actual study

The analysis plan was preregistered in the Open Science Framework prior to accessing the data (https://osf.io/ug5jp/).

The preregistered analyses were designed primarily to replicate and extend the findings of Rimfeld et al.^16^ using a similar methodological framework. SNP heritability, and PGS predictive accuracy were compared between individuals assigned to the Soviet and post-Soviet eras in Estonia. Differences in SNP heritability were assessed using a Z-test, and the proportion of phenotypic variance explained by PGSs was compared using bootstrap procedures.

The preregistered analyses were conducted as planned, with modifications to three analyses as described below. Preregistered fecundity analysis was not conducted because it had already been conducted and published elsewhere^62^. Additional analyses not included in the preregistered plan are identified as non-preregistered or post hoc. The methodology and results of preregistered analyses not presented in the main text are described in Supplementary Note 3.

The preregistered plan included several complementary analyses. We planned stratified analyses by sex and type of settlement (rural versus urban, with the urban category further divided into towns and cities). In addition to PGSs constructed from transmitted alleles, we planned to construct PGSs based on non-transmitted alleles inferred from genotyped trios using KING software. We also planned to evaluate the predictive accuracy of PGSs for cognitive and non-cognitive components of EA. We planned to estimate genetic correlations between EA in the EstBB and the GWAS sample used for PGS calculation. Height and body mass index (BMI) were planned to be analysed as control traits along with EA and OS. Phenotypic differences in EA and OS between the Soviet and post-Soviet eras were assessed using MANOVA.

For the main analyses, the Soviet and the post-Soviet era cohorts were defined using the preregistered plan specified birth-year cut-offs of 1976 and 1981, corresponding to 15 and 10 years in 1991, respectively. Individuals aged 25 years or younger at the time of data collection were excluded. Participants in the transition group, defined as those who were 16–25 years old in 1991, and other more finely defined birth cohorts were analysed in decade- and half-decade-based birth subcohort analyses. We also planned to estimate the genetic correlations between the Soviet and the post-Soviet era for each trait using bivariate GCTA-GREML. SNP heritability was estimated using unrelated individuals (stringent- and relaxed-threshold models) and a family-inclusive model, corresponding to “pedigree data” analysis specified in the preregistration.

Several deviations from the preregistered analysis plan were made. First, the preregistration specified an analysis of socioeconomic status (SES) as a composite measure based on EA and OS. We did not pursue this analysis in the revised study. A comprehensive SES measure would ideally incorporate additional components such as income and wealth. We did not have information on wealth and had income data for only a limited subset of participants. In addition, income measured at the time of data collection is less informative as a measure of lifetime socioeconomic position among older participants because it largely reflects pension income rather than peak or cumulative earnings. Because analysing a composite measure based only on EA and OS would not provide qualitatively different information from analysing these indicators separately, we focused on EA and OS as distinct indicators of socioeconomic position.

Second, although GCTA-GREML was specified in the preregistration, the primary SNP heritability analyses were performed using the GREML implementation in LDAK. We selected LDAK because its heritability model and implementation were considered more appropriate for the present analyses and computationally more efficient. We nevertheless repeated the principal analyses using GCTA-GREML and obtained consistent results, which provides a software-specific robustness check.

Third, the preregistration specified the construction of polygenic scores using the pipeline developed by Pain et al.^6364,65^. Third, we used precomputed PGSs from the PGI repository, some of which were based on GWAS summary statistics that are not publicly available and included the 23andMe cohort in the meta-analysis. We also constructed PGSs in-house using the same SBayesR method as a robustness analysis and to assess the extent to which the results depended on the inclusion of additional GWAS cohorts.

Forth, the analysis of fecundity, including comparisons between the Soviet and post-Soviet eras and analyses stratified by EA and EA PGS, was conducted separately and has been reported in an independent study^62^. We therefore cite that study but do not reproduce the fecundity results in the present manuscript.

In addition to the preregistered analyses and the deviations described above, we performed several additional analyses to further investigate the observed differences between eras, examine patterns at a finer temporal scale, assess the robustness of the findings to analytical choices, and address reviewers‟ comments during revision.

First, we performed G×Era analyses to investigate whether genome-wide genetic effects and genetic effects captured by PGSs differ between the Soviet and post-Soviet eras. REML G×Era analysis was used to estimate changes in genome-wide genetic effects, while linear regression analyses were used to investigate potential PGS×Era interactions. The latter models additionally included a PGS×Year of Birth interaction to account for temporal trends. Using these approaches, we further tested whether differences between the eras in SNP heritability and PGS predictive accuracy could be explained solely by changes in environmental variance or whether there was also evidence for changes in genetic effects.

Second, to examine whether the observed differences could be attributed specifically to the transition between the Soviet and post-Soviet eras rather than to broader temporal trends, we estimated SNP heritability and PGS predictive accuracy within decade- and half-decade-based birth cohorts. We also performed bandwidth analyses of SNP heritability using narrower birth-year windows around the transition.

Third, we investigated potential effects of participation bias. These analyses included stratification by wave of biobank enrolment, assessment of the relationship between phenotype distributions and PGS predictive accuracy, and analyses incorporating corrections for participation bias.

We also assessed the robustness of the findings to phenotype scaling and alternative analytical approaches. In addition to years of education as a continuous measure of EA, we analysed EA as a binary university-degree phenotype and after inverse normal transformation (INT). We also performed heteroscedastic regression as an alternative to ordinary least-squares regression. Bayesian analyses were conducted for the main comparisons to provide complementary inference for the principal findings.

### Participants

The participants were drawn from the Estonian Biobank (EsBB), a volunteer-based cohort of the Estonian adult population^35,36^. It includes genetic and phenotypic data on 210,438 individuals (72,708 men and 137,730 women), representing approximately 20% of Estonia’s contemporary adult population (14% men and 24% women). Participants’ year of birth ranges from 1905 to 2003 (mean, 1972; SD, 16.5). The participants were recruited over a two-decade period, from 2000 to 2021, across the country, covering all regions and a variety of different settings, thereby providing socio-economic and cultural heterogeneity. In addition to genetic and demographic data, participants provided health information, blood samples, and lifestyle details. Participants born outside the country were excluded from the analysis.

The sample was divided into two historical eras, the Soviet era and the post-Soviet era, following the preregistered plan and the methodology of Rimfeld et al.^16^. Estonia regained independence in 1991. For the primary 15-year cutoff, participants born in 1976 or later were assigned to the post-Soviet era and those born before 1976 to the Soviet era. We also used a 10-year cutoff, assigning participants who were aged 10 years or younger in 1991 to the post-Soviet group and those who were older to the Soviet group.

We additionally conducted analyses using narrower birth-year bins. In these analyses, 1976 was used as the boundary year between the Soviet and post-Soviet groups. The “transition” group described in the preregistration, comprising participants who were 16–25 years old in 1991, was also analysed within this framework. In the current report, we defined a birth cohort as a “transition” cohort when it overlapped with both historical eras (for example, a cohort born between 1971 and 1980).

For analyses comparing the full Soviet and post-Soviet eras, we included individuals older than 25 years at enrolment, as they were expected to have largely completed their highest level of education by the time of participation (Supplementary Note 1). However, this threshold may have been overly stringent because EstBB data are regularly synchronised with the population register. The 25-year age threshold was not applied in the decade- and half-decade-based birth-cohort analyses.

Sex was considered in the study design and determined using both self-reported and genetically inferred sex; individuals with a mismatch between the two were excluded during quality control. Sex was included as a covariate in the main analyses and used for sex-stratified analyses. Gender was not collected or analysed in this study. Accordingly, we use the term *sex* throughout the manuscript to refer to the biological sex variable used in the analyses.

### Ethics statement

The activities of the EstBB are regulated by the Human Gene Research Act, which was adopted in 2000 specifically for the operations of the EstBB. Individual-level data analysis in the EstBB was carried out under ethical approval “1.1-12/3593” from the Estonian Committee on Bioethics and Human Research (Estonian Ministry of Social Affairs), using data according to release application “4-1.6/GI/79” from the Estonian Biobank.

### Genotyping and quality control

Samples were genotyped on the Infinium Global Screening Array (GSA) of different versions (depending on the time of recruitment) with approximately 550,000 overlapping positions. Samples with <95% call rate or a mismatch between genetic and self-reported sex were excluded. Before the imputation step, all non-SNP polymorphisms and strand-ambiguous SNPs were filtered out. The final number of SNPs before the imputation step was 309,258. The genotypes were imputed with Beagle 5.4^66^ using the Estonian Reference panel as a reference set^67^.

Relationships were inferred with KING version 2.2.7^68^. Related individuals with a second-degree or closer relationship were excluded from the heritability analyses using the relaxed-threshold model, the PGS predictive accuracy analyses, and the linear regression analyses of interactions. KING output was also used to identify parent-offspring trios for the analysis of PGSs based on non-transmitted alleles.

### Ancestry and PCA

Genetic ancestry grouping was estimated using imputed genotypes with bigsnpr, following the original workflow^69^. For ancestry inference, genotypes were imputed using 1000 Genomes Project phase 3 samples^70^ for better representation of global haplotype diversity. Individuals from „Europe (East)‟, „Europe (North-West)‟ and „Finland‟ inferred ancestry groups were kept for further analysis. Next, individuals with self-reported ethnicity other than „Estonian‟ were excluded from the subsequent analyses, except for the reproduction of the results of Rimfeld et al.^16^, where only the genetic outliers were filtered out. These steps were implemented to recruit a relatively homogeneous group of participants with respect to genetics and culture.

A principal component analysis (PCA) was conducted to capture genetic structure. Before the analysis, genotypes were filtered for minor allele frequency >0.01, Hardy–Weinberg equilibrium (HWE) *p*-value >10^-5^ and missingness <0.05. Long-range linkage disequilibrium regions were removed^71^. Genotypes were pruned for linkage disequilibrium with PLINK2^72,73^ with a window size of 50kb, step 5kb and r^2^ threshold of 0.1. The first 100 principal components (PCs) were calculated on this SNP set using flashPCA version 2^74^.

### Phenotypes

Information on the highest level of education was obtained by taking the maximum reported value across the Estonian Population Register and the EstBB questionnaires. Continuous and binary traits corresponding to EA were considered. The continuous „years of education‟ phenotype was derived according to the ISCED 2011^75^ methodology. The link table for the reported level of education, ISCED 2011 and „years of education‟ is presented in Supplementary Table 19. Alternatively, attainment of a Bachelor’s (University) degree or higher was used as a binary phenotype (0 - not having a Bachelor‟s degree; 1 - having a Bachelor‟s degree or higher; not preregistered).

As a non-preregistered sensitivity analysis, EA was transformed to an inverse normal distribution before estimating SNP heritability, PGS performance and linear regression. EA was first residualised for sex, age, sex×age, age^2^, and 40 PCs. The residuals were then rank-based inverse normal transformed separately within each sex and era or more narrow birth cohorts.

Occupational status was assessed as in the study by Rimfeld et al. 2018^16^. The International Standard Classification of Occupations (ISCO)^76^ categories were used. The data from the questionnaire about “current professional status” and “main professional status” were combined. Military professions were excluded from the records. First, the mean value was calculated for each of the fields in case of multiple answers. Next, the mean of these values was used as the trait value.

Height and weight were measured by medical personnel. First available measurements were used. The age of the measurement was used as a covariate in the corresponding analyses. Height and Body Mass Index (BMI) were transformed to the logarithm scale to avoid the scale effect^77,78^. Individuals with values deviating from the means on at least one of the traits by more than 4 SD were excluded from the analysis.

### Heritability estimation and comparison

Trait heritability was estimated by LDAK version 6.2^21,79^, where not otherwise stated. The imputed genotypes were filtered using PLINK2 to retain only biallelic single-nucleotide polymorphisms (SNPs) with a minor allele frequency (MAF) >0.01, a Hardy–Weinberg equilibrium (HWE) *p*-value >10⁻⁵. Next, SNPs were thinned using a squared correlation (r^2^) threshold of 0.98 within a 100-kb window. Per-chromosome GRMs were calculated assuming the LDAK model, assuming equal weights and scaling parameter (--power) of -0.25. These per-chromosome GRMs were then merged into a single GRM. LDAK generalised REML (restricted maximum likelihood) solver was used to get heritability estimates. Sex, age, sex×age, age^2^, and 40 genetic principal components (PCs) were used as covariates.

Three models differing in their treatment of related individuals were used. First, a family-inclusive model based on the approach of Zaitlen et al.^37^ was applied, in which related and unrelated individuals were retained to maximise statistical power. This model uses two GRMs to partition phenotypic variance. The first GRM, calculated using all pairwise genetic relationships, captures variance attributable to the genotyped and imputed SNPs. The second GRM is derived by setting relatedness values below 0.05 to zero and captures additional variance attributable to genetic effects not captured by the genotyped or imputed SNPs. This component may also capture phenotypic covariance between relatives due to shared environmental factors, which may be particularly relevant for socioeconomic traits. We therefore did not interpret or analyse this component and focused on the SNP heritability estimated from the first GRM.

The second and third models used the standard approach and included only unrelated individuals, using different thresholds to define relatedness. In the stringent-threshold model, individuals from pairs with a GRM relatedness estimate ≥0.05 were excluded. In the relaxed-threshold model, individuals with a second-degree or closer genetic relationship, as inferred with KING, were excluded.

For a complementary heritability analysis with GCTA-GREML^34^ (GCTA version 1.94.1), the genotyped SNP data were used (following methodology from Rimfeld et al.^16^) after keeping SNPs with minor allele frequency >0.01, Hardy–Weinberg equilibrium (HWE) *p*-value >10^-5^ and missingness <0.015. GRMs were calculated per chromosome and merged afterwards. GCTA-GREML was used to estimate SNP heritability. Sex, age, sex×age, age^2^, and 10 PCs were used as covariates. A relatedness threshold of 0.05 was used.

The binary EA (University degree) heritability estimates were transformed from the observed to the liability scale based on census population prevalence^80,81^ using the Robertson transformation^82^.

Heritability estimates were compared with a paired Z-test.

### Bandwidth analysis (not preregistered)

Bandwidth analysis was performed to allow for flexible birth-year ranges when defining cohorts and to assess the potential effect of the transition from the Soviet to the post-Soviet era. Birth-year windows ranging from 2 to 20 years on each side of the 1976 boundary were used. For each bandwidth, individuals within the specified number of years on either side of the boundary were included, and SNP heritability was estimated separately for the Soviet and post-Soviet eras using the same LDAK approach and covariates as in the main analysis. The difference in heritability between eras was then calculated for each bandwidth.

### G**×**Era REML analysis (not preregistered)

To test whether the genome-wide contribution to EA differed between the Soviet and post-Soviet eras, we performed a G×E analysis using the REML framework implemented in LDAK, following the approach described in the original GCTA G×E analysis^34^. Individuals with a pairwise genetic relatedness greater than 0.05 were excluded. The model included separate genetic relationship matrices representing the overall genetic effect and its interaction with Era, allowing the genetic variance to differ between the Soviet and post-Soviet groups. The G×E variance component was used to test whether genetic effects differed between eras. The analysis was performed in the combined sample using the same phenotype, covariates and genetic data processing as the main heritability analyses.

### Polygenic score calculation

In the polygenic score analyses, two sets of PGSs were used. The first set was obtained from the PGI repository^39^. There, PGS weights were calculated using SBayesR based on summary statistics from: the EA4 GWAS^25^, which included the 23andMe cohort but excluded the EstBB cohort from the meta-analysis; the meta-analysis of UK Biobank and 23andMe cohorts for height; and the UK Biobank GWAS for BMI.

The second set of the PGSs was calculated by this study. EA PGS was calculated using summary statistics from EA4, with Estonian individuals and the 23andMe cohort excluded from the meta-analysis^25^. The height and BMI PGSs were calculated using summary statistics from GWAS in the European ancestry cohort of the UK Biobank conducted by the Pan-UKBB project^83^. To create polygenic scores, we extracted a set of 1,075,599 autosomal HapMap 3 SNPs with a minor allele count >5 and info score >0.7 in the EstBB. Polygenic scores were calculated using SBayesR^40^ (gctb_2.02) with default parameters (--pi 0.95,0.02,0.02,0.01 --gamma 0.0,0.01,0.1,1 --chain-length 10000 --burn-in 2000 --out-freq 10), including an LD matrix built using data on 50,000 UK Biobank participants^40^.

### Polygenic score predictive accuracy comparison

To assess the PGS performance, the incremental R^2^ was calculated using sex, age, sex×age, age^2^, and 40 PCs as covariates. Alternatively, traits were pre-adjusted for these covariates, and then R^2^ was derived from a univariate linear regression with PGS as the predictor. 95% confidence intervals (95% CIs) were calculated using bootstrap with 1000 replicates. Two-tailed *p*-values were derived from the same bootstrap replicates. With 1000 bootstrap replicates, the smallest attainable *p*-value was <0.002. Incremental R^2^ 95% CIs for half-decade birth cohorts were calculated using Fisher z-transformation, assuming that incremental R^2^ is approximately equal to the squared correlation coefficient.

### Linear regression of PGS×Era interactions (not preregistered)

To assess whether the associations between the phenotypes and the corresponding polygenic scores differed between the Soviet and post-Soviet eras, we conducted interaction analyses using linear and logistic regression models.

First, we modelled EA (years of education) and OS (treated as continuous), height and BMI as dependent variables in unrelated individuals (excluding second-degree and closer relatives). The primary ordinal least squares linear regression model included main effects of PGS, era (coded 0 for the Soviet era and 1 for the post-Soviet era), and their interaction term (PGS×Era). The model additionally controlled for participation wave (0 = wave 1; 1 = wave 2), year of birth (YoB, z-standardised), sex (0 = male; 1 = female), and all pairwise interactions among these covariates, including a quadratic YoB term. Following recommendations for modelling G×E interactions^41^, we also included interaction terms between PGS and wave, YoB, and sex to reduce confounding due to omitted interactions. Because era is a binary variable derived from YoB, we also estimated models that excluded the PGS×YoB term.

Analyses were performed in the full sample and separately within each participation wave. The statistical significance of the PGS×Era term was used to evaluate whether PGS effects differed between eras.

To assess robustness to treating ordinal traits as continuous^42^, we repeated the analyses using logistic regression with binary outcomes. EA was dichotomised as the presence vs. absence of a university degree. OS was dichotomised by grouping managers, professionals, and technicians and associate professionals (ISCO major groups 1–3) versus all other occupational categories (excluding “unclassified” and “armed forces”), yielding approximately balanced classes.

All models included 40 genetic principal components to control for population structure.

For EA, we additionally performed heteroscedastic regression with inverse normal-transformed years of education using R package *gamlss*^84^. These models jointly estimated the effects of predictors on the conditional mean and conditional variance of EA, with the same covariates included in both components.

### Statistical significance

Bonferroni-corrected significance thresholds (α) were defined as 0.05 divided by the number of tests. For the main analyses combining waves 1 and 2, four independent tests were considered, corresponding to the four traits analysed. For analyses testing differences between eras within each wave, or differences between waves within each era, eight independent tests were considered. Analyses using alternative phenotype scales and sensitivity analyses were not treated as independent tests; therefore, the same significance thresholds were applied to these analyses. Raw *p*-values are reported alongside the corresponding thresholds.

### Bayesian analysis (not preregistered)

Bayesian analyses were performed for the primary comparisons of the Soviet and post-Soviet eras for EA. We estimated the posterior distribution of the difference between eras using a normal prior centred at zero with a weakly informative, wide standard deviation (0.05 for *h^2^_SNP_* and PGS R^2^; 0.032-0.036 for β_PGSxEra_, equal to s.e. of β_PGS_; Supplementary Table 6). The prior therefore expressed no directional expectation while allowing a broad range of effect sizes. Posterior means and 95% credible intervals were calculated, together with the posterior probability that the difference was greater than zero. Bayesian analyses were used as a complementary assessment of the primary frequentist results.

### Genetic correlations

To estimate the genetic correlation between EstBB EA and EA4 meta-analysis EA, we first conducted a GWAS in the EstBB cohort. We used the SAIGE method^85^ following the guidelines provided. Genetic correlation was estimated from the GWAS summary statistics using LD score regression with default LD scores derived from 1000 Genomes European ancestry individuals^86^.

Genetic correlations between the Soviet and the post-Soviet eras (not preregistered) were estimated for EA, OS, height and BMI using the bivariate GCTA-GREML analysis following the methodology described in the “Heritability estimation and comparison” section for the GCTA-GREML analysis.

### Matching the EA distribution (not preregistered)

To evaluate how differences in the distribution of EA affect the predictive performance of PGS_EA_, we conducted a resampling analysis across four subcohorts defined by recruitment wave and era. First, the EA phenotype was pre-adjusted for sex, age, sex×age, age^2^, and 40 genetic principal components by regressing these covariates on EA and using the residuals. For each subcohort (“source group”), individuals were randomly sampled with replacement to create new subsamples whose EA distribution matched that of a different subcohort (“target group”). Matching was performed either by EA alone or jointly by EA and sex. For each subsample, the variance explained by the PGS_EA_ was quantified as the squared Pearson correlation between the pre-adjusted EA residuals and the PGS, which is equivalent to R^2^ in a univariate linear model. This procedure was repeated 1,000 times for each source-target combination to generate distributions of R^2^ values. These distributions were compared with the original R^2^ values of the source and target subcohorts to assess how differences in EA (and sex) distributions influence PGS performance.

### Correction for participation bias (not preregistered)

No established procedure for correction for participation bias exists for the EstBB. Thus, we used three variables known to have a strong effect on participation in biobank initiatives, whose summary data are easily accessible for the corresponding population: age, sex and EA^49,53^. Increasing the number of variables used for the correction decreases the effective sample size in further analyses. To explore how this affects the results, two weighting schemes were implemented: (1) marginal adjustment, where the EA distribution within each sex and age stratum was matched to that of the census; and (2) joint adjustment, where the full joint distribution of age, sex, and EA matched that in the census population.

We used the 2011 census data^80^ as a population reference for wave 1 of participation and the 2021 census data^81^ for wave 2. This is justified by the time between census data collection and biobank participation, as well as by the fact that participants kept in the study were older than 25 at the time of enrolment and had therefore mostly completed their highest level of education. The age groups were divided into 5-year bins. EA had three categories: primary, secondary, and tertiary education. The post-stratification weighting method was used for weight calculation, further used in weighted linear regression to calculate incremental R^2^. Bootstrap procedure with 1000 replicates was used to calculate R^2^ 95% CIs. In each replicate, a subsample with replacement was selected, after which weighting was applied.

### AI-assisting writing

GPT-5.6 Luna (OpenAI) was used for language editing to improve readability of the manuscript. All scientific content, analyses and interpretations were produced by the authors. The final manuscript was reviewed and approved by the authors.

## Data availability

Access to the Estonian Biobank data (https://genomics.ut.ee/en/content/estonian-biobank) is restricted to approved researchers and can be requested.

## Code availability

Custom R code used for statistical analyses is available on GitHub (https://github.com/ivkuz/HeritabilityChangeEst).

## Acknowledgements

We acknowledge the participants of the Estonian Biobank. The research was conducted using the Estonian Center of Genomics/Roadmap II, funded by the Estonian Research Council (project number TT17). Data analysis was carried out in part in the High-Performance Computing Center of the University of Tartu. IK and AM were supported by the Centre of Excellence for Personalised Medicine, funded by grant TK214 from the Estonian Ministry of Education and Research. KR was supported by a UK Medical Research Council Grant UKRI1503. KL was supported by the Estonian Research Council (grant no. PSG615) and by the Estonian Centre of Excellence for Well-Being Sciences (EstWell), funded by grant TK218 from the Estonian Ministry of Education and Research. PH was funded by the Estonian Research Council grant PRG1137.

## Author contributions

I.A.K., K.L., V.P., and K.R. designed the study. The Estonian Biobank Research Team and A.M. collected and provided the genotype and phenotype data used in the study. I.A.K. performed computational analyses. All authors except I.A.K. and V.P. contributed to the preregistration. V.P. and K.R. supervised the study. I.A.K., V.P., and K.R. wrote the first draft of the manuscript. All other authors critically reviewed the manuscript and provided feedback.

