## Supplementary Notes, Tables and Figures for "Revisiting genetic associations with educational outcomes during and after the Soviet era in Estonia"

Supplementary Material

Ivan A. Kuznetsov, Margherita Malanchini, Oliver Pain, Jonathan Coleman, Philip S. Dale, Jean-Baptiste Pingault, Kathleen Rastle, Estonian Biobank Research Team, Peeter Hõrak, Robert Plomin, Andres Metspalu, Kelli Lehto, Vasili Pankratov, Kaili Rimfeld

Supplementary Table 13. Comparison of variance explained by PGS from the PGI repository in the post-Soviet and Soviet eras, and the eras additionally divided by the wave of participation, with the 15-year era cutoff. ..23

### Supplementary Note 1. Age at biobank enrolment threshold

The analyses comparing the Soviet and post-Soviet subcohorts were restricted to individuals aged at least 25 years old at the time of biobank enrolment to minimise misclassification due to incomplete education. This threshold represents a compromise between ensuring completed education and maximising sample size. According to OECD statistics, the average age at first university graduation is approximately 23 years, and about 75% of first-degree graduates complete their studies before the age of 25[1]. Although some individuals continue their education beyond this age, they constitute a relatively small proportion of the population. For example, individuals with doctoral degrees account for approximately 1% of the Estonian population[2].

Furthermore, educational attainment in the Estonian Biobank is linked to the national population register, allowing education records to be updated after enrolment, although updates may not always be complete. Consequently, educational attainment at the time of analysis does not necessarily reflect the status at enrolment.

As an additional sensitivity analysis, we examined university degree (binary EA), which is less sensitive to educational progression after age 25 because most bachelor's degrees are completed before this age. It is also less affected by changes in the structure of higher education, such as the transition to the Bologna model[3]. The consistency of the results for this phenotype further suggests that incomplete education is unlikely to explain the observed differences between the Soviet and post-Soviet cohorts.

### Supplementary Note 2. Reproduction of Rimfeld et al. (2018)

In this note, we reproduce the analyses of educational attainment (EA) polygenic score prediction ($PGS_{EA}$ $R^2$) and SNP heritability ($h^2_{SNP}$) reported in Rimfeld et al. (2018)[4] using the original Estonian Biobank subsample used in that study.

#### Analysis of heritability

We first reproduced the SNP heritability analyses, focusing primarily on EA, with occupational status (OS) examined for completeness. Rimfeld et al. adjusted EA for sex and age and then applied a rank-based inverse normal transformation (INT). We repeated this approach using GCTA-GREML[5]. Alternatively, we estimated $h^2_{SNP}$ using the untransformed phenotype while including sex and age as covariates directly in the model.

The original study did not restrict the sample by self-reported ethnicity. Here, we report results for (i) all individuals of European genetic ancestry and (ii) only self-reported Estonians. The patterns of results were highly similar across these specifications (Supplementary Fig. 33, Supplementary Table 20).

In line with the original findings, we did not observe statistically significant differences in $h^2_{SNP}$ between the Soviet and post-Soviet eras under any era cutoff. EA $h^2_{SNP}$ in the post-Soviet era, with a cutoff of 10 years, was not significantly different from zero and was not reported in the original study. For the 15-year cutoff, the point estimate of $h^2_{SNP}$ in the post-Soviet era was higher than in the Soviet era, mirroring the original report, but the difference was not statistically significant in either analysis.

We further estimated $h^2_{SNP}$ using the LDAK model[6,7], which incorporates LD structure and minor allele frequency (MAF). Here, imputed genotype data were used for kinship estimation, resulting in higher $h^2_{SNP}$ estimates overall due to the inclusion of more variants. Although the LDAK estimates for the two eras were closer in magnitude compared with GCTA, large standard errors precluded meaningful comparison. Similarly, wide confidence intervals prevented strong conclusions regarding OS $h^2_{SNP}$.

**Predictive accuracy of a polygenic score**

We next assessed whether the predictive accuracy of $PGS_{EA}$ differed between the Soviet and post-Soviet eras. To maximise power, we used the $PGS_{EA}$ from the PGI repository (with Estonian Biobank sample excluded from the meta-analysis). Incremental $R^2$ was estimated using models that included 10 genetic principal components (40 in the main analysis) and demographic covariates (sex, age, sex×age, $age^2$). As an alternative approach, we pre-adjusted the phenotype for covariates prior to model fitting. Standard errors for $R^2$ were obtained via bootstrapping.

$R^2$ was higher in the post-Soviet era than in the Soviet era across all combinations of subsamples, traits, and $R^2$ calculation approaches (Supplementary Figs. 34 and 35, and Supplementary Table 21). These differences were statistically significant for EA measured in years of education in all cases ($\alpha$ = 0.05, following the original study). Direct comparison with the original study is complicated due to several methodological differences. First, the genotypes in our sample were regenotyped and reimputed, which may affect SNP quality and imputation accuracy. Second, we used a PGS based on the most recent GWAS summary statistics from EA4, whereas the original study was based on EA2; this introduces differences in SNP effect sizes and coverage. Third, SBayesR was used for current PGS construction, which differs from the method applied in the original analysis. All these factors can influence the predictive power of the PGS and the resulting $R^2$ estimates.Nevertheless, in line with the original study, we observed a substantial increase in PGS predictive accuracy for OS and, especially, EA in the post-Soviet era when using the original study sample.

### Supplementary Note 3. Preregistered analyses not included in the main text

In this note, we present the results of the analyses listed in the preregistration plan on the Open Science Framework (https://osf.io/ug5jp/), which are not included in the main manuscript text.

#### MANOVA

We used multivariate analysis of variance (MANOVA) to assess differences between the Soviet and post-Soviet eras in educational attainment and occupational status, with era as the independent variable and EA and OS as joint outcomes. It revealed significant multivariate differences between the Soviet and post-Soviet eras (Pillai's trace = 0.012, $F_{2,73304} = 430$, $p < 2.2\times10^{-16}$). Univariate follow-up analyses showed that cohort differences were significant for both EA ($F_{1,73305} = 645$, $p < 2.2\times10^{-16}$) and OS ($F_{1,73305} = 709$, $p < 2.2\times10^{-16}$).

Similar multivariate and univariate differences were observed when the analyses were performed separately for wave 1 (Pillai's trace = 0.015, $F_{2,24568} = 192$, $p < 2.2\times10^{-16}$) and wave 2 (Pillai's trace = 0.005, $F_{2,57008} = 137$, $p < 2.2\times10^{-16}$), with both EA and OS differing significantly between the Soviet and post-Soviet cohorts in each phase (all $p < 2.2\times10^{-16}$).

Despite significant differences between the eras, low Pillai's trace values indicate that the effects of era on mean EA and OS are relatively small.

#### Predictive accuracy of polygenic scores for cognitive and non-cognitive components of educational attainment

We investigated whether the predictive accuracies of polygenic scores for cognitive and non-cognitive components of educational attainment differ between the Soviet and post-Soviet eras. This was motivated by the hypothesis that societal transformation may have altered the relative importance of cognitive and non-cognitive traits in shaping educational attainment, thereby changing the predictive accuracy of the corresponding polygenic scores for EA and OS. Separate polygenic scores capturing the cognitive ($PGS_{Cog}$) and non-cognitive ($PGS_{NonCog}$) components of EA were calculated using summary statistics from the GWAS-by-subtraction with the Estonian Biobank cohort excluded from the meta-analysis[8].

We estimated incremental $R^2$ for EA and OS and $R^2$ for the pre-adjusted traits (Supplementary Fig. 36, Supplementary Table 22). Significant differences between the eras were observed for EA with both $PGS_{Cog}$ and $PGS_{NonCog}$. Notably, the directions of these differences were opposite: $PGS_{Cog}$ explained more variance in the Soviet era while $PGS_{NonCog}$ explained more variance in the post-Soviet era. No significant between-era differences were observed for OS.

These findings should be interpreted with caution. Although the genetic correlation between the cognitive and non-cognitive EA components is zero by construction in the GWAS-by-subtraction framework, $PGS_{Cog}$ and $PGS_{NonCog}$ are correlated (r = -0.25 in the Estonian Biobank). Consequently, the opposite direction of the between-era differences in $R^2$ may partly reflect this correlation rather than genuine differences in the contributions of cognitive and non-cognitive genetic factors.

**Predictive accuracy of non-transmitted-allele polygenic scores**

Besides direct genetic effects, parental (or dynastic) effects contribute substantially to socioeconomic traits, including EA and OS[9]. We used genotyped parent-offspring trios from the Estonian Biobank (n = 4,542 for EA; n = 3,263 for OS; n = 4,275 for height; n = 4,275 for BMI) to evaluate the predictive accuracy of polygenic scores based on non-transmitted parental alleles. Higher $R^2$ of the non-transmitted PGS is consistent with stronger parental (dynastic) effects and implies a greater contribution of these indirect effects to the predictive accuracy of PGS based on transmitted alleles (Supplementary Fig. 37, Supplementary Table 23). Owing to the limited sample size, this analysis had limited statistical power. We did not detect significant differences between the Soviet and post-Soviet eras, either in the full sample or within subcohorts by participation wave. However, for EA and OS, the $R^2$ estimates were relatively high in the post-Soviet era in wave 1. This pattern suggests that parental effects may be stronger in this specific cohort and could partly contribute to the higher predictive accuracy of the transmitted $PGS_{EA}$.

#### Supplementary Note 4. Absolute genetic and environmental trait variance across birth cohorts

Change in heritability can occur when genetic variance relevant to the trait, environmental variance or both change. Trait variance unexplained by the genetic relatedness matrix can be approximated as environmental variance, although it also includes genetic variance not captured by genotyped SNPs and variation due to other types of polymorphisms. We compared the trends in genetic (G) and residual ("environmental", E) variance for each trait using the stringent-threshold heritability model. For EA, residual variance remained relatively stable around the era transition, whereas genetic variance increased, suggesting that the increase in heritability was driven by increased genetic (not necessarily direct) effects on EA rather than by increased environmental homogeneity (Supplementary Fig. 38).

For OS and BMI, genetic variance was relatively stable, whereas residual variance changed over time. In contrast, for height, the residual variance was relatively stable whereas genetic variance increased consistently across birth cohorts.

### Supplementary Tables

**Supplementary Table 1. Proportion of individuals with different levels of highest education achieved in the Estonian Biobank and in the Census of a corresponding year.** Individuals born before 1976, and therefore older than 15 at the time of the collapse of the Soviet Union, belong to the Soviet era, while younger individuals belong to the post-Soviet era. Recruitment wave 1 spanned from 2000 to 2016, and wave 2 spanned from 2017 to 2021. The National Censuses of 2011 (wave 1) and 2021 (wave 2) were used as the population reference.

| Wave | Era | Sample | Educational Attainment | | |
|---|---|---|---|---|---|
| | | | Primary, % | Secondary, % | Tertiary, % |
| 1 | Soviet | Census 2011 | 22 | 42.8 | 35.3 |
| 1 | Soviet | Rimfeld et al. | 3.9 | 65.9 | 30.2 |
| 1 | Soviet | EstBB | 15.1 | 57.3 | 27.6 |
| 1 | post-Soviet | Census 2011 | 17.1 | 43.7 | 39.2 |
| 1 | post-Soviet | Rimfeld et al. | 0.4 | 49.5 | 50.1 |
| 1 | post-Soviet | EstBB | 11.9 | 46.6 | 41.5 |
| 2 | Soviet | Census 2021 | 15.3 | 44.6 | 40.1 |
| 2 | Soviet | EstBB | 3.2 | 47.9 | 48.9 |
| 2 | post-Soviet | Census 2021 | 14.8 | 42.1 | 43.1 |
| 2 | post-Soviet | EstBB | 5 | 35.3 | 59.6 |

**Supplementary Table 2. Sample size for each trait and subcohort.** Individuals born before 1976 (cutoff 15) or 1981 (cutoff 10) belong to the Soviet era, while younger individuals belong to the post-Soviet era. Recruitment wave 1 spanned from 2000 to 2016, and wave 2 spanned from 2017 to 2021. Individuals aged 25 years or younger at the time of data collection are excluded.

| | **Soviet** | **post-Soviet** | **Soviet - wave 1** | **post-Soviet wave 1** | **Soviet - wave 2** | **post-Soviet wave 2** |
|---|---|---|---|---|---|---|
| **Complete subcohort sample sizes** | | | | | | |
| **Cutoff 15** | | | | | | |
| EA | 92,715 | 53,786 | 26,148 | 4,290 | 66,567 | 49,496 |
| OS | 67,726 | 37,150 | 25,777 | 4,097 | 41,949 | 33,053 |
| Height | 89,289 | 50,589 | 26,075 | 4,265 | 63,214 | 46,324 |
| BMI | 89,289 | 50,589 | 26,075 | 4,265 | 63,214 | 46,324 |
| **Cutoff 10** | | | | | | |
| EA | 109,875 | 36,626 | 29,053 | 1,385 | 80,822 | 35,241 |
| OS | 80,120 | 24,756 | 28,562 | 1,312 | 51,558 | 23,444 |
| Height | 105,547 | 34,331 | 28,960 | 1,380 | 76,587 | 32,951 |
| BMI | 105,547 | 34,331 | 28,960 | 1,380 | 76,587 | 32,951 |
| **Sample sizes after excluding second-degree or closer relatives** | | | | | | |
| **Cutoff 15** | | | | | | |
| EA | 62,998 | 43,496 | 20,977 | 4,087 | 49,604 | 40,550 |
| OS | 49,684 | 31,644 | 20,718 | 3,908 | 34,303 | 28,529 |
| Height | 60,494 | 40,865 | 20,885 | 4,061 | 47,129 | 37,939 |
| BMI | 60,494 | 40,865 | 20,885 | 4,061 | 47,129 | 37,939 |
| **Cutoff 10** | | | | | | |
| EA | 68,332 | 29,324 | 22,718 | 1,363 | 54,400 | 28,241 |
| OS | 56,139 | 21,921 | 22,494 | 1,297 | 40,365 | 20,810 |
| Height | 69,079 | 29,343 | 22,738 | 1,363 | 55,144 | 28,260 |
| BMI | 69,079 | 29,343 | 22,738 | 1,363 | 55,144 | 28,260 |

**Supplementary Table 3. Descriptive statistics for age, sex, and SES-related variables.** Individuals born before 1976 (cutoff 15) or 1981 (cutoff 10) belong to the Soviet era, while those born after belonging to the post-Soviet era. Recruitment wave 1 spanned from 2000 to 2016, and wave 2 spanned from 2017 to 2021. Age was calculated as the difference between 2022 and the year of birth for individuals alive in 2022, or between the year of death and the year of birth for deceased individuals. Sample size is wave-era specific, regardless of the phenotype data available. 'N' denotes the complete sample sizes of the corresponding groups, 'N unrelated' denotes the sample sizes after excluding second-degree or closer relatives. Individuals aged 25 years or younger at the time of data collection are excluded. Trait-specific sample size information is presented in Supplementary Table 2.

| Wave | Era | N | N unrelated | Female proportion, % | Mean age | SD age | Mean EA | SD EA | Mean OS | SD OS | Mean height | SD height | Mean BMI | SD BMI |
|---|---|---|---|---|---|---|---|---|---|---|---|---|---|---|
| **Cutoff 15** | | | | | | | | | | | | | | |
| All | Soviet | 93,981 | 63,910 | 66.1 | 61.5 | 10.4 | 16.1 | 3.2 | 5.9 | 2.2 | 170.1 | 9.1 | 27.4 | 5.1 |
| All | post-Soviet | 53,840 | 43,533 | 62.5 | 38.2 | 5.0 | 16.6 | 3.1 | 6.3 | 2.0 | 173.2 | 9.2 | 24.9 | 4.7 |
| 1 | Soviet | 26,174 | 20,997 | 66.2 | 66.0 | 12.4 | 15.4 | 3.3 | 5.5 | 2.4 | 168.9 | 9.1 | 27.4 | 5.3 |
| 1 | post-Soviet | 4,293 | 4,088 | 61.5 | 42.7 | 2.5 | 16.4 | 3.2 | 6.1 | 2.3 | 172.8 | 9.2 | 24.5 | 4.6 |
| 2 | Soviet | 67,807 | 50,516 | 66.1 | 60.7 | 9.8 | 16.3 | 3.2 | 6.1 | 2.1 | 170.3 | 9.1 | 27.4 | 5.1 |
| 2 | post-Soviet | 49,547 | 40,585 | 62.5 | 37.9 | 4.9 | 16.6 | 3.1 | 6.4 | 2.0 | 173.2 | 9.2 | 25.0 | 4.7 |
| **Cutoff 10** | | | | | | | | | | | | | | |
| All | Soviet | 111,168 | 69,064 | 65.8 | 58.5 | 11.4 | 16.2 | 3.2 | 6.0 | 2.2 | 170.5 | 9.1 | 27.1 | 5.1 |
| All | post-Soviet | 36,653 | 29,333 | 62.8 | 35.6 | 3.5 | 16.7 | 3.1 | 6.3 | 2 | 173.5 | 9.3 | 24.6 | 4.5 |
| 1 | Soviet | 29,082 | 22,737 | 65.8 | 63.8 | 13.5 | 15.5 | 3.3 | 5.5 | 2.4 | 169.3 | 9.2 | 27.1 | 5.3 |
| 1 | post-Soviet | 1,385 | 1,362 | 60.6 | 39.9 | 1.3 | 16.9 | 3.1 | 6.3 | 2.2 | 173.2 | 9.2 | 24.1 | 4.4 |
| 2 | Soviet | 82,086 | 55,130 | 65.7 | 57.5 | 10.8 | 16.4 | 3.1 | 6.1 | 2.1 | 170.8 | 9.1 | 27.1 | 5.1 |
| 2 | post-Soviet | 35,268 | 28,250 | 62.9 | 35.5 | 3.5 | 16.7 | 3.1 | 6.3 | 2.0 | 173.5 | 9.3 | 24.6 | 4.5 |

**Supplementary Table 4. Genetic correlation between the analysed traits in the Soviet and post-Soviet eras.** The estimates were obtained using bivariate GREML-GCTA with 15- and 10-year era cutoffs. The Bonferroni-corrected significance level for testing whether the genetic correlation differed from 1 (one-tailed test) was $\alpha = 0.0125$, corresponding to four independent tests.

| **Trait** | **Cutoff, age in 1991** | **rG** | **SE** | ***p*-value** |
|---|---|---|---|---|
| EA | 15 | 0.969 | 0.115 | 0.395 |
| EA | 10 | 1.000 | 0.131 | 0.500 |
| OS | 15 | 0.905 | 0.357 | 0.402 |
| OS | 10 | 0.528 | 0.222 | 0.053 |
| Height | 15 | 0.990 | 0.038 | 0.392 |
| Height | 10 | 0.978 | 0.044 | 0.301 |
| BMI | 15 | 0.984 | 0.069 | 0.410 |
| BMI | 10 | 0.906 | 0.077 | 0.125 |

**Supplementary Table 5. Comparison of heritability estimates in the Soviet and post-Soviet eras (age in 1991 - cutoff of 15) and the eras additionally divided by the wave of participation.** Abbreviations: EA, educational attainment; OS, occupational status; BMI, body mass index; University, university degree (on liability scale); INT EA, inverse normal-transformed educational attainment. Two-tailed Wald test *p*-values are reported. The Bonferroni-corrected significance level was $\alpha = 0.0125$, corresponding to four independent tests, in analyses of the combined wave 1 and wave 2 samples and $\alpha = 6.25\times10^{-3}$, corresponding to eight independent tests, in analyses stratified by wave. *P*-values below the corresponding significance threshold are shown in bold.

| Model | Trait | Soviet vs. post-Soviet | | | Soviet | post-Soviet |
|---|---|---|---|---|---|---|
| | | All | Wave 1 | Wave 2 | Wave 1 vs. Wave 2 | Wave 1 vs. Wave 2 |
| Family-inclusive | EA | **$2.7\times10^{-4}$** | 1.00 | 0.02 | 0.74 | 0.81 |
| Stringent-threshold | EA | **$4.5\times10^{-3}$** | 0.79 | 0.74 | 0.98 | 0.73 |
| Relaxed-threshold | EA | **$2.7\times10^{-3}$** | 0.990 | 0.014 | 0.890 | 0.69 |
| Stringent-threshold | INT EA | **$9.4\times10^{-4}$** | 0.440 | 0.048 | 0.960 | 0.22 |
| Family-inclusive | University | 0.068 | 0.330 | 0.280 | 0.300 | 0.37 |
| Stringent-threshold | University | 0.550 | 0.150 | 0.280 | 0.620 | 0.29 |
| Relaxed-threshold | University | 0.19 | 0.51 | 0.3 | 0.32 | 0.57 |
| Family-inclusive | OS | 0.36 | 0.53 | 0.38 | 0.68 | 0.69 |
| Stringent-threshold | OS | 0.27 | 0.65 | 0.39 | 0.055 | 0.72 |
| Relaxed-threshold | OS | 0.27 | 0.3 | 0.48 | 0.057 | 0.65 |
| Family-inclusive | Height | **$5.6\times10^{-10}$** | 0.084 | **$7.4\times10^{-6}$** | **$3.3\times10^{-5}$** | 0.77 |
| Stringent-threshold | Height | **$9.0\times10^{-4}$** | 0.24 | 0.021 | **$4.1\times10^{-3}$** | 0.83 |
| Relaxed-threshold | Height | **$1.5\times10^{-4}$** | 0.065 | 0.011 | **$3.3\times10^{-5}$** | 0.61 |
| Family-inclusive | BMI | **$1.3\times10^{-3}$** | 0.7 | **$1.4\times10^{-3}$** | 0.49 | 0.48 |
| Stringent-threshold | BMI | 0.17 | 0.57 | 0.35 | 0.14 | 0.74 |
| Relaxed-threshold | BMI | **$8.0\times10^{-3}$** | 0.94 | $7.5\times10^{-3}$ | 0.99 | 0.71 |

**Supplementary Table 6. Bayesian estimates for the main analyses of differences in genetic effects between the Soviet and post-Soviet eras.** P(pS > S) denotes the posterior probability that the corresponding statistic is larger in the post-Soviet era than in the Soviet era. "CI 95 lower" and "CI 95 upper" denote the lower and upper bonds of 95% credible interval.

| Statistic | Comment | Estimate | SE | Prior Mean | Prior SD | Posterior Mean | Posterior SD | CI 95 lower | CI 95 upper | *p*-value (pS > S) |
|---|---|---|---|---|---|---|---|---|---|---|
| $h^2$ | — | 0.036 | 0.010 | 0 | 0.05 | 0.035 | 0.010 | 0.016 | 0.054 | 0.9998 |
| $R^2$ | Incremental | 0.011 | 0.010 | 0 | 0.05 | 0.011 | 0.010 | -0.008 | 0.030 | 0.8704 |
| $\beta_{PGSxEra}$ | PGSxYoB covariate | 0.264 | 0.036 | 0 | 0.036 | 0.133 | 0.025 | 0.083 | 0.183 | >0.9999 |
| $\beta_{PGSxEra}$ | no PGSxYoB covariate | 0.060 | 0.022 | 0 | 0.032 | 0.040 | 0.018 | 0.004 | 0.076 | 0.9854 |

**Supplementary Table 7. Comparison of heritability estimates from GCTA-GREML in the Soviet and post-Soviet eras, as well as the eras further divided by the wave of participation.** The 15-year cutoff was used. Two-tailed Wald test *p*-values are reported. The Bonferroni-corrected significance level was $\alpha = 0.0125$, corresponding to four independent tests, in analyses of the combined wave 1 and wave 2 samples and $\alpha = 6.25\times10^{-3}$, corresponding to eight independent tests, in analyses stratified by wave. *P*-values below the corresponding significance threshold are shown in bold.

| Threshold | Soviet vs. post-Soviet | | | Soviet | post-Soviet |
|---|---|---|---|---|---|
| | All | Wave 1 | Wave 2 | Wave 1 vs. Wave 2 | Wave 1 vs. Wave 2 |
| Stringent | 0.092 | 0.720 | 0.097 | 0.180 | 0.740 |
| Relaxed | $7.5\times10^{-3}$ | 0.980 | 0.020 | 0.560 | 0.850 |

**Supplementary Table 8. Comparison of heritability estimates in the Soviet and post-Soviet eras (age in 1991 - cutoff of 10) and the eras additionally divided by the wave of participation.** Abbreviations: EA, educational attainment; OS, occupational status; BMI, body mass index; University, university degree (on liability scale). Two-tailed Wald test *p*-values are reported. The Bonferroni-corrected significance level was α = 0.0125, corresponding to four independent tests, in analyses of the combined wave 1 and wave 2 samples and α = $6.25\times10^{-3}$, corresponding to eight independent tests, in analyses stratified by wave. *P*-values below the corresponding significance threshold are shown in bold.

| Model | Trait | Soviet vs. post-Soviet | | | Soviet | post-Soviet |
|---|---|---|---|---|---|---|
| | | All | Wave 1 | Wave 2 | Wave 1 vs. Wave 2 | Wave 1 vs. Wave 2 |
| Family-inclusive | EA | **$9.4\times10^{-4}$** | 0.52 | 0.04 | 0.97 | 0.59 |
| Stringent-threshold | EA | 0.044 | 0.36 | 0.041 | 0.69 | 0.48 |
| Relaxed-threshold | EA | 0.012 | 0.92 | 0.74 | 0.20 | 0.67 |
| Family-inclusive | University | 0.011 | 0.57 | 0.051 | 0.47 | 0.61 |
| Stringent-threshold | University | 0.70 | 0.36 | 0.041 | 0.69 | 0.48 |
| Relaxed-threshold | University | 0.22 | 0.65 | 0.39 | 0.47 | 0.64 |
| Family-inclusive | OS | 0.53 | **$5.8\times10^{-4}$** | 0.77 | 0.63 | **$4.3\times10^{-4}$** |
| Stringent-threshold | OS | 0.14 | 0.50 | 0.56 | 0.017 | 0.33 |
| Relaxed-threshold | OS | 0.29 | 0.21 | 0.89 | 0.063 | 0.17 |
| Family-inclusive | Height | **$8.3\times10^{-10}$** | 0.70 | **$1.6\times10^{-7}$** | **$7.7\times10^{-6}$** | 0.89 |
| Stringent-threshold | Height | **$7.5\times10^{-4}$** | 0.78 | 0.010 | 0.011 | 0.84 |
| Relaxed-threshold | Height | **$1.9\times10^{-4}$** | 0.76 | **$1.7\times10^{-3}$** | **$1.4\times10^{-5}$** | 0.85 |
| Family-inclusive | BMI | 0.046 | 0.22 | 0.06 | 0.78 | 0.26 |
| Stringent-threshold | BMI | 0.68 | 0.40 | 0.40 | 0.64 | 0.41 |
| Relaxed-threshold | BMI | 0.22 | 0.28 | 0.35 | 0.64 | 0.33 |

**Supplementary Table 9. Comparison of EA heritability estimated in the stringent-threshold model in the Soviet and post-Soviet eras in the sample stratified by sex.** Two-tailed Wald test *p*-values are reported. The Bonferroni-corrected significance level was α = 0.025, corresponding to two independent tests. *P*-values below the corresponding significance threshold are shown in bold.

| Sex | Cutoff, age in 1991 | *p*-value |
|---|---|---|
| Men | 15 | 0.50 |
| Men | 10 | 0.80 |
| Women | 15 | **0.012** |
| Women | 10 | **0.013** |

**Supplementary Table 10. Comparison of EA heritability estimated in the stringent-threshold model in the Soviet and post-Soviet eras in the sample stratified by type of settlement of residence.** Two-tailed Wald test *p*-values are reported. The Bonferroni-corrected significance level was α = 0.017, corresponding to three independent tests. *P*-values below the corresponding significance threshold are shown in bold.

| Type of settlement | Cutoff, age in 1991 | *p*-value |
|---|---|---|
| Rural | 15 | 0.68 |
| Town | 15 | 0.34 |
| City | 15 | 0.47 |
| Rural | 10 | **0.010** |
| Town | 10 | 048 |
| City | 10 | 0.76 |

**Supplementary Table 11. Comparison of variance explained by PGS from the PGI repository in the post-Soviet and Soviet eras, and the eras additionally divided by the wave of participation.** PGS from the PGI repository. Two-tailed bootstrap *p*-values are reported for $R^2$ comparison. Covariates were not included in the models using inverse normal-transformed (INT) EA because the phenotype had already been adjusted for these covariates before the transformation. The Bonferroni-corrected significance level was $\alpha = 0.0125$, corresponding to four independent tests, in analyses of the combined wave 1 and wave 2 samples and $\alpha = 6.25 \times 10^{-3}$, corresponding to eight independent tests, in analyses stratified by wave. *P*-values below the corresponding significance threshold are shown in bold.

| Trait | $R^2$ | Soviet vs. post-Soviet | | | Soviet | post-Soviet |
|---|---|---|---|---|---|---|
| | | All | Wave 1 | Wave 2 | Wave 1 vs. Wave 2 | Wave 1 vs. Wave 2 |
| EA | Incremental | **<0.002** | **<0.002** | 0.018 | 0.034 | 0.04 |
| EA | Pre-adjusted trait | **0.002** | **<0.002** | 0.03 | 0.024 | **0.004** |
| INT EA | — | 0.044 | **<0.002** | 0.024 | 0.488 | **0.004** |
| OS | Incremental | 0.13 | **<0.002** | 0.108 | 0.744 | **<0.002** |
| OS | Pre-adjusted trait | 0.556 | **<0.002** | 0.798 | 0.902 | **<0.002** |
| INT OS | — | 0.766 | **<0.002** | 0.458 | 0.232 | **<0.002** |
| Height | Incremental | **<0.002** | **0.002** | **<0.002** | **<0.002** | 0.048 |
| Height | Pre-adjusted trait | **<0.002** | **<0.002** | **<0.002** | **<0.002** | 0.026 |
| BMI | Incremental | 0.626 | 0.544 | 0.678 | 0.052 | 0.156 |
| BMI | Pre-adjusted trait | 0.586 | 0.75 | 0.508 | 0.018 | 0.078 |

**Supplementary Table 12. Comparison of variance explained by PGS calculated by this study in the post-Soviet and Soviet eras, and the eras additionally divided by the wave of participation.** Two-tailed bootstrap *p*-values are reported for $R^2$ comparison. The Bonferroni-corrected significance level was $\alpha = 0.0125$, corresponding to four independent tests, in analyses of the combined wave 1 and wave 2 samples and $\alpha = 6.25\times10^{-3}$, corresponding to eight independent tests, in analyses stratified by wave. *P*-values below the corresponding significance threshold are shown in bold.

| Trait | $R^2$ | Soviet vs. post-Soviet | | | Soviet | post-Soviet |
|---|---|---|---|---|---|---|
| | | All | Wave 1 | Wave 2 | Wave 1 vs. Wave 2 | Wave 1 vs. Wave 2 |
| EA | Incremental | 0.070 | 0.088 | 0.058 | 0.024 | 0.196 |
| EA | Pre-adjusted trait | 0.080 | 0.032 | 0.040 | 0.016 | 0.050 |
| OS | Incremental | 0.250 | **<0.002** | 0.374 | 0.798 | **<0.002** |
| OS | Pre-adjusted trait | 0.670 | **<0.002** | 0.888 | 0.626 | **<0.002** |
| Height | Incremental | **<0.002** | 0.614 | **<0.002** | 0.032 | **0.002** |
| Height | Pre-adjusted trait | **<0.002** | 0.690 | **<0.002** | **<0.002** | **0.002** |
| BMI | Incremental | 0.182 | 0.456 | 0.170 | 0.046 | 0.216 |
| BMI | Pre-adjusted trait | 0.874 | 0.498 | 0.954 | 0.316 | 0.210 |

**Supplementary Table 13. Comparison of variance explained by PGS from the PGI repository in the post-Soviet and Soviet eras, and the eras additionally divided by the wave of participation, with the 15-year era cutoff.** Two-tailed bootstrap *p*-values are reported for $R^2$ comparison. The Bonferroni-corrected significance level was $\alpha = 0.0125$, corresponding to four independent tests, in analyses of the combined wave 1 and wave 2 samples and $\alpha = 6.25\times10^{-3}$, corresponding to eight independent tests, in analyses stratified by wave. *P*-values below the corresponding significance threshold are shown in bold.

| Trait | $R^2$ | Soviet vs. post-Soviet | | | Soviet | post-Soviet |
|---|---|---|---|---|---|---|
| | | All | Wave 1 | Wave 2 | Wave 1 vs. Wave 2 | Wave 1 vs. Wave 2 |
| EA | Incremental | 0.06 | 0.012 | 0.242 | **0.004** | 0.138 |
| EA | Pre-adjusted trait | 0.056 | **0.006** | 0.286 | **0.004** | 0.056 |
| OS | Incremental | 0.916 | 0.034 | 0.376 | 0.392 | 0.04 |
| OS | Pre-adjusted trait | 0.316 | 0.012 | 0.54 | 0.672 | 0.01 |
| Height | Incremental | **<0.002** | 0.072 | **<0.002** | **<0.002** | 0.268 |
| Height | Pre-adjusted trait | **<0.002** | 0.012 | **<0.002** | **<0.002** | 0.374 |
| BMI | Incremental | 0.726 | 0.58 | 0.36 | **0.004** | 0.26 |
| BMI | Pre-adjusted trait | 0.796 | 0.732 | 0.426 | **<0.002** | 0.272 |

**Supplementary Table 14. Comparison of EA variance explained by PGS between the Soviet and post-Soviet eras, stratified by sex.** Two-tailed bootstrap *p*-values are reported for $R^2$ comparison. The Bonferroni-corrected significance level was $\alpha = 0.025$, corresponding to two independent tests. *P*-values below the corresponding significance threshold are shown in bold.

| **Sex** | **$R^2$** | ***p*-value** |
|---|---|---:|
| Men | Incremental | 0.026 |
| Men | Pre-adjusted trait | 0.072 |
| Women | Incremental | **0.002** |
| Women | Pre-adjusted trait | **0.006** |

**Supplementary Table 15. Comparison of EA variance explained by PGS between the Soviet and post-Soviet eras, stratified by type of settlement.** Two-tailed bootstrap *p*-values are reported for $R^2$ comparison. The Bonferroni-corrected significance level was $\alpha = 0.017$, corresponding to three independent tests. *P*-values below the corresponding significance threshold are shown in bold.

| **Sex** | **$R^2$** | ***p*-value** |
|---|---|---:|
| Rural | Incremental | **0.002** |
| Rural | Pre-adjusted trait | **0.008** |
| Town | Incremental | 0.318 |
| Town | Pre-adjusted trait | 0.246 |
| City | Incremental | 0.712 |
| City | Pre-adjusted trait | 0.638 |

**Supplementary Table 16. G×Era interaction analysis for EA within the REML framework using the combined sample with the stringent relatedness threshold.** Sample size N = 34,856.

| Relatedness matrix | Proportion of variance explained | SE |
|---|---|---|
| Main | 0.216 | 0.016 |
| Interaction | -0.017 | 0.019 |
| Total | 0.199 | 0.015 |

**Supplementary Table 17.Comparison of variance explained by PGS in weighted subsamples.** Two-tailed bootstrap *p*-values are reported for incremental $R^2$ comparison. Subsamples are based on era and wave of biobank enrolment. The Bonferroni-corrected significance level was $\alpha = 6.25\times10^{-3}$, corresponding to eight independent tests. *P*-values below the corresponding significance threshold are shown in bold.

| Trait | Weighting | Soviet vs. post-Soviet | | Soviet | post-Soviet |
|---|---|---|---|---|---|
| | | Wave 1 | Wave 2 | Wave 1 vs. Wave 2 | Wave 1 vs. Wave 2 |
| EA | No weighting | **<0.002** | 0.014 | 0.018 | 0.064 |
| EA | Inside groups | **<0.002** | 0.008 | 0.092 | 0.082 |
| EA | Overall | **0.002** | **<0.002** | 0.088 | 0.284 |
| OS | No weighting | **<0.002** | 0.114 | 0.742 | **<0.002** |
| OS | Inside groups | **<0.002** | 0.044 | 0.242 | **<0.002** |
| OS | Overall | **<0.002** | 0.324 | 0.730 | **<0.002** |
| Height | No weighting | **<0.002** | **<0.002** | **<0.002** | 0.040 |
| Height | Inside groups | **<0.002** | **<0.002** | **0.002** | 0.092 |
| Height | Overall | **<0.002** | **0.006** | **0.006** | 0.134 |
| BMI | No weighting | 0.546 | 0.632 | 0.060 | 0.156 |
| BMI | Inside groups | 0.34 | 0.872 | 0.040 | 0.038 |
| BMI | Overall | 0.6 | 0.602 | 0.044 | 0.104 |

**Supplementary Table 18. Absolute and relative differences in $h^2_{SNP}$ and $R^2$ for EA between the Soviet and post-Soviet eras estimated by this study and by Rimfeld et al.** Estimates are shown for the 15-year cutoff in both studies. For the current study, $h^2_{SNP}$ estimates from the family-inclusive model and incremental $R^2$ estimates are shown. EA is considered at the scale of the years of education.

| Statistic | Cutoff | Study | Estimates | | | |
|---|---|---|---|---|---|---|
| | | | Soviet | post-Soviet | Δ (difference) | / (proportion) |
| $h^2_{SNP}$ | 15 | Current study | 0.181 [0.171; 0.192] | 0.217 [0.201; 0.234] | 0.036 | 1.20 |
| | | Rimfeld et al. | 0.18 [0.10; 0.26] | 0.37 [0.10; 0.64] | 0.19 | 2.06 |
| | 10 | Current study | 0.183 [0.174; 0.192] | 0.225 [0.202; 0.247] | 0.042 | 1.23 |
| $R^2$ | 15 | Current study | 0.105 [0.100; 0.109] | 0.116 [0.111; 0.122] | 0.011 | 1.11 |
| | | Rimfeld et al. | 0.020 [0.015; 0.025] | 0.032 [0.019; 0.049] | 0.012 | 1.6 |
| | 10 | Current study | 0.107 [0.103; 0.112] | 0.114 [0.108; 0.121] | 0.007 | 1.07 |
| | | Rimfeld et al. | 0.021 [0.018; 0.028] | 0.061 [0.032; 0.102] | 0.040 | 2.90 |

**Supplementary Table 19. Mapping from Education level to Years of education and University degree.**

| Education level | ISCED 2011 mapping | Years of education | University degree |
|---|---|---|---|
| Early childhood education | 0 | 1 | 0 |
| Primary education | 1 | 7 | 0 |
| Lower secondary education | 2 | 10 | 0 |
| Upper secondary education | 2 | 10 | 0 |
| Post-secondary non-tertiary education | 3 | 13 | 0 |
| Short-cycle tertiary education | 4 | 15 | 0 |
| Bachelor's or equivalent level | 6 | 18 | 1 |
| Master's or equivalent level | 7 | 20 | 1 |
| Doctoral or equivalent level | 8 | 22 | 1 |

**Supplementary Table 20. Two-tailed *p*-values for the difference in heritability in the post-Soviet and Soviet eras in the original sample.** The cutoff age refers to the age of individuals in 1991. INT - Rank-Based Inverse Normal Transformed trait; otherwise, the trait was considered at its original scale. GCTA and LDAK indicate a specific method used for estimating $h^2$. The cutoff of 10 or 15 years old divides the cohort into Soviet and post-Soviet eras, and corresponds to the age of an individual in 1991, when Estonia regained its independence. The significance level was set at $\alpha = 0.05$, following the original study.

| Trait | All | | Self-rep. Estonians | |
|---|---|---|---|---|
| | Cutoff 10 | Cutoff 15 | Cutoff 10 | Cutoff 15 |
| GCTA, INT EA | 0.464 | 0.455 | 0.807 | 0.789 |
| GCTA, EA | 0.379 | 0.374 | 0.569 | 0.558 |
| LDAK, EA | - | - | 0.888 | 0.865 |
| LDAK, OS | - | - | 0.119 | 0.127 |

**Supplementary Table 21. Comparison of variance explained by PGS in the post-Soviet and Soviet eras in the original sample.** Two-tailed bootstrap *p*-values are reported for $R^2$ comparison. “EA categories” - a discrete level of EA (ordinal EA); “EA years” - the trait was translated to Years of Education. The cutoff age refers to the age of individuals in 1991. Incremental and pre-adjusted $R^2$ values are reported. The significance level was set at α = 0.05, following the original study.

| Trait | $R^2$ | All | | Self-reported Estonians | |
|---|---|---|---|---|---|
| | | Cutoff 10 | Cutoff 15 | Cutoff 10 | Cutoff 15 |
| EA years | Incremental | **<0.002** | **<0.002** | **0.012** | **<0.002** |
| EA years | Pre-adjusted | **<0.002** | **<0.002** | **0.002** | **<0.002** |
| EA categories | Incremental | **0.028** | **0.002** | 0.14 | **0.022** |
| EA categories | Pre-adjusted | **0.004** | **0.002** | **0.03** | **0.012** |
| OS | Incremental | 0.168 | **0.01** | 0.218 | **0.008** |
| OS | Pre-adjusted | **0.044** | **0.002** | 0.05 | **0.002** |

**Supplementary Table 22. Comparison of EA and OS variance explained by $PGS_{Cog}$ and $PGS_{NonCog}$ in the Soviet and post-Soviet eras.** Two-tailed bootstrap *p*-values are reported for $R^2$ comparison. The Bonferroni-corrected significance level was α = 0.0125, corresponding to four independent tests. *P*-values below the corresponding significance threshold are shown in bold.

| Trait | PGS | $R^2$ | *p*-value |
|---|---|---|---|
| EA | Cog | Incremental | **<0.002** |
| EA | NonCog | Incremental | **0.006** |
| OS | Cog | Incremental | 0.600 |
| OS | NonCog | Incremental | 0.166 |
| EA | Cog | Pre-adjusted trait | **<0.002** |
| EA | NonCog | Pre-adjusted trait | 0.016 |
| OS | Cog | Pre-adjusted trait | 0.782 |
| OS | NonCog | Pre-adjusted trait | 0.334 |

**Supplementary Table 23. Comparison of EA and OS variance explained by PGS based on non-transmitted alleles in the Soviet and post-Soviet eras.** Two-tailed bootstrap *p*-values are reported for $R^2$ comparison. The Bonferroni-corrected significance level was $\alpha = 0.0125$, corresponding to four independent tests.

| **Trait** | $R^2$ | ***p*-value** |
|---|---|---|
| EA | Incremental | 0.596 |
| EA | Pre-adjusted trait | 0.518 |
| OS | Incremental | 0.678 |
| OS | Pre-adjusted trait | 0.466 |
| Height | Incremental | 0.446 |
| Height | Pre-adjusted trait | 0.124 |
| BMI | Incremental | 0.620 |
| BMI | Pre-adjusted trait | 0.442 |

#### Supplementary Figures

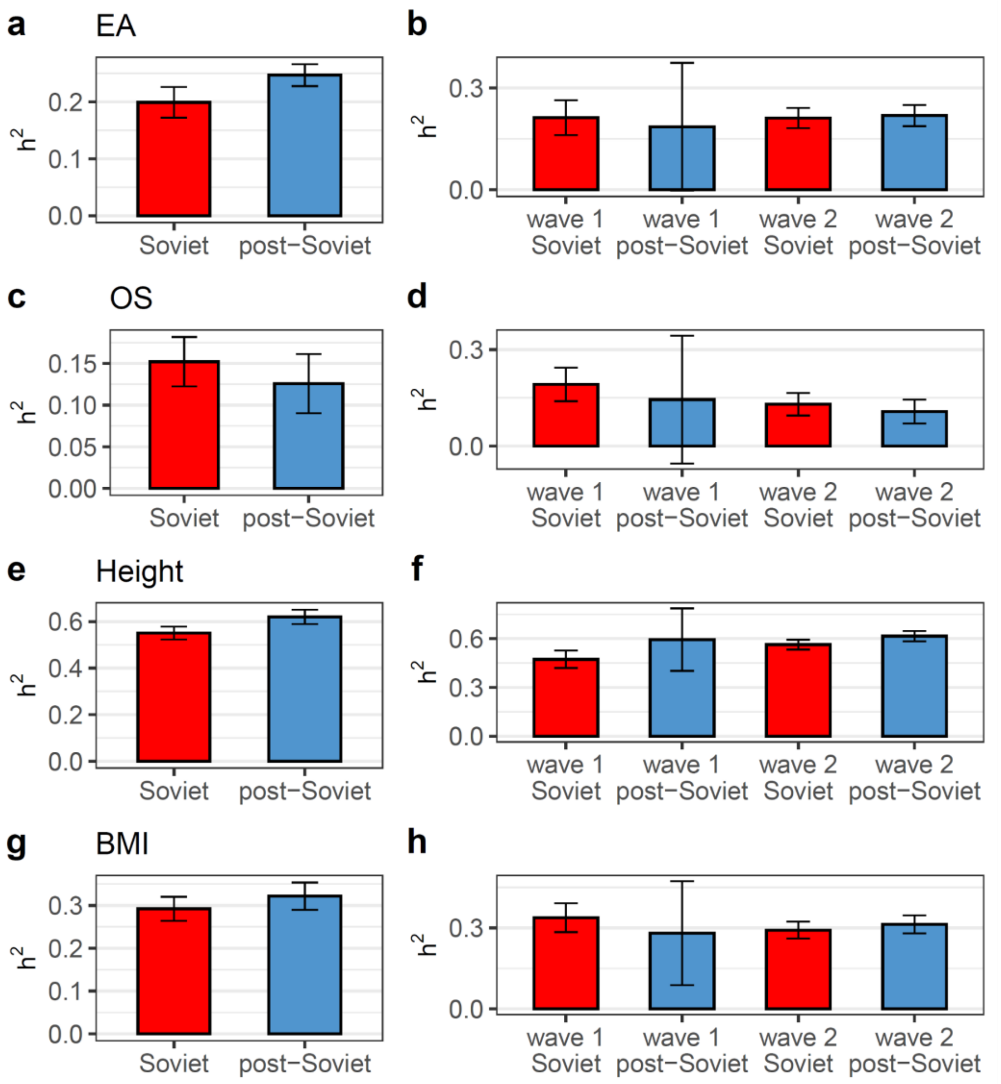

**Supplementary Fig. 1. Trait heritability in the Soviet and post-Soviet eras in the stringent-threshold model.** A 15-year cutoff was used. Trait heritability ($h^2_{SNP}$) is shown in groups divided by era (in the left panels), by era and wave of biobank enrolment (in the right panels) for educational attainment (EA) (**a** and **b**, respectively), occupational status (OS) (**c** and **d**, respectively), height (**e** and **f**, respectively) and body mass index (BMI) (**g** and **h**, respectively). Those subcohorts belonging to the Soviet era are coloured in red, and the post-Soviet era in light blue. The family-inclusive model was used to estimate $h^2_{SNP}$. Error bars correspond to 95% CI.

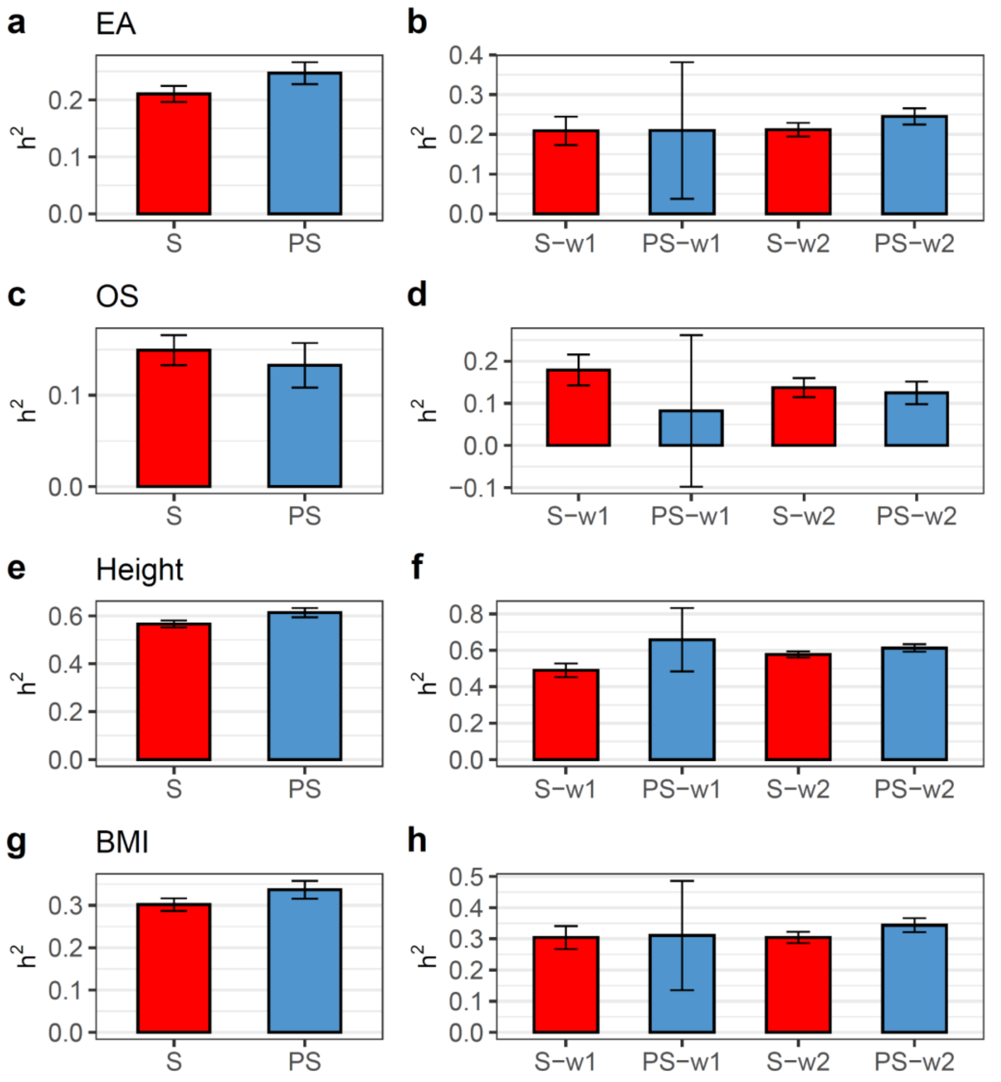

**Supplementary Fig. 2. Trait heritability in the Soviet and post-Soviet eras in the relaxed-threshold model.** A 15-year cutoff was used. Trait heritability ($h^2_{SNP}$) is shown in groups divided by era (in the left panels), by era and wave of biobank enrolment (in the right panels) for educational attainment (EA) (**a** and **b**, respectively), occupational status (OS) (**c** and **d**, respectively), height (**e** and **f**, respectively) and body mass index (BMI) (**g** and **h**, respectively). Those subcohorts belonging to the Soviet era are coloured in red, and the post-Soviet era in light blue. The family-inclusive model was used to estimate $h^2_{SNP}$. Error bars correspond to 95% CI.

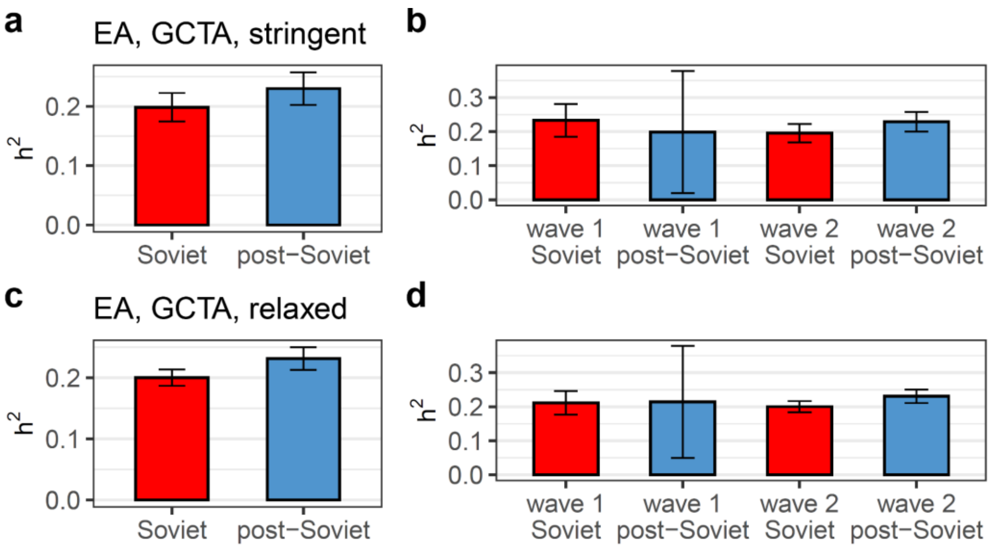

**Supplementary Fig. 3. EA heritability estimates from GREML-GCTA in the Soviet and post-Soviet eras.** A 15-year cutoff and stringent- (**a**, **b**) or relaxed-threshold (**c**, **d**) models were used. Trait heritability ($h^2_{SNP}$) is shown in groups divided by era (in the left panels), by era and wave of biobank enrolment (in the right panels. Those subcohorts belonging to the Soviet era are coloured in red, and the post-Soviet era in light blue. Error bars correspond to 95% CI.

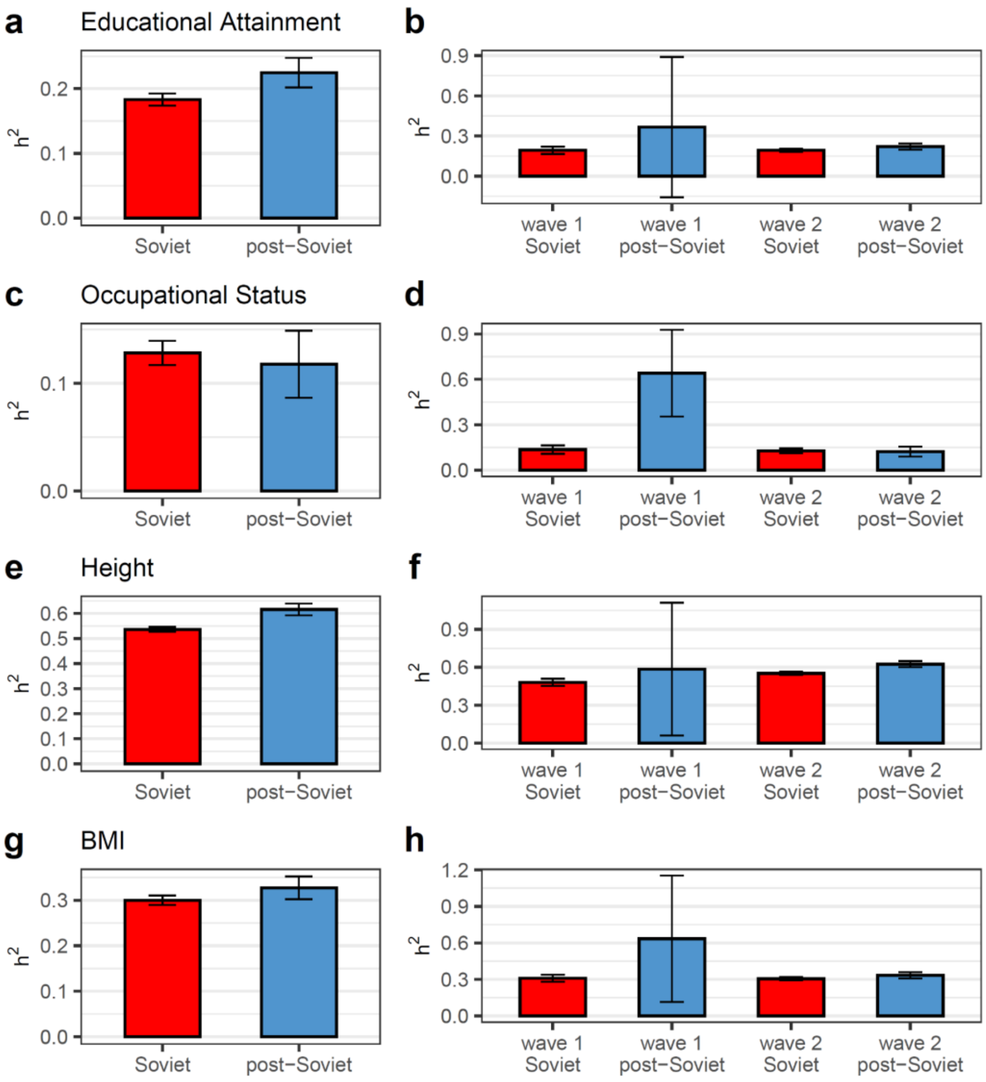

**Supplementary Fig. 4. Trait heritability in the Soviet and post-Soviet eras with a 10-year-old cutoff.** Trait heritability ($h^2_{SNP}$) is shown in groups divided by era (in the left panels), by era and wave of biobank enrolment (in the right panels) for educational attainment (EA) (**a** and **b**, respectively), occupational status (OS) (**c** and **d**, respectively), height (**e** and **f**, respectively) and body mass index (BMI) (**g** and **h**, respectively). Those subcohorts belonging to the Soviet era are coloured in red, and the post-Soviet era in light blue. The family-inclusive model was used to estimate $h^2_{SNP}$. Error bars correspond to 95% CI.

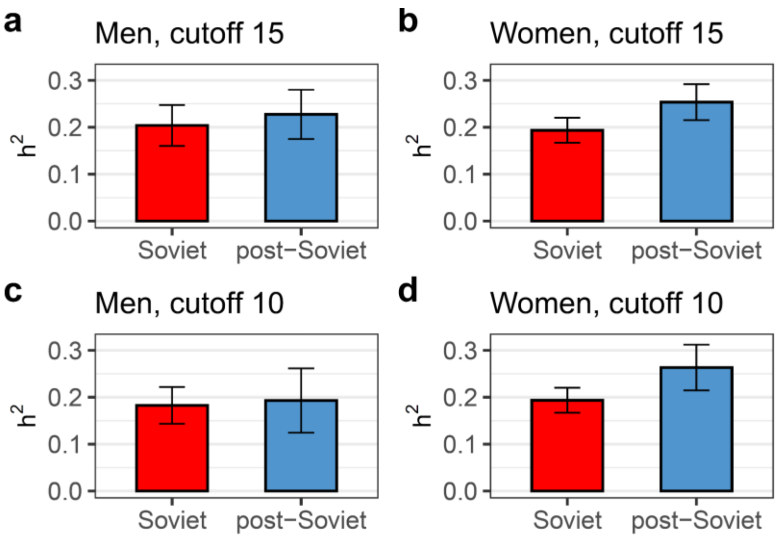

**Supplementary Fig. 5. Educational attainment heritability in the Soviet and post-Soviet eras, stratified by sex.** The heritability was estimated for men (**a**, **c**) and women (**b**, **d**), divided by era, using the stringent-threshold model (relatedness < 0.05) with 15- or 10-year cutoffs. Those subcohorts belonging to the Soviet era are coloured in red, and the post-Soviet era in light blue. Error bars correspond to 95% CI.

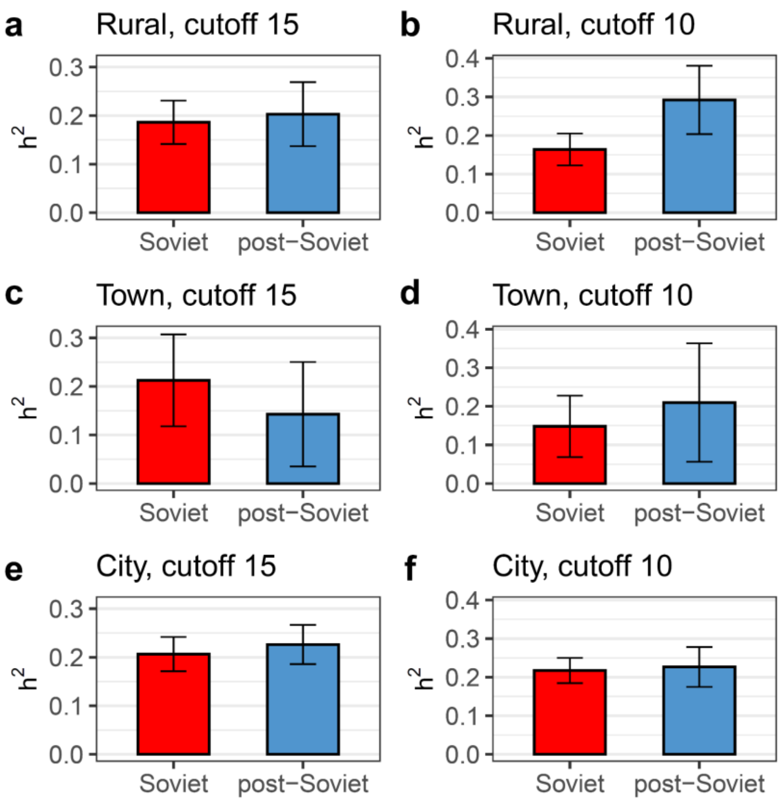

**Supplementary Fig. 6. Educational attainment heritability in the Soviet and post-Soviet eras, stratified by type of settlement of residence.** The heritability was estimated for individuals residing in rural settlements (**a**, **b**), towns (**c**, **d**), or cities (**e**, **f**), divided by era, using the stringent-threshold model (relatedness < 0.05) with 15- (**a**, **c, e**) or 10-year (**b**, **d, f**) cutoffs. Those subcohorts belonging to the Soviet era are coloured in red, and the post-Soviet era in light blue. Error bars correspond to 95% CI.

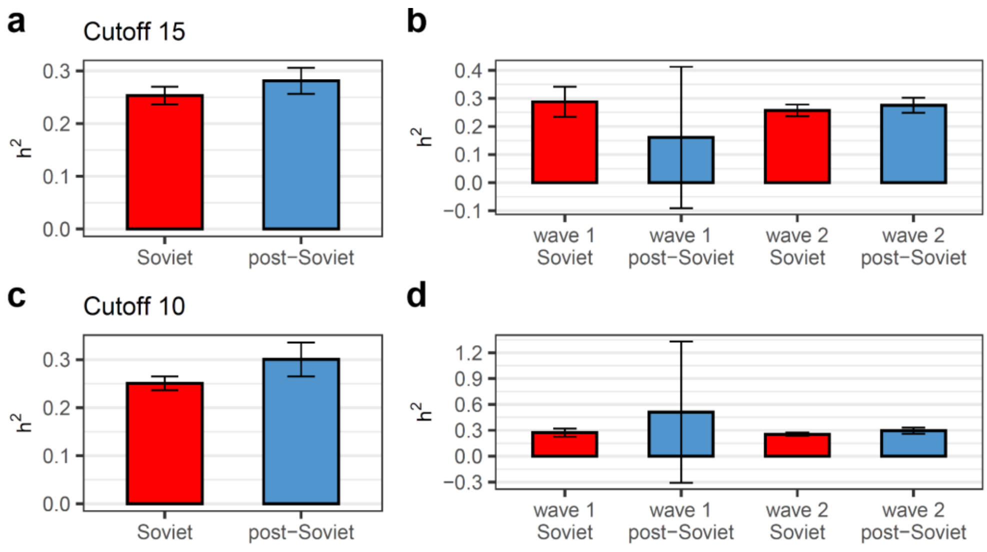

**Supplementary Fig. 7. University degree heritability in the Soviet and post-Soviet eras.** Pairwise comparisons of heritability ($h^2_{SNP}$) estimated with the family-inclusive model were conducted between eras (**a**, **c**), and between waves within each era or between eras within each wave (**b**, **d**) for university degree (binary EA) with a 15-year cutoff (**a** and **b**, respectively), and with a 10-year cutoff (**c** and **d**, respectively). Those subcohorts belonging to the Soviet era are coloured in red, and the post-Soviet era in light blue. Error bars correspond to 95% CI.

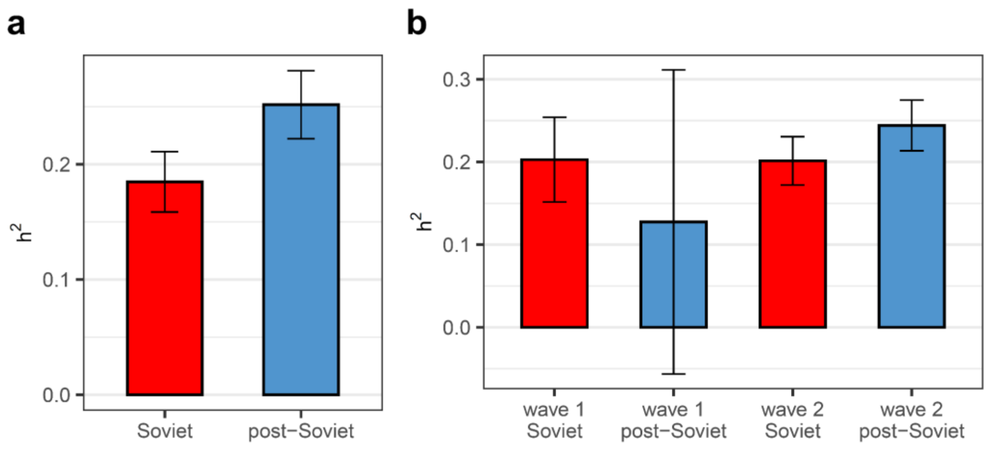

**Supplementary Fig. 8. Heritability of inverse normal-transformed educational attainment in the Soviet and post-Soviet eras.** Pairwise comparisons of heritability ($h^2_{SNP}$) were conducted between eras (**a**), and between waves within each era or between eras within each wave (**b**). $h^2_{SNP}$ was estimated using the stringent-threshold model with a 15-year cutoff. Those subcohorts belonging to the Soviet era are coloured in red, and the post-Soviet era in light blue. Error bars correspond to 95% CI.

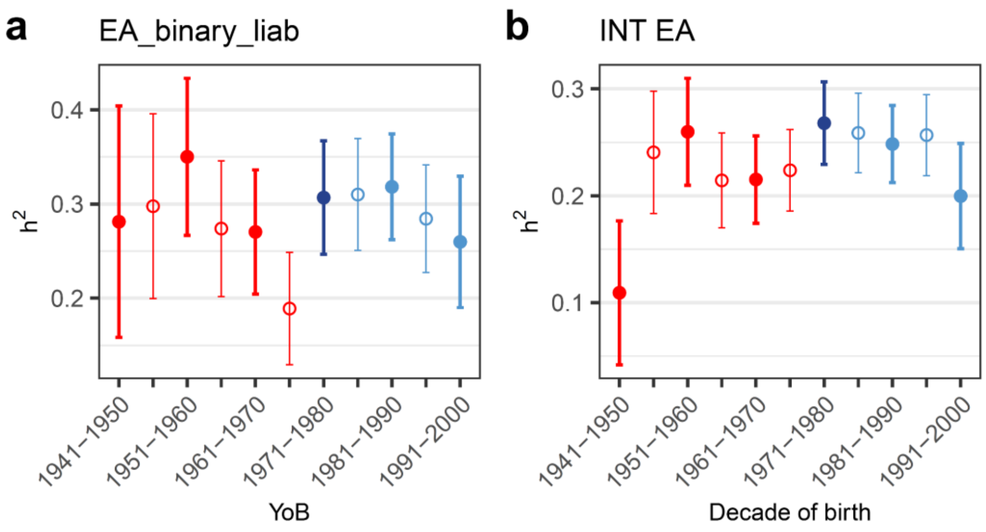

**Supplementary Fig. 9. Heritability across birth year cohorts for university degree (a) and inverse normal-transformed EA (INT EA) (b).** The dataset is split into decade-long birth subcohorts with a 5-year overlap between adjacent subcohorts. Thick error bars and filled circles or thin error bars and empty circles represent non-overlapping subcohorts. Subcohorts from the Soviet era are coloured red, and those from the post-Soviet era are in light blue. A transition group comprising individuals from both the Soviet and post-Soviet eras is shaded in dark blue. $h^2_{SNP}$ was estimated using the stringent-threshold model. Error bars correspond to 95% CI.

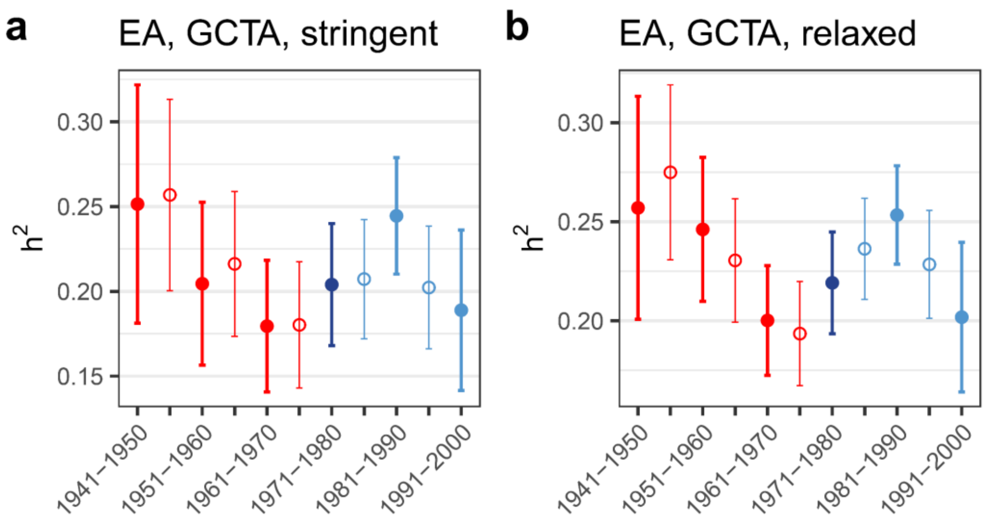

**Supplementary Fig. 10. EA heritability across birth year cohorts estimated with GCTA-GREML.** Stringent- (**a**) or relaxed-threshold (**b**) models were used. The dataset is split into decade-long birth subcohorts with a 5-year overlap between adjacent subcohorts. Thick error bars and filled circles or thin error bars and empty circles represent non-overlapping subcohorts. Subcohorts from the Soviet era are coloured red, and those from the post-Soviet era are in light blue. A transition group comprising individuals from both the Soviet and post-Soviet eras is shaded in dark blue. Error bars correspond to 95% CI.

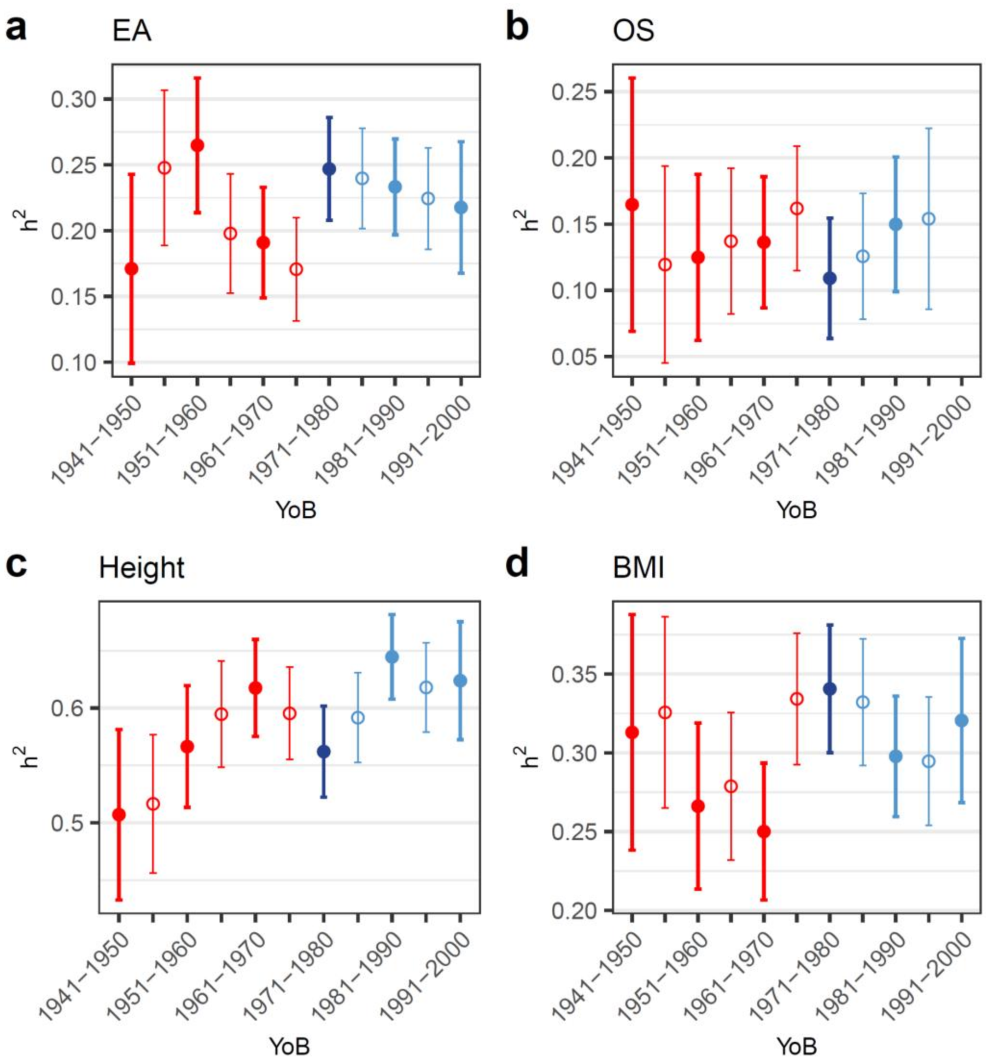

**Supplementary Fig. 11. Heritability across birth year cohorts estimated in the stringent-threshold model.** The dataset is split into decade-long birth subcohorts with a 5-year overlap between adjacent subcohorts. Thick error bars and filled circles or thin error bars and empty circles represent non-overlapping subcohorts. Subcohorts from the Soviet era are coloured red, and those from the post-Soviet era are in light blue. A transition group comprising individuals from both the Soviet and post-Soviet eras is shaded in dark blue. Error bars correspond to 95% CI.

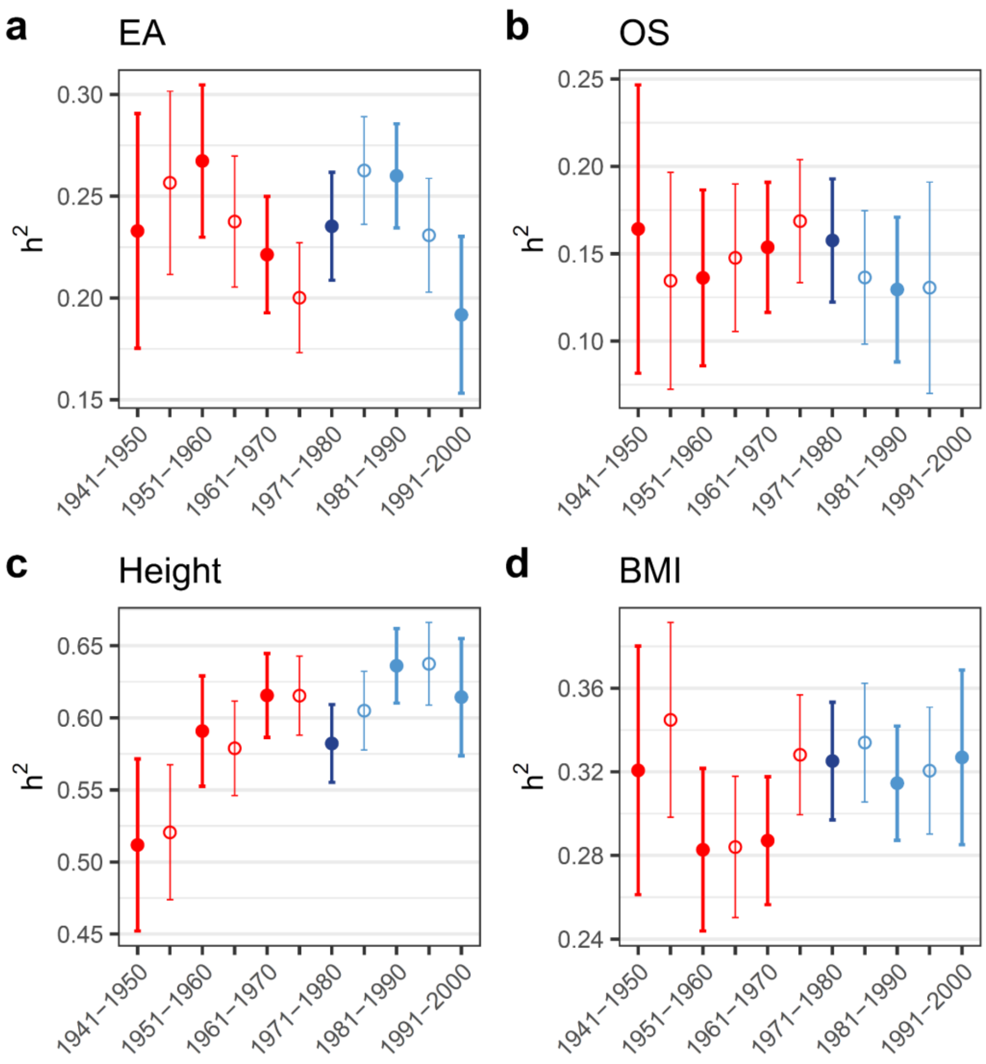

**Supplementary Fig. 12. Heritability across birth year cohorts estimated in the relaxed-threshold model.** The dataset is split into decade-long birth subcohorts with a 5-year overlap between adjacent subcohorts. Thick error bars and filled circles or thin error bars and empty circles represent non-overlapping subcohorts. Subcohorts from the Soviet era are coloured red, and those from the post-Soviet era are in light blue. A transition group comprising individuals from both the Soviet and post-Soviet eras is shaded in dark blue. Error bars correspond to 95% CI.

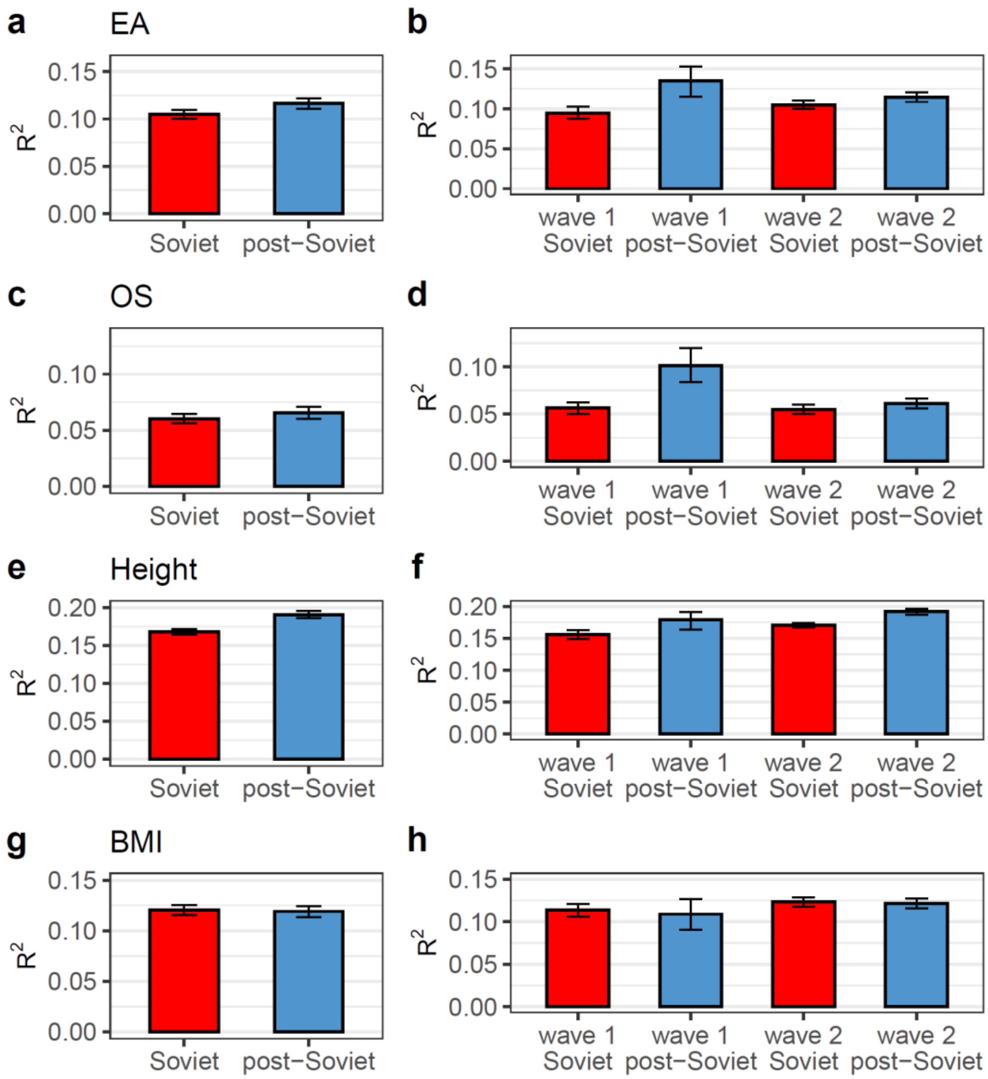

**Supplementary Fig. 13. Trait variance explained by PGS from the PGI repository ($R^2$ for pre-adjusted trait) in the Soviet and post-Soviet eras.** $R^2$ is shown in groups divided by era (in the left panels), or by era and wave of biobank enrolment (in the right panels) for educational attainment (**a, b,** respectively), occupational status (OS) (**c** and **d**, respectively), height (**e** and **f**, respectively), and body mass index (BMI) (**g** and **h**, respectively). Pairwise comparisons of $R^2$ were conducted between eras (**a**, **c**, **e**, **g**), and between waves within each era or between eras within each wave (**b**, **d**, **f**, **h**). Those subcohorts belonging to the Soviet era are coloured in red, and the post-Soviet era in light blue. Error bars correspond to 95% CI.

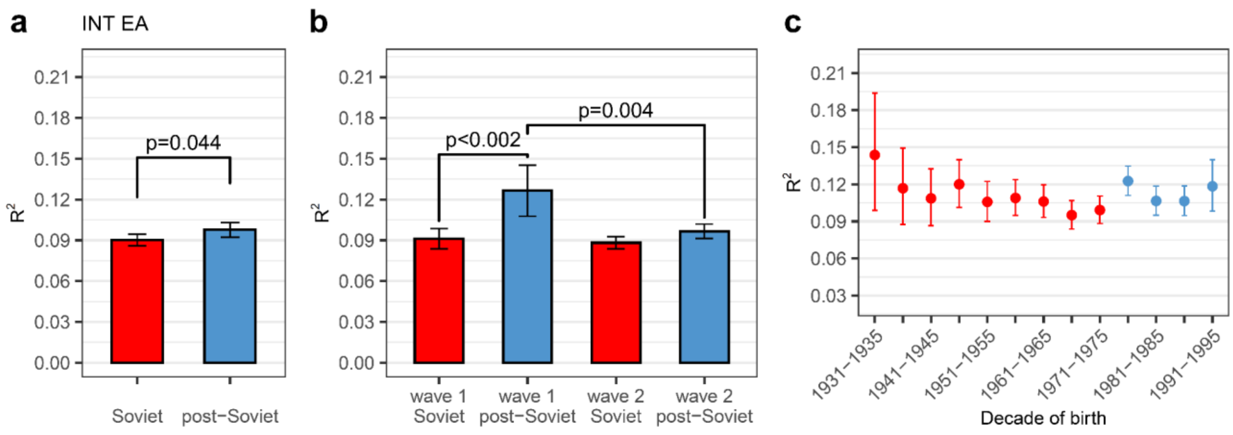

**Supplementary Fig. 14. Variance of inverse normal-transformed educational attainment explained by PGS (incremental $R^2$) in the Soviet and post-Soviet eras.** PGS was from the PGI repository. $R^2$ is shown in groups divided by era (**a**), by era and wave of biobank enrolment (**b**), or by half-decade of birth (**c**). Pairwise comparisons of $R^2$ were conducted between eras (**a**), and between waves within each era or between eras within each wave (**b**). Those subcohorts belonging to the Soviet era are coloured in red, and the post-Soviet era in light blue. Only *p*-values below the significance threshold corrected for multiple testing are shown on panel **b**. The significance thresholds are $\alpha = 0.0125$ (**a**) and $\alpha = 0.006$ (**b**). Error bars correspond to 95% CI.

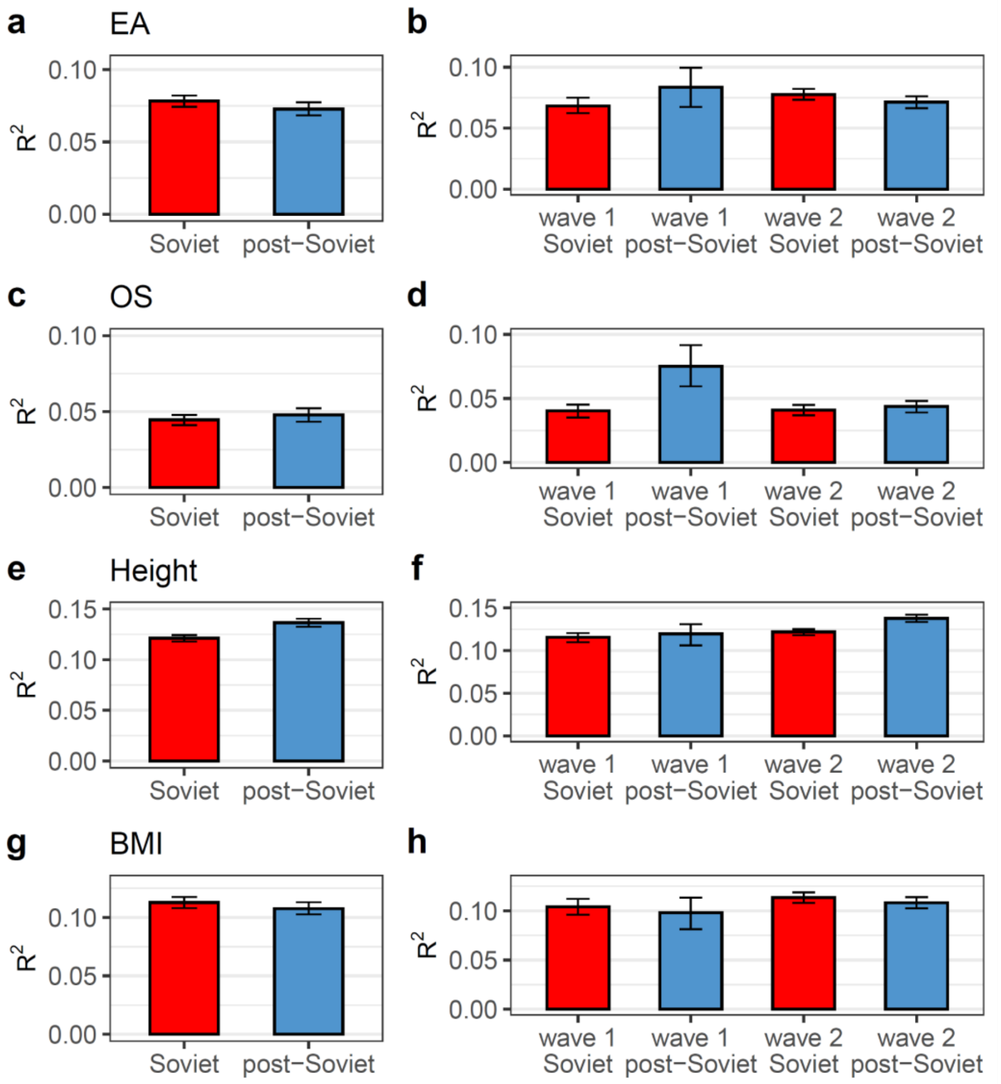

**Supplementary Fig. 15. Trait variance explained by PGS calculated by this study (without the 23andMe cohort included, incremental $R^2$) in the Soviet and post-Soviet eras.** $R^2$ is shown in groups divided by era (in the left panels), or by era and wave of biobank enrolment (in the right panels) for educational attainment (**a, b,** respectively), occupational status (OS) (**c** and **d**, respectively), height (**e** and **f**, respectively), and body mass index (BMI) (**g** and **h**, respectively). Pairwise comparisons of $R^2$ were conducted between eras (**a**, **c**, **e**, **g**), and between waves within each era or between eras within each wave (**b**, **d**, **f**, **h**). Those subcohorts belonging to the Soviet era are coloured in red, and the post-Soviet era in light blue. Error bars correspond to 95% CI.

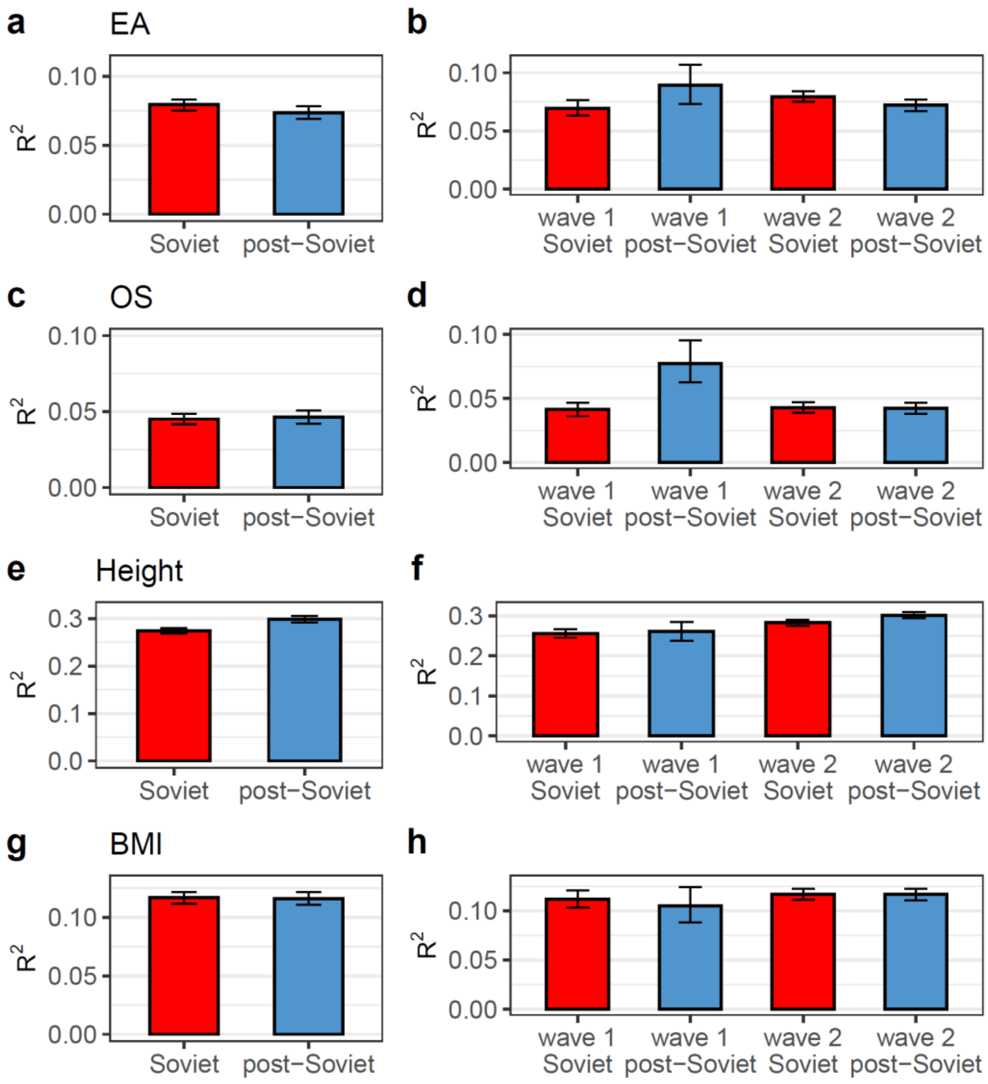

**Supplementary Fig. 16. Trait variance explained by PGS calculated by this study (without the 23andMe cohort included, $R^2$ for pre-adjusted trait) in the Soviet and post-Soviet eras.** $R^2$ is shown in groups divided by era (in the left panels), or by era and wave of biobank enrolment (in the right panels) for educational attainment (**a, b,** respectively), occupational status (OS) (**c** and **d**, respectively), height (**e** and **f**, respectively), and body mass index (BMI) (**g** and **h**, respectively). Pairwise comparisons of $R^2$ were conducted between eras (**a**, **c**, **e**, **g**), and between waves within each era or between eras within each wave (**b**, **d**, **f**, **h**). Those subcohorts belonging to the Soviet era are coloured in red, and the post-Soviet era in light blue. Error bars correspond to 95% CI.

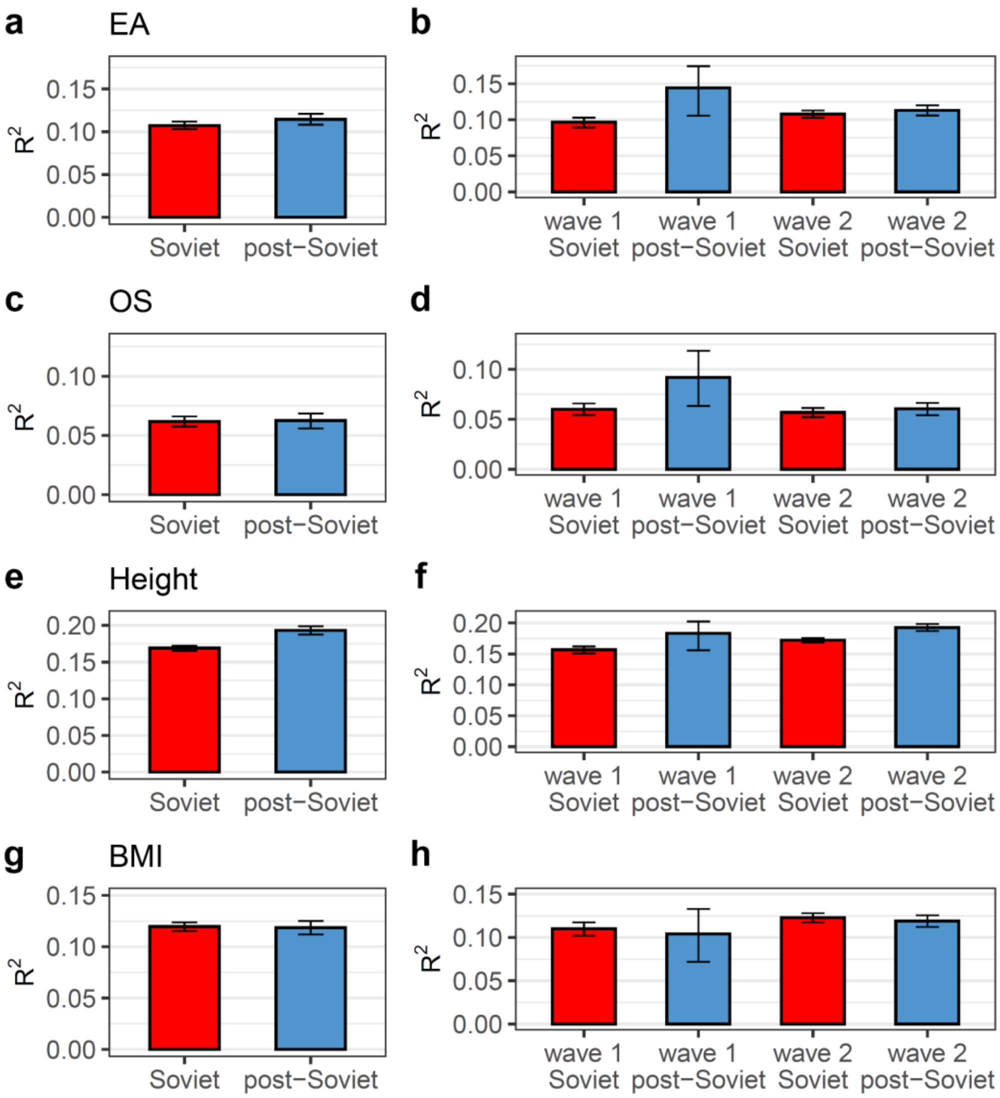

**Supplementary Fig. 17. Trait variance explained by PGS from the PGI repository (incremental $R^2$) in the Soviet and post-Soviet eras, with the 10-year era cutoff.** $R^2$ is shown in groups divided by era (in the left panels), or by era and wave of biobank enrolment (in the right panels) for educational attainment (**a**, **b**, respectively), occupational status (OS) (**c** and **d**, respectively), height (**e** and **f**, respectively), and body mass index (BMI) (**g** and **h**, respectively). Pairwise comparisons of $R^2$ were conducted between eras (a, c, e, g), and between waves within each era or between eras within each wave (**b**, **d**, **f**, **h**). Those subcohorts belonging to the Soviet era are coloured in red, and the post-Soviet era in light blue. Error bars correspond to 95% CI.

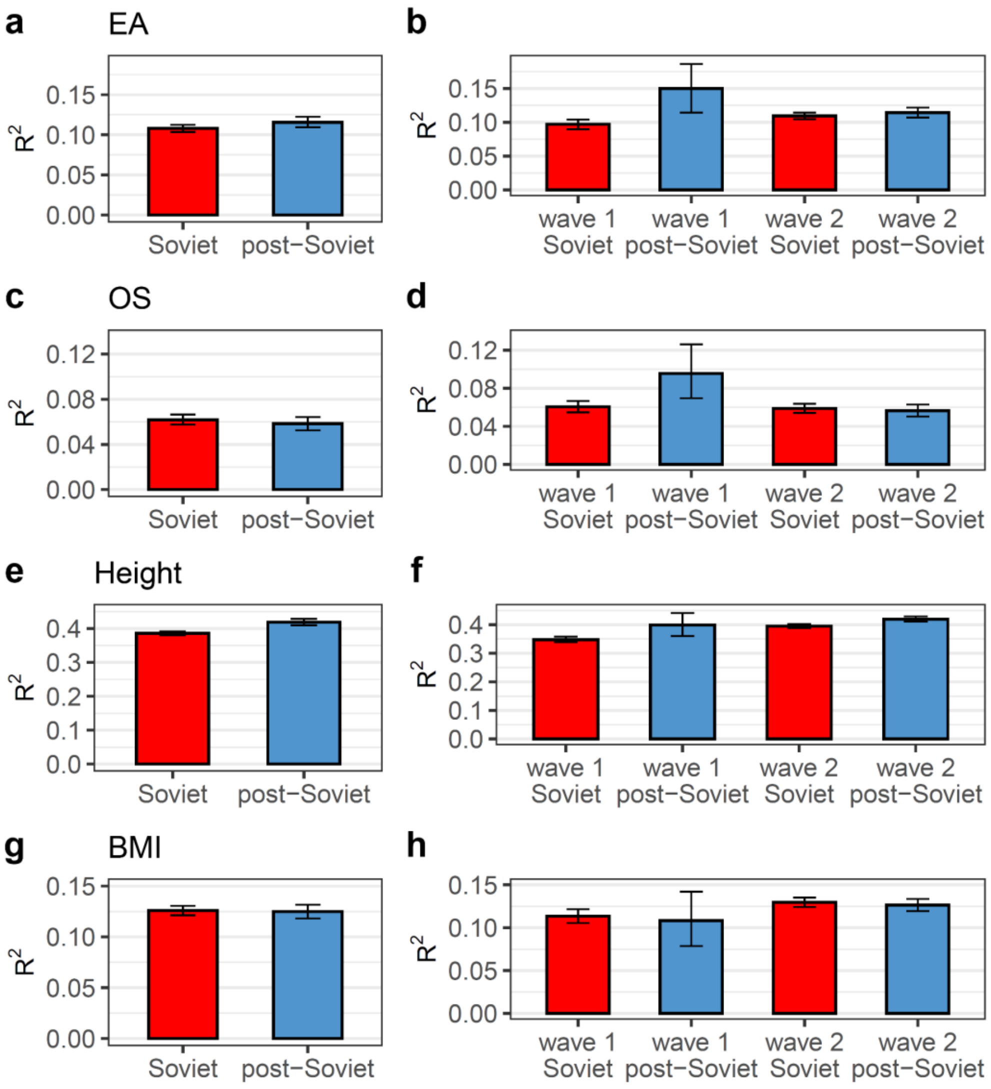

**Supplementary Fig. 18. Trait variance explained by PGS from the PGI repository ($R^2$ for pre-adjusted trait) in the Soviet and post-Soviet eras, with the 10-year era cutoff.** $R^2$ is shown in groups divided by era (in the left panels), or by era and wave of biobank enrolment (in the right panels) for educational attainment (**a**, **b**, respectively), occupational status (OS) (**c** and **d**, respectively), height (**e** and **f**, respectively), and body mass index (BMI) (**g** and **h**, respectively). Pairwise comparisons of $R^2$ were conducted between eras (a, c, e, g), and between waves within each era or between eras within each wave (**b**, **d**, **f**, **h**). Those subcohorts belonging to the Soviet era are coloured in red, and the post-Soviet era in light blue. Error bars correspond to 95% CI.

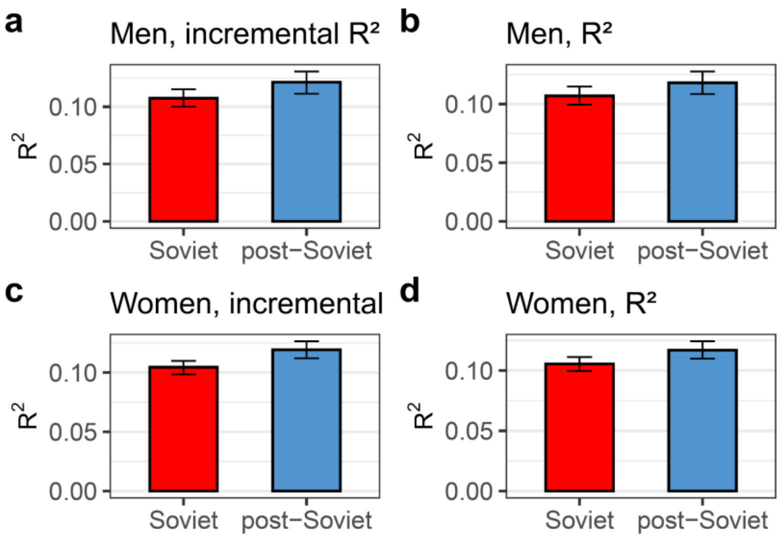

**Supplementary Fig. 19. Educational attainment variance explained by PGS in the Soviet and post-Soviet eras, stratified by sex.** Incremental $R^2$ (**a**, **c**) and $R^2$ for the pre-adjusted trait (**b**, **d**) were calculated for men (**a**, **b**) and women (**c**, **d**), divided by era, using a 15-year cutoff and $PGS_{EA}$ from the PGI repository. Those subcohorts belonging to the Soviet era are coloured in red, and the post-Soviet era in light blue. Error bars correspond to 95% CI.

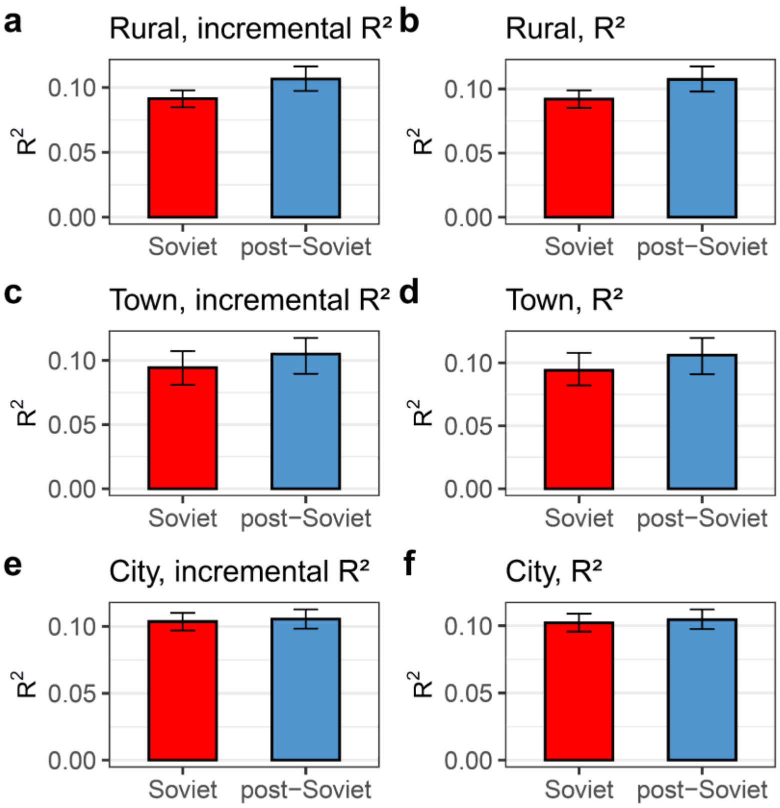

**Supplementary Fig. 20. Educational attainment variance explained by PGS in the Soviet and post-Soviet eras, stratified by type of settlement of residence.** Incremental $R^2$ (**a**, **c, e**) and $R^2$ for the pre-adjusted trait (**b**, **d, f**) were calculated for individuals residing in rural settlements (**a**, **b**), towns (**c**, **d**), and cities (**e**, **f)**, divided by era, using a 15-year cutoff and $PGS_{EA}$ based on summary statistics with the 23andMe cohort included in the meta-analysis. Those subcohorts belonging to the Soviet era are coloured in red, and the post-Soviet era in light blue. Error bars correspond to 95% CI.

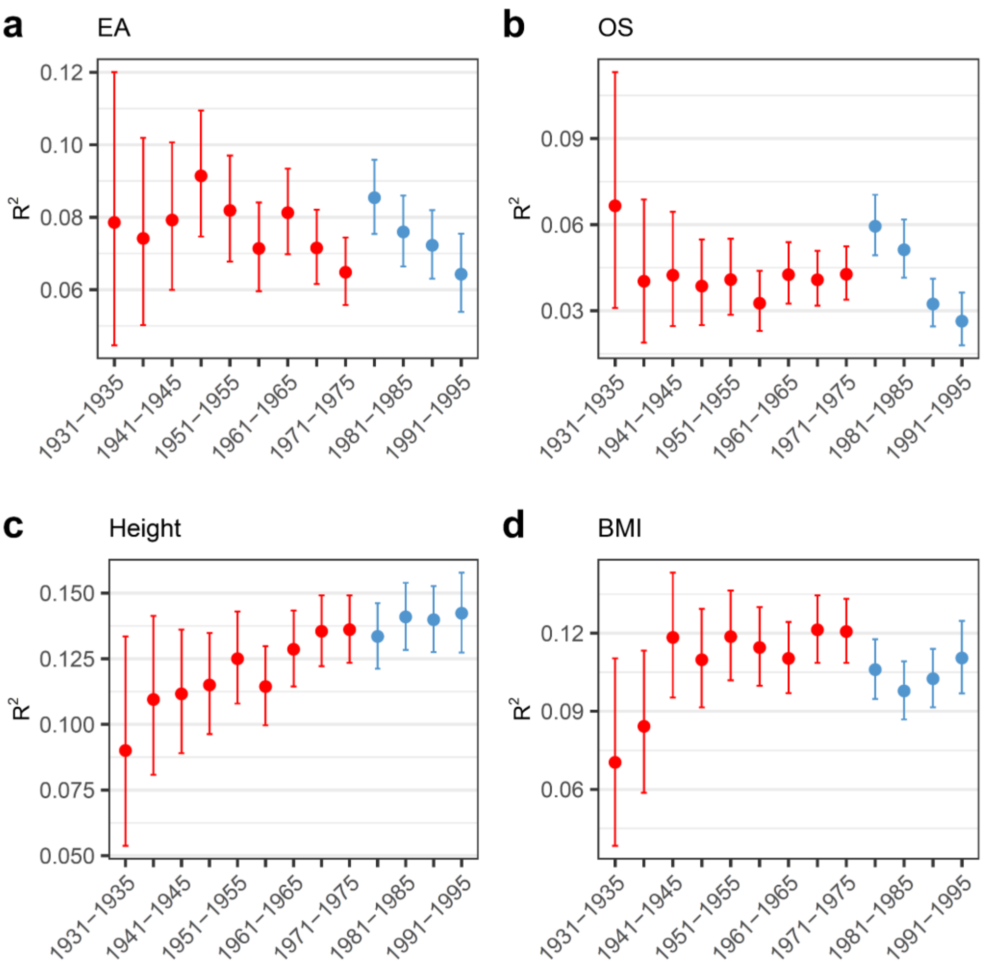

**Supplementary Fig. 21. Trait variance explained by PGS calculated in this study (without the 23andMe cohort included, incremental $R^2$) in the birth cohorts by half-decade.** Those cohorts from the Soviet era are coloured red, and those from the post-Soviet era are coloured light blue. (**a**) height, (**b**) body mass index (BMI). Error bars correspond to 95% CIs.

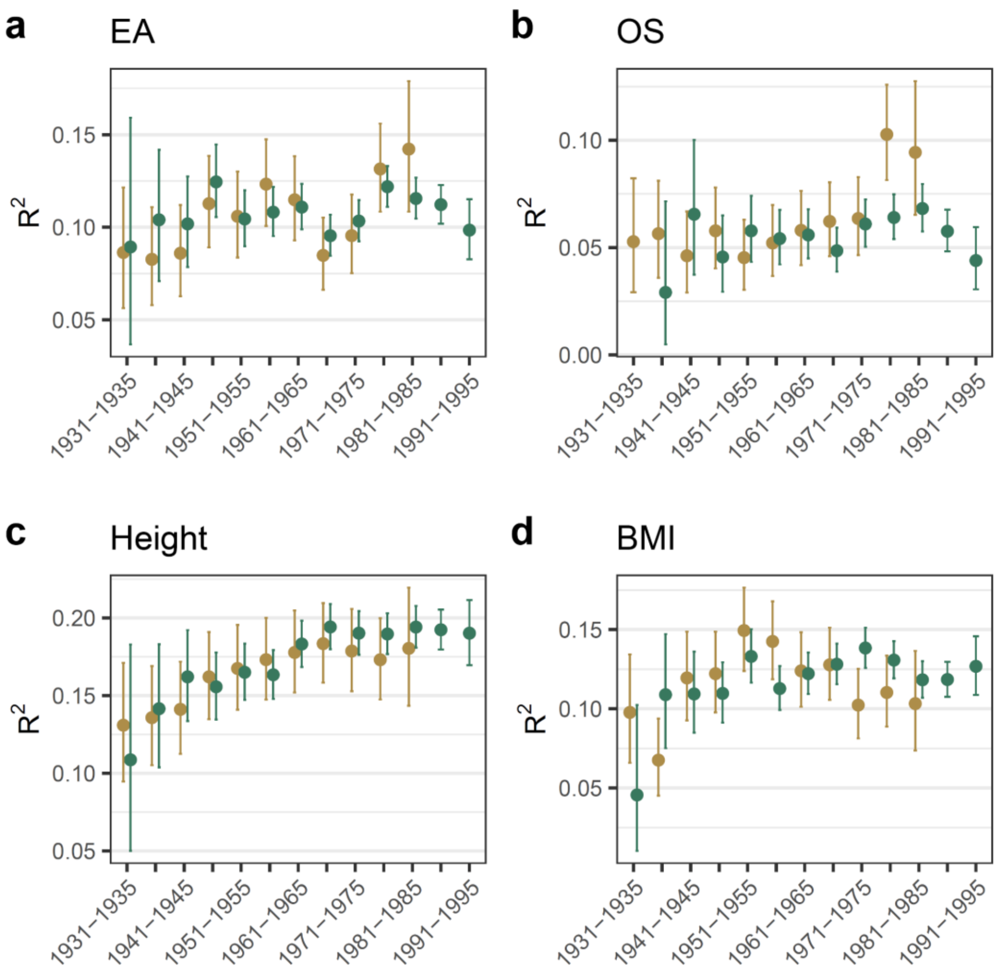

**Supplementary Fig. 22. Trait variance explained by PGS from the PGI repository (incremental $R^2$) in the birth cohorts by half-decade.** Wave 1 is shown with a golden colour, wave 2 is shown with a green colour. (**a**) Educational attainment (EA), (**b**) occupational status (OS), (**c**) height, (**d**) body mass index (BMI). Error bars correspond to 95% CIs.

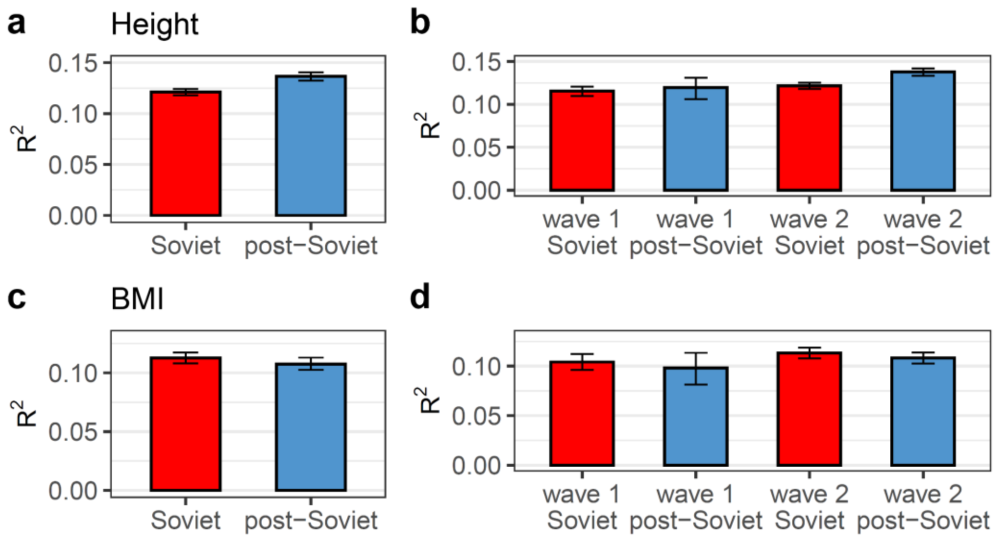

**Supplementary Fig. 23. Trait variance explained by PGS from the PGI repository (incremental $R^2$) for height and BMI in the Soviet and post-Soviet eras.** $R^2$ is shown in groups divided by era (in the left panels) or by era and wave of biobank enrolment (in the right panels) for height (**e** and **f**, respectively) and body mass index (BMI) (**g** and **h**, respectively). Those subcohorts belonging to the Soviet era are coloured in red, and the post-Soviet era in light blue. Error bars correspond to 95% CI.

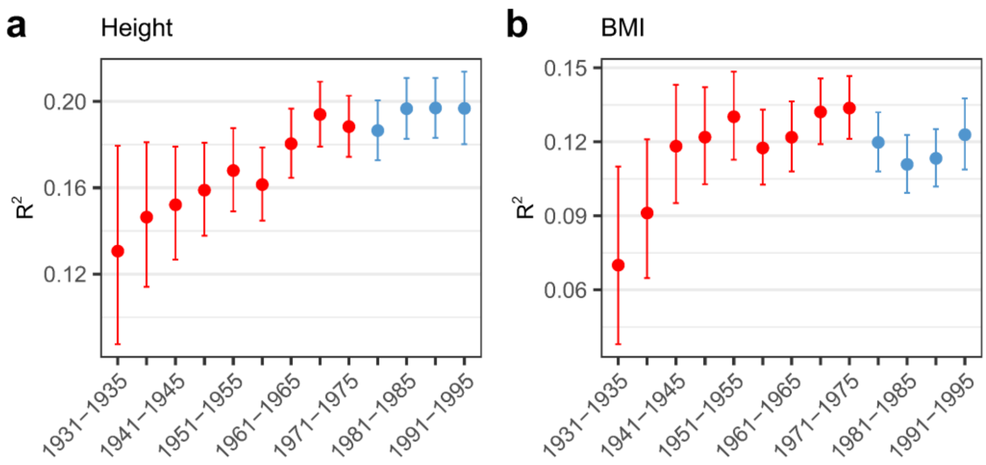

**Supplementary Fig. 24. Trait variance explained by PGS from the PGI repository (incremental $R^2$) for height and BMI in the birth cohorts by half-decade.** Those cohorts from the Soviet era are coloured red, and those from the post-Soviet era are coloured light blue. (**a**) height, (**b**) body mass index (BMI). Error bars correspond to 95% CIs.

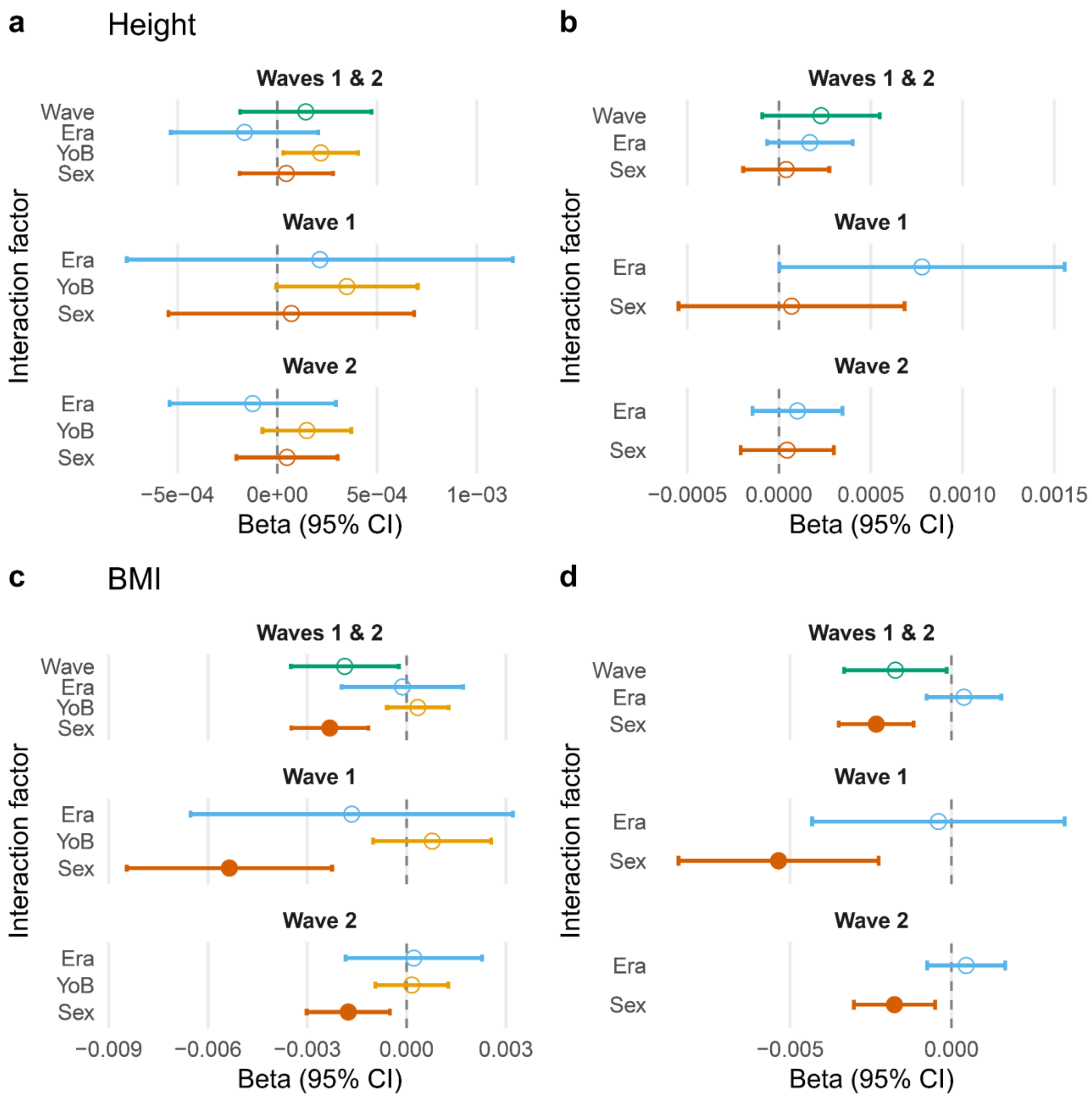

**Supplementary Fig. 25. PGS interaction effects with era and demographic factors across outcomes with 15-year cutoff.** PGSs were obtained from the PGI repository. Linear ordinary least squares regression models for height (a–b) and BMI (c–d) show estimated interaction effects (β, 95% CI) between the corresponding PGS and era, wave, sex, and year of birth (YoB). Models in the left column include the PGS×YoB interaction; models in the right column exclude it. Points represent estimated interaction effects (β) with 95% confidence intervals. Filled circles denote statistically significant effects. The Bonferroni-corrected significance level was $\alpha = 0.0125$, corresponding to four independent tests, in analyses of the combined wave 1 and wave 2 samples (Waves 1 & 2) and $\alpha = 6.25\times10^{-3}$, corresponding to eight independent tests, in analyses stratified by wave.

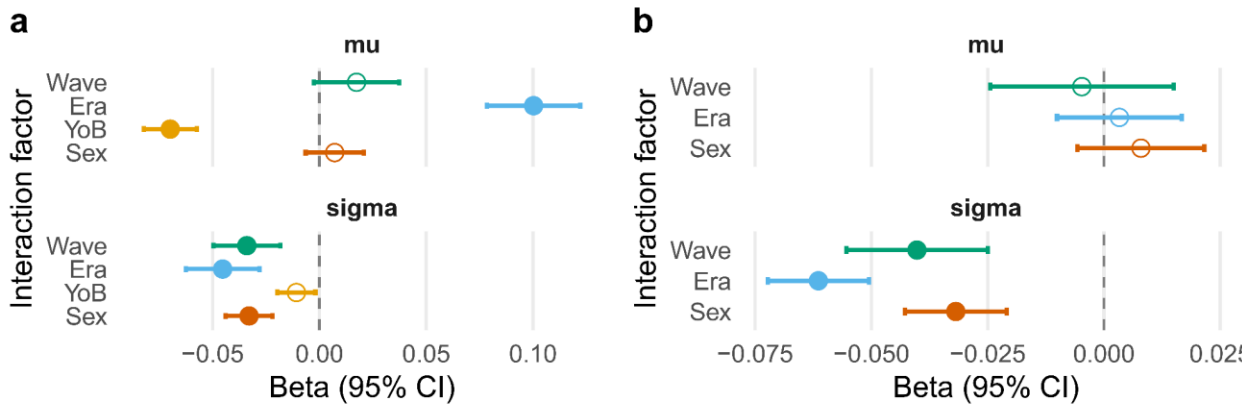

**Supplementary Fig. 26. PGS interaction effects with era and demographic factors across outcomes in a heteroscedastic regression model for EA.** $PGS_{EA}$ was obtained from the PGI repository. Both the mean ("mu") and variance ("sigma") of EA were modelled as functions of the predictors. Estimated interaction effects (β, 95% CI) between $PGS_{EA}$ and era, wave, sex, and year of birth (YoB) are shown. The model in panel **a** includes the PGS×YoB interaction; the model in panel **b** excludes it. Points represent estimated interaction effects (β) with 95% confidence intervals. Filled circles denote statistically significant effects. The Bonferroni-corrected significance level was α = 0.0125, corresponding to four independent tests.

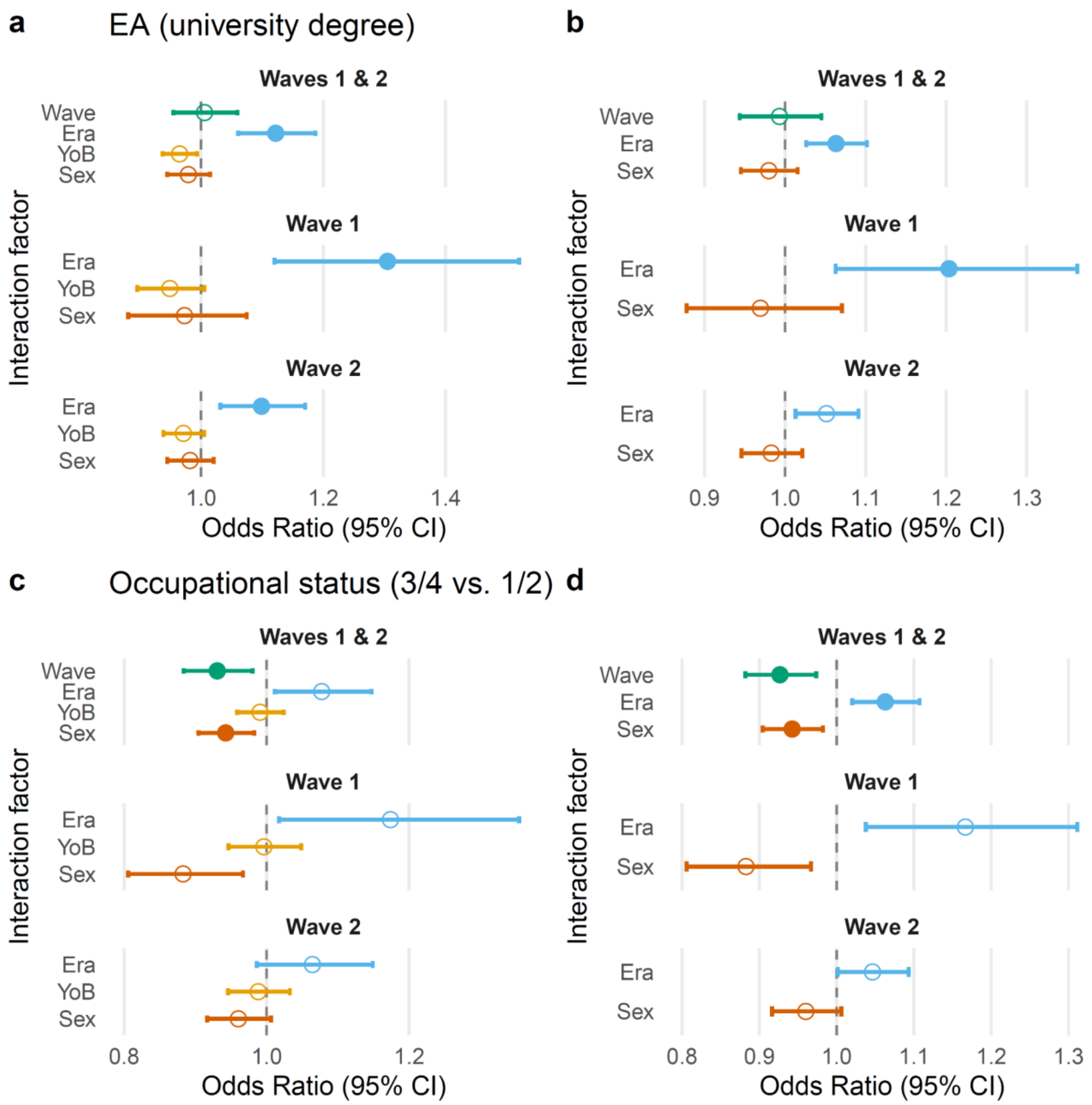

**Supplementary Fig. 27. PGS interaction effects with era and demographic factors across outcomes for binary traits.** $PGS_{EA}$ was obtained from the PGI repository. Logistic regression models for binary EA (university degree) (a–b) and binary occupational status (c–d) show estimated interaction effects (odds ratio, OR, 95% CI) between $PGS_{EA}$ and era, wave, sex, and year of birth (YoB). Models in the left column include the PGS×YoB interaction; models in the right column exclude it. Points represent estimated interaction effects (OR) with 95% confidence intervals. Filled circles denote statistically significant effects. The Bonferroni-corrected significance level was $\alpha = 0.0125$, corresponding to four independent tests, in analyses of the combined wave 1 and wave 2 samples (Waves 1 & 2) and $\alpha = 6.25\times10^{-3}$, corresponding to eight independent tests, in analyses stratified by wave.

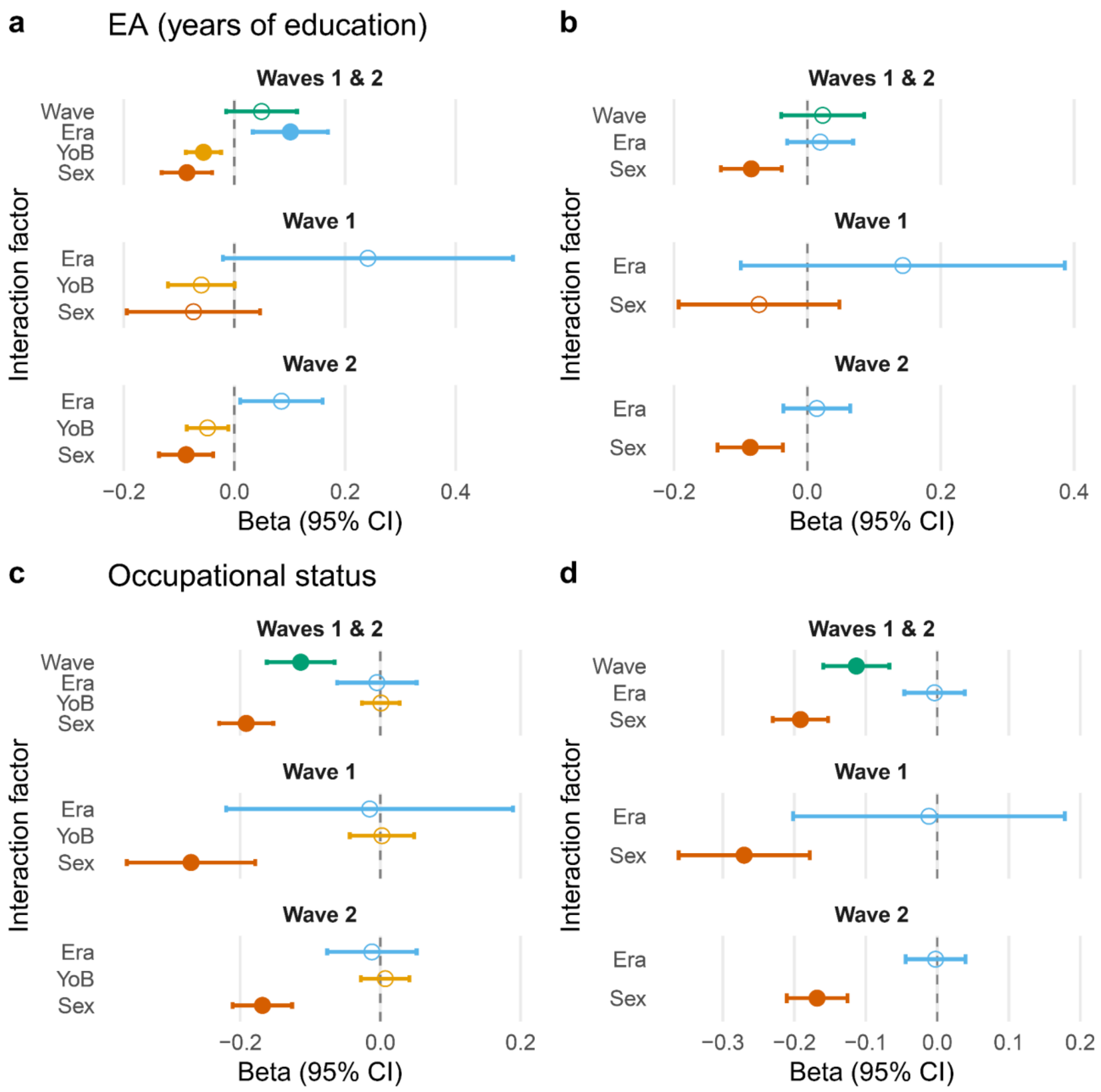

**Supplementary Fig. 28. PGS interaction effects with era and demographic factors across outcomes with 10-year cutoff.** $PGS_{EA}$ was obtained from the PGI repository. Linear ordinary least squares regression models for EA (years of education) (**a**–**b**) and occupational status (**c**–**d**) show estimated interaction effects (β, 95% CI) between $PGS_{EA}$ and era, wave, sex, and year of birth (YoB). Models in the left column include the PGS×YoB interaction; models in the right column exclude it. Points represent estimated interaction effects (β) with 95% confidence intervals. Filled circles denote statistically significant effects. The Bonferroni-corrected significance level was $\alpha = 0.0125$, corresponding to four independent tests, in analyses of the combined wave 1 and wave 2 samples (Waves 1 & 2) and $\alpha = 6.25\times10^{-3}$, corresponding to eight independent tests, in analyses stratified by wave.

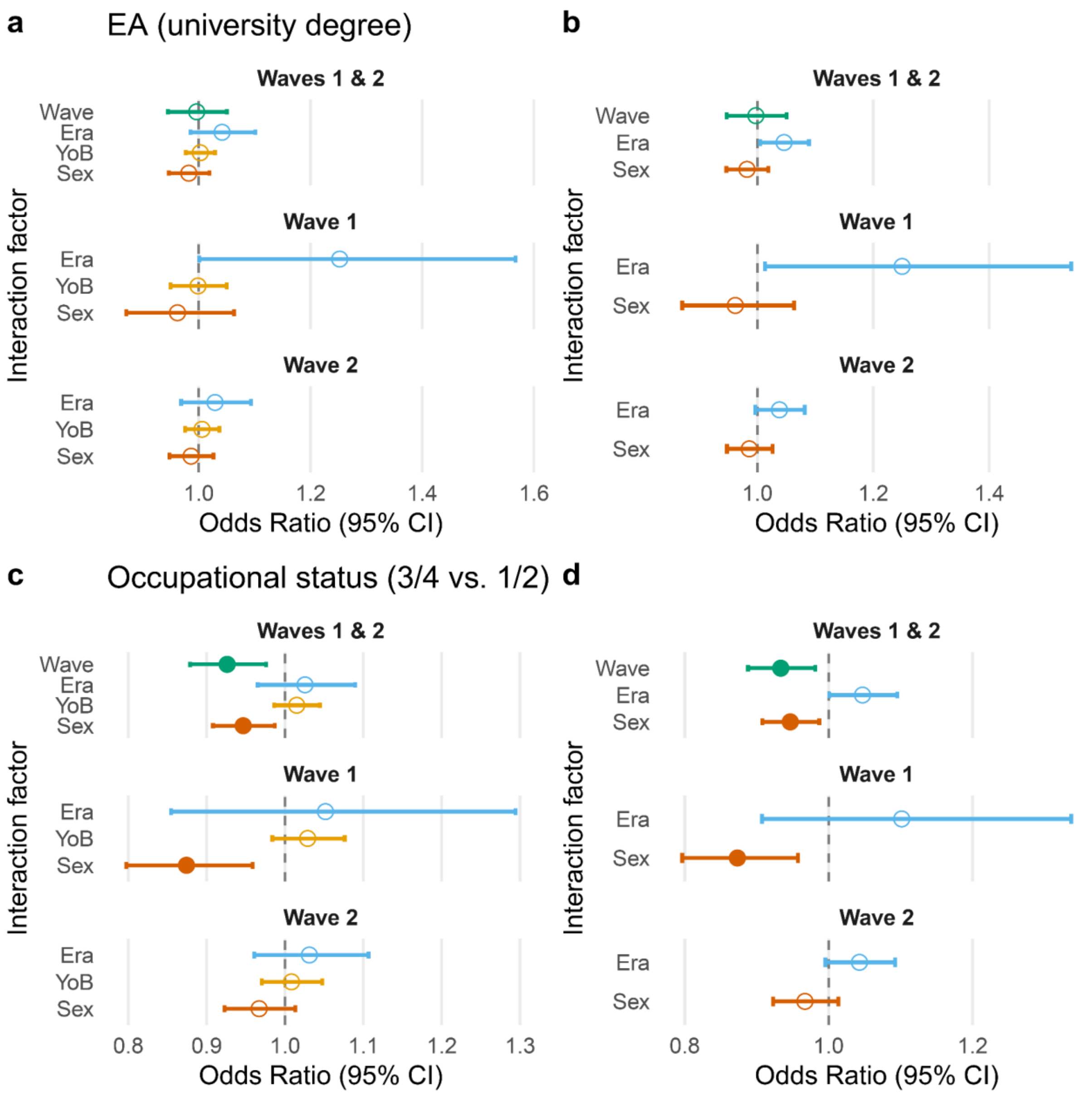

**Supplementary Fig. 29. PGS interaction effects with era and demographic factors across outcomes for binary traits with a 10-year cutoff.** $PGS_{EA}$ was obtained from the PGI repository. Linear regression models for years of education (**a**, **b**) and occupational status (**c**, **d**) show estimated interaction effects (β, 95% CI) between $PGS_{EA}$ and era, wave, sex, and year of birth (YoB). Models in the left column include the PGS×YoB interaction; models in the right column exclude it. Points represent estimated interaction effects (β) with 95% confidence intervals, indicating how the association between $PGS_{EA}$ and the phenotype differs by wave, era, YoB, or sex. Filled circles denote statistically significant effects. The Bonferroni-corrected significance level was α = 0.0125, corresponding to four independent tests, in analyses of the combined wave 1 and wave 2 samples (Waves 1 & 2) and $\alpha = 6.25\times10^{-3}$, corresponding to eight independent tests, in analyses stratified by wave.

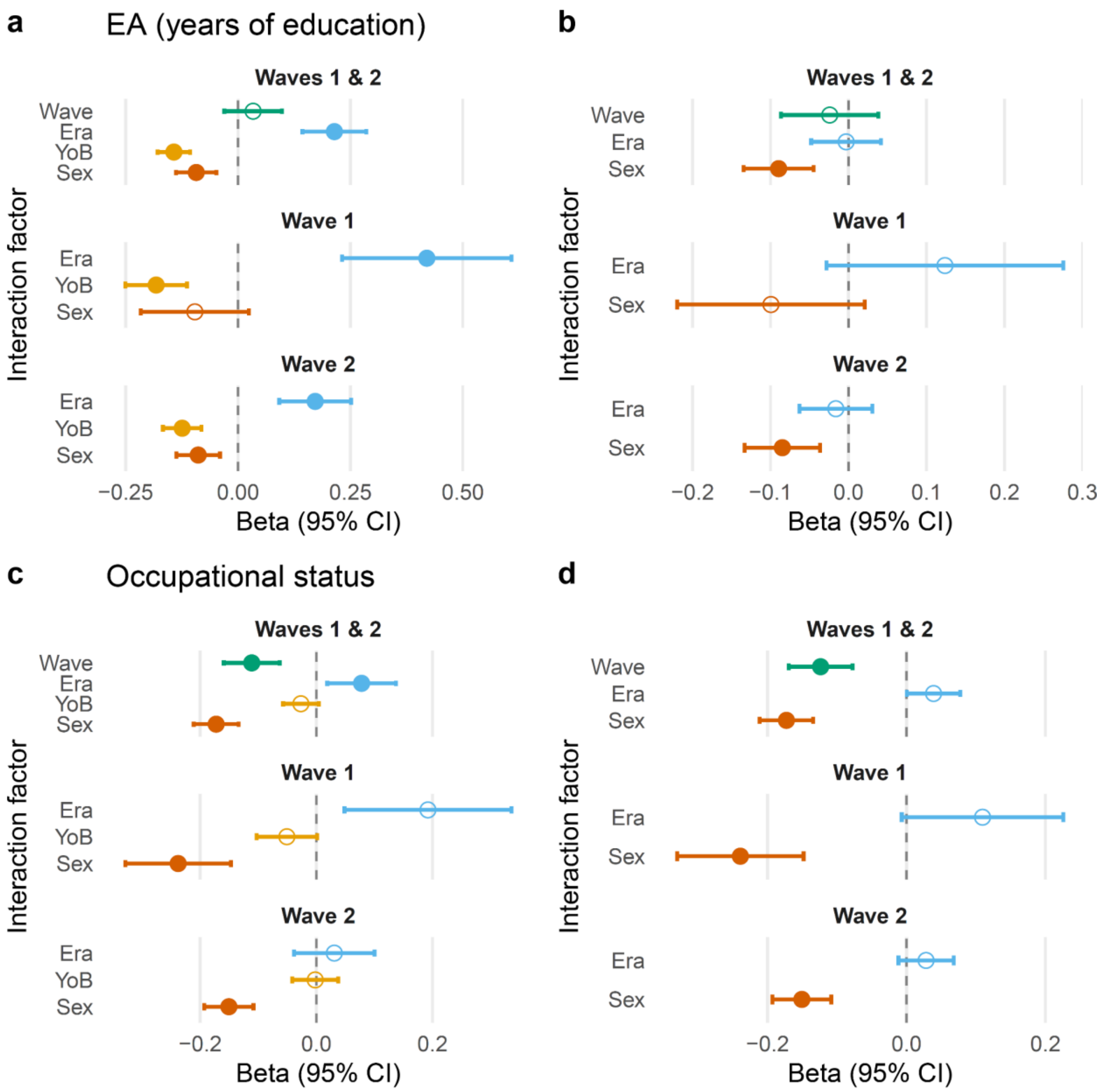

**Supplementary Fig. 30. PGS interaction effects with era and demographic factors across outcomes with $PGS_{EA}$ calculated by this study (without the 23andMe cohort included).** A 15-year cutoff was used. Linear ordinary least squares regression models for EA (years of education) (**a**–**b**) and occupational status (**c**–**d**) show estimated interaction effects (β, 95% CI) between $PGS_{EA}$ and era, wave, sex, and year of birth (YoB). $PGS_{EA}$ was obtained from the PGI repository. Models in the left column include the PGS×YoB interaction; models in the right column exclude it. Points represent estimated interaction effects (β) with 95% confidence intervals. Filled circles denote statistically significant effects. The Bonferroni-corrected significance level was $\alpha = 0.0125$, corresponding to four independent tests, in analyses of the combined wave 1 and wave 2 samples (Waves 1 & 2) and $\alpha = 6.25\times10^{-3}$, corresponding to eight independent tests, in analyses stratified by wave.

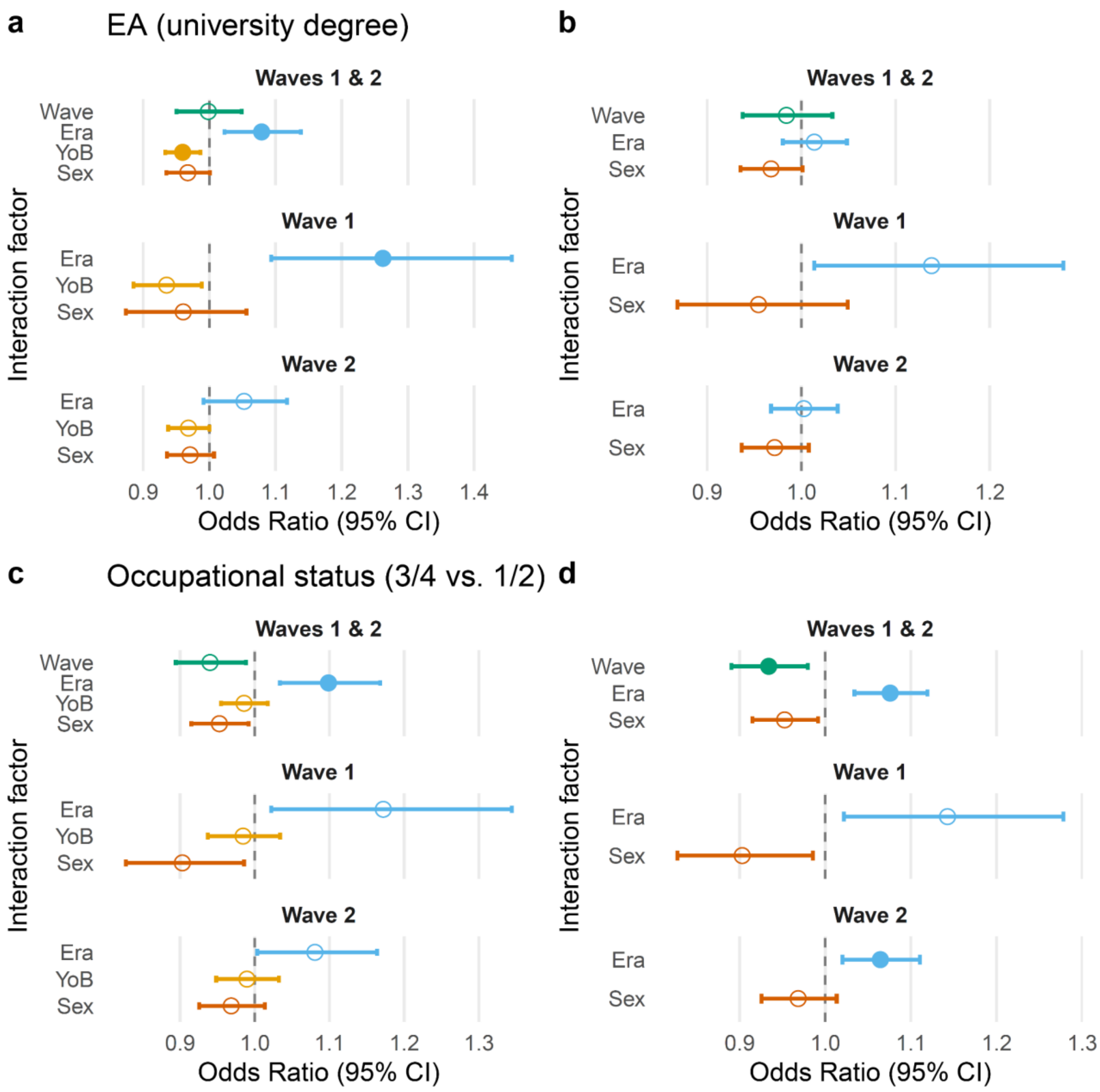

**Supplementary Fig. 31. PGS interaction effects with era and demographic factors across outcomes for binary traits with PGS$_{EA}$ calculated by this study (without the 23andMe cohort included).** A 15-year cutoff was used. Logistic regression models for binary EA (university degree) (a–b) and binary occupational status (c–d) show estimated interaction effects (odds ratio, OR, 95% CI) between PGS$_{EA}$ and era, wave, sex, and year of birth (YoB). Models in the left column include the PGS×YoB interaction; models in the right column exclude it. Points represent estimated interaction effects (OR) with 95% confidence intervals. Filled circles denote statistically significant effects. The Bonferroni-corrected significance level was $\alpha = 0.0125$, corresponding to four independent tests, in analyses of the combined wave 1 and wave 2 samples (Waves 1 & 2) and $\alpha = 6.25\times10^{-3}$, corresponding to eight independent tests, in analyses stratified by wave.

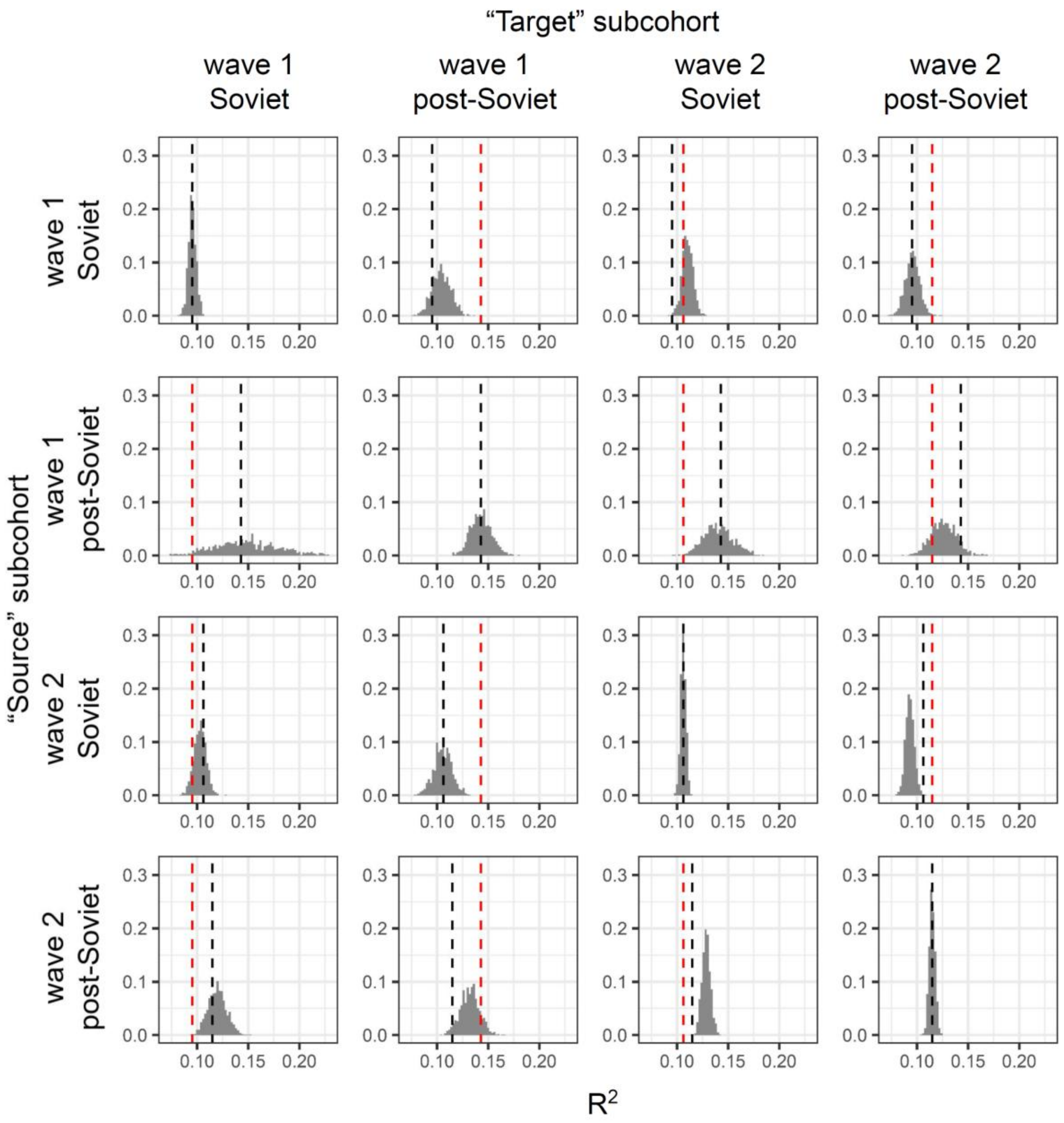

**Supplementary Fig. 32. Distribution of variance explained ($R^2$) by $PGS_{EA}$ in subsamples generated to match the joint sex-EA distribution of a target subcohort.** Subsamples were drawn with replacement from a "source" subcohort defined by recruitment wave and era, matching the joint distribution of EA and sex. Rows indicate the source subcohort from which individuals were sampled, columns indicate the target subcohort whose EA×sex distribution was used for matching. For each source–target pair, 1,000 resampled subsamples were generated, and $R^2$ was calculated as the squared Pearson correlation between the pre-adjusted EA residuals (adjusted for sex, age, sex×age, age², and 40 PCs) and $PGS_{EA}$. Histograms show the distribution of $R^2$ across resamples. Black lines indicate the original $R^2$ in the source subcohort, red lines indicate the original $R^2$ in the target subcohort. Diagonal panels represent cases where source and target eras are identical. $PGS_{EA}$ was obtained from the PGI repository.

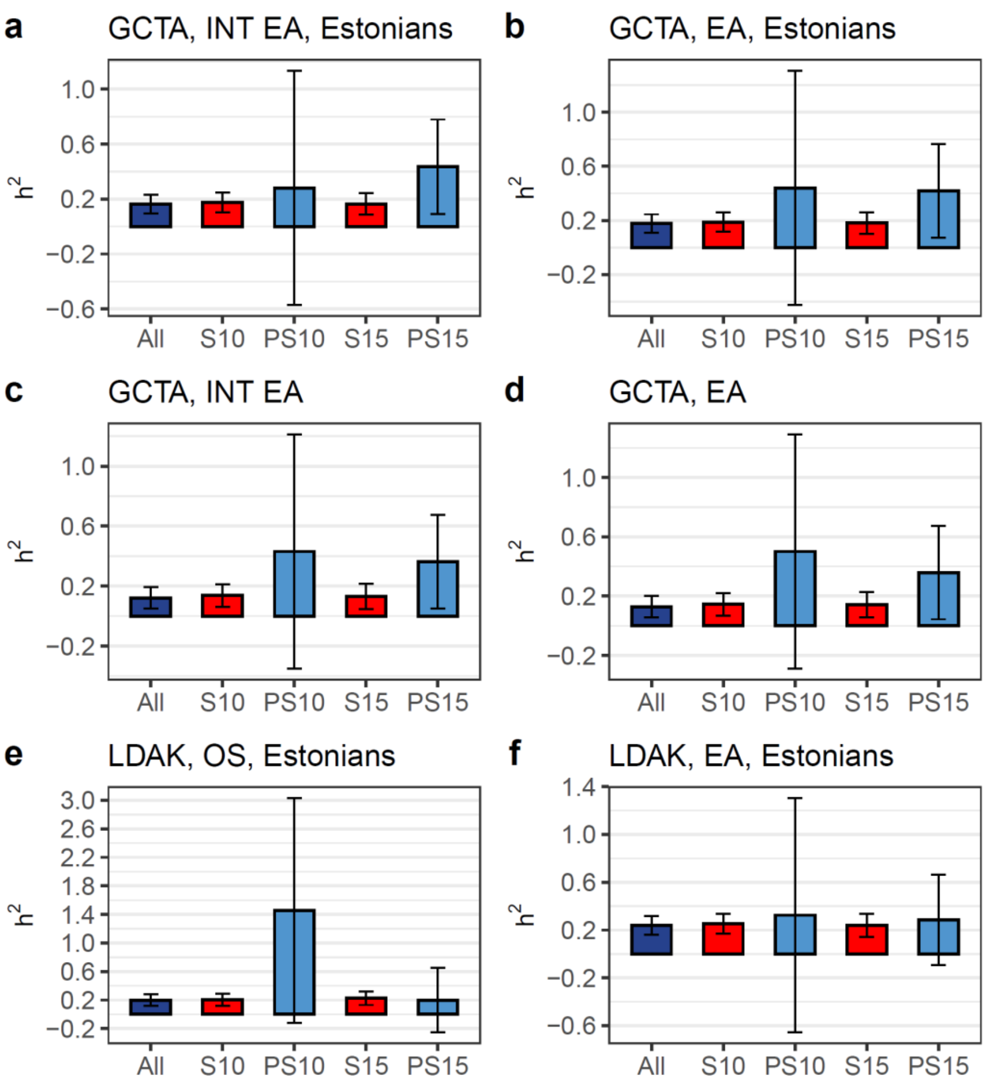

**Supplementary Fig. 33. Heritability in the Soviet and post-Soviet eras in the original sample.** S - Soviet, PS - post-Soviet eras by era; index 10 or 15 means cutoff between the eras - the age of individuals in 1991. INT - Rank-Based Inverse Normal Transformed trait; otherwise, the trait was considered at its original discrete scale of the level of education. “Estonians” in the title means that only self-reported Estonian individuals were used in the analysis. GCTA and LDAK indicate a specific method used for estimating $h^2$. Error bars correspond to 95% CI.

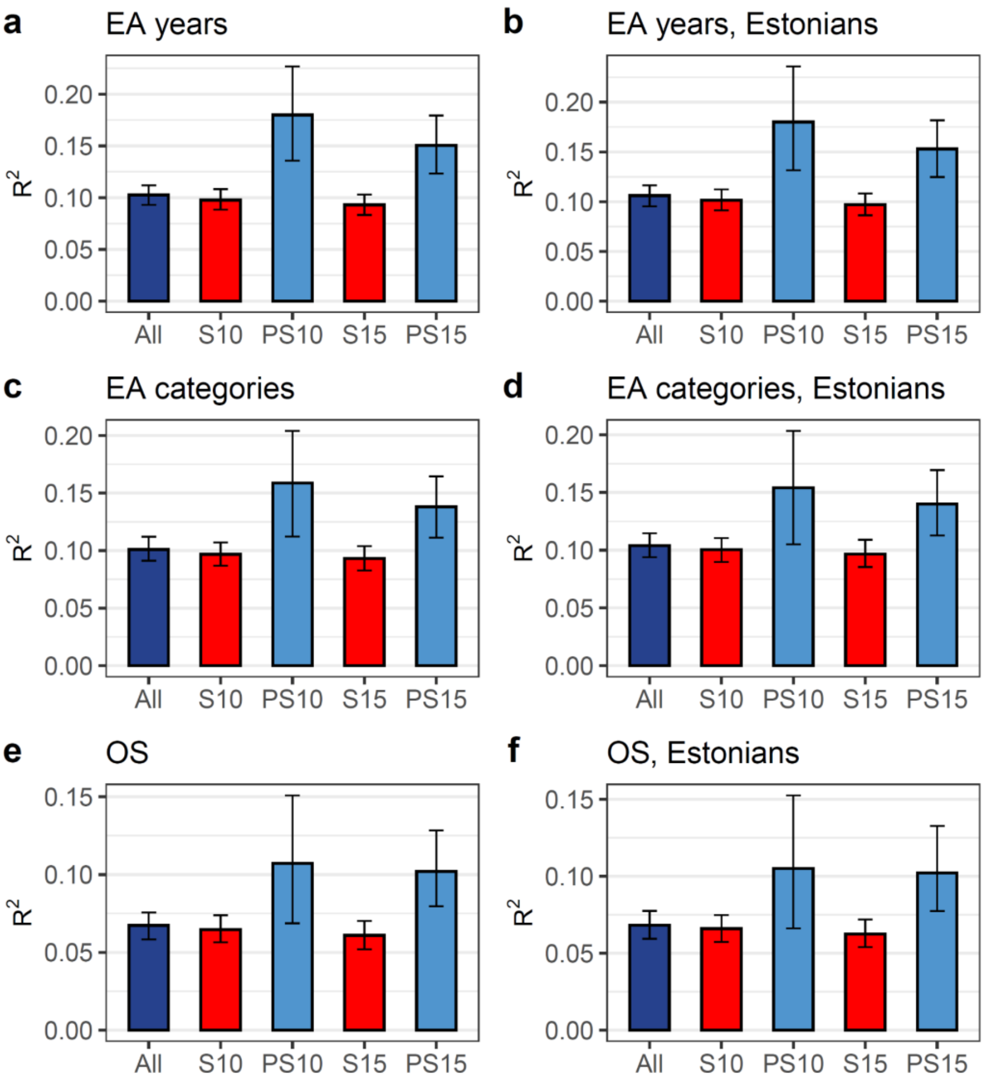

**Supplementary Fig. 34. Variance explained by PGS ($R^2$ for pre-adjusted traits) in the post-Soviet and Soviet eras in the original sample.** S - Soviet, PS - post-Soviet eras by era; index 10 or 15 means cutoff between the eras - the age of individuals in 1991. "EA categories" refer to education as a discrete variable (ordinal EA), while "EA years" indicate that the trait was modelled as years of education. "Estonians" in the title refers to individuals who self-identified as Estonian and were the only ones included in the analysis. Error bars correspond to 95% CI.

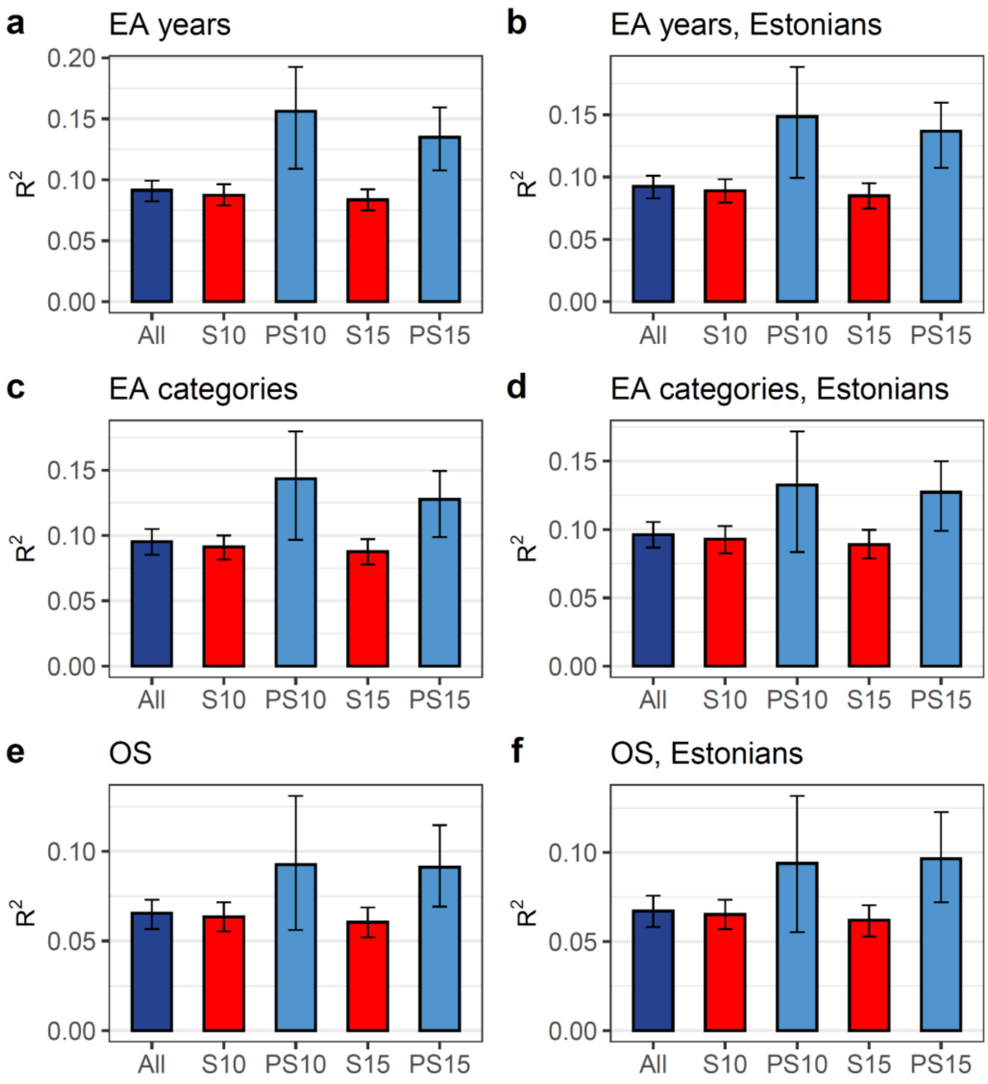

**Supplementary Fig. 35. Variance explained by EA PGS (incremental $R^2$) in the Soviet and post-Soviet eras in the original sample.** S - Soviet, PS - post-Soviet eras by era; index 10 or 15 means cutoff between the eras - the age of individuals in 1991. “EA categories” refer to education as a discrete variable (ordinal EA), while “EA years” indicate that the trait was modelled as years of education. “Estonians” in the title refers to individuals who self-identified as Estonian and were the only ones included in the analysis. EA PGS was calculated with SBayesR based on the EA4 summary statistics with 23andMe and EstBB participants excluded from the meta-analysis. Error bars correspond to 95% CI.

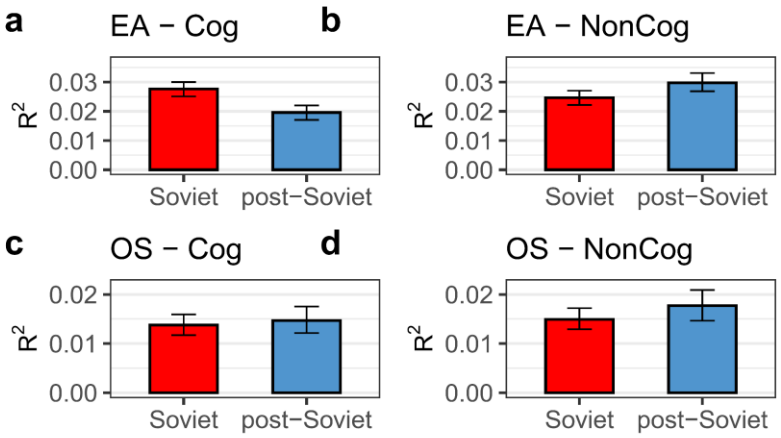

**Supplementary Fig. 36. Educational attainment and occupational status variance explained by $PGS_{Cog}$ and $PGS_{NonCog}$ in the Soviet and post-Soviet eras.** Incremental $R^2$ was calculated in samples divided by era, using a 15-year cutoff. Error bars correspond to 95% CI.

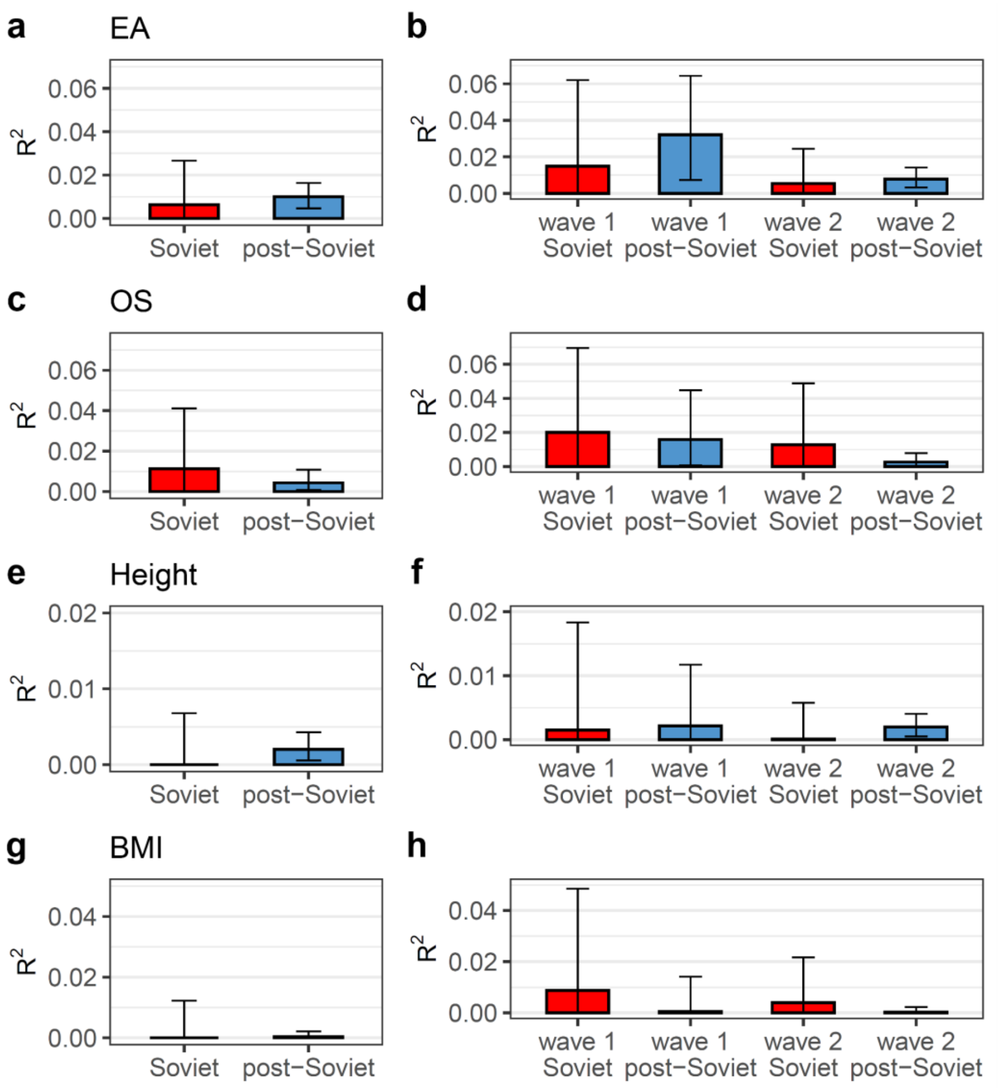

**Supplementary Fig. 37. Educational attainment and occupational status variance explained by PGS based on non-transmitted alleles in the Soviet and post-Soviet eras.** Incremental $R^2$ is shown in groups divided by era (in the left panels), or by era and wave of biobank enrolment (in the right panels) for educational attainment (EA) (**a, b,** respectively), occupational status (OS) (**c** and **d**, respectively), height (**e** and **f**, respectively), and body mass index (BMI) (**g** and **h**, respectively). Error bars correspond to 95% CI.

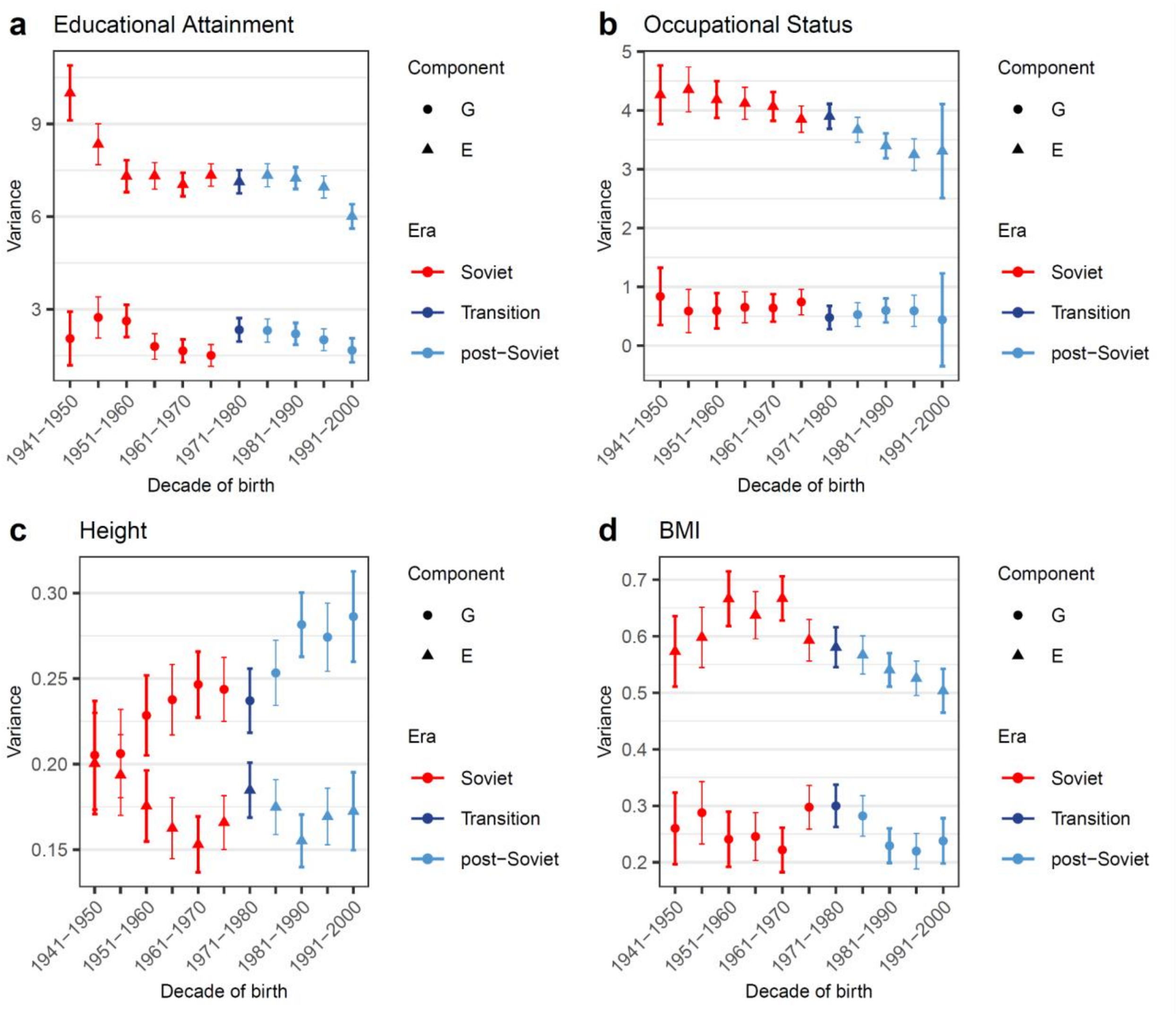

**Supplementary Fig. 38. Absolute genetic (G) variance explained by the genetic relatedness matrix in the stringent-threshold model and residual (E) variance estimated in decade-long birth cohorts with a 5-year overlap between adjacent cohorts.** Those subcohorts belonging to the Soviet era are coloured in red, and the post-Soviet era in light blue. A transition group comprising individuals from both the Soviet and post-Soviet eras is shaded in dark blue. Error bars correspond to 95% CI.
